# Robust axis formation requires both short-range adhesion and long-range attraction

**DOI:** 10.1101/2025.09.27.678924

**Authors:** Guoye Guan, Suxuan Wang, T. Glenn Shields, Seong Ho Pahng, Claire Xinyu Shao, Juns Ye, Christoph Budjan, Sahand Hormoz

## Abstract

Establishing the anterior-posterior (A-P) axis is a key symmetry-breaking event in mammalian development. Gastruloids, aggregates of embryonic stem cells, undergo a similar process, forming a single posterior pole from an initially spherical aggregate. This requires the aggregate to convert an inside-outside difference in cell state into one A-P axis. We asked what cell-cell interactions can make this transition robust. We built an agent-based coarse-grained model with an outer cell population and an inner cell population. Cells interacted through short-range adhesion and, in some simulations, through an effective long-range attraction. Adhesion alone rarely produced a single stable axis, often leaving weakly separated or unstable clusters. Adding long-range attraction between outer cells changed this behavior. The outer cells collected into one pole, the inner cells remained in the opposite domain, and most cells stayed in one aggregate. This occurred across a broad range of adhesion strengths. The model also made a simple prediction. If two polarized gastruloids merge in opposite orientations, the two posterior poles should move around the fused aggregate and join into one pole. Human gastruloid merging experiments showed this behavior. Same-orientation pairs fused rapidly. Opposite-orientation pairs fused more slowly as their posterior poles moved around the aggregate and converged. Finally, we built a minimal gene-regulatory model that generated inner and outer states and turned on the same mechanical interactions. In this model, an initially uniform cell population formed one polarized axis. Together, these results suggest that short-range adhesion and long-range attraction can work together to convert radial patterning into one body axis. They also suggest a design principle for synthetic developmental biology: adhesion can sort cells locally, but building one reproducible axis may require interactions that act beyond direct cell-cell contact.

## INTRODUCTION

The anterior-posterior (A-P) axis is the main body axis of bilaterally symmetric animals. Forming this axis is a key symmetry-breaking event in mammalian development. During gastrulation, the primitive streak forms on the posterior side of the embryo. Cells then move through this region and give rise to mesoderm and endoderm. Gastruloids, three-dimensional aggregates of mouse or human pluripotent stem cells, provide a simple in vitro system to study this transition. They begin as roughly spherical clusters. They then elongate and form one posterior primitive-streak-like domain [Beccari et al. 2018; Moris et al. 2020]. They do this without extraembryonic tissues and without a spatially localized external cue. Gastruloids therefore isolate a self-organizing part of A-P patterning. They let us ask how local cell states and cell-cell interactions produce one global body axis.

Recent work suggests that gastruloids do not form the A-P axis from a uniform state. They first form an inside-outside difference in cell state of spherical aggregates. Outer cells exhibit higher Wnt activity and tend to acquire primitive-streak-like fates. Inner cells show higher Nodal activity and remain more pluripotent [Suppinger et al. *Cell Stem Cell* 2023; McNamara et al. *Nat. Cell Biol*. 2024; Dias et al. *bioRxiv* 2025]. This radial pattern precedes the formation of the A-P axis. It has no unique direction on the surface of the aggregate. To form a single body axis, the aggregate must convert radial patterning into one posterior pole and one opposite anterior pole.

Adhesion is a natural candidate for this conversion. Cadherin expression and other adhesion-related genes change during gastruloid polarization [Mayran et al. *Cell Rep*. 2025; McNamara et al. *Nat. Cell Biol*. 2024]. Adhesion can sort cells with different identities. But local sorting does not guarantee one global axis. Synthetic assemblies built with adhesion or local signaling can form multiple clusters or unstable structures rather than one pole [Toda et al. *Science* 2018; Stevens et al. *Nature* 2023; Yamada et al. *Cell* 2025]. This raises the central question of this study: what cell-cell interactions make a radially patterned aggregate form one stable axis?

We addressed this question with a coarse-grained agent-based model. We represented each cell as a point agent in an overdamped medium. We started with two radially arranged cell populations: an outer shell and an inner core. Cells interacted through short-range adhesion-like forces, and, in a subset of conditions, through an effective long-range attraction or repulsion. This long-range term is phenomenological. It represents any process that biases cells toward or away from other cells over distances longer than direct cell-cell contact. We scanned the interaction strengths and asked which architectures produced one stable axis while keeping most cells in one aggregate.

Differential adhesion alone rarely produced this outcome. It often left weak separation, unstable clusters, or fragmentation. Adding long-range attraction between outer cells changed the behavior. Outer cells collected into one pole. Inner cells remained in the opposite domain. Most cells stayed in one aggregate. This occurred across a broad range of adhesion strengths. Thus, in the model, long-range attraction among outer cells converts radial patterning into a single axis without requiring fine-tuned adhesion.

The model also makes a direct prediction for merging gastruloids. If two polarized gastruloids merge in the same orientation, their axes should join rapidly. If they merge in opposite orientations, their posterior poles should move around the fused aggregate and converge into one pole. Human gastruloid merging experiments showed this behavior. Same-orientation pairs fused rapidly. Opposite-orientation pairs fused more slowly as their posterior domains moved around the aggregate and joined into one single pole. This result supports the model and links the inferred long-range interaction to an experimentally observed collective movement.

Finally, we asked whether gene regulation could generate the cell states and mechanical interactions used in the model. We built a minimal gene-regulatory model with a timer, a transient bifurcation module, and a lock-in module. The network generated inner and outer states from an initially uniform population. It then coupled these states to adhesion and long-range attraction. In this model, a uniform aggregate formed one polarized axis. We implemented this framework in *DevSim*, a simulation platform for coupled genetic and mechanical rules. Together, these results suggest that short-range adhesion and long-range attraction can work together to convert radial patterning into one body axis. This has two implications. First, local adhesion may be insufficient to establish a single reproducible axis, and that robust patterning may depend on mechanisms that coordinate cells across the aggregate. Second, synthetic developmental systems that aim to build reproducible multicellular forms may need more than programmable adhesion. They may also need engineered signaling, motility, or attraction-like interactions that act over longer distances.

## RESULTS

### Human gastruloids reproducibly form a single A-P axis in suspension culture

To first establish the experimental basis for our theoretical work, we asked whether 3D human gastruloids reproducibly undergo axis formation under defined culture conditions. To this end, we applied a recently established protocol for generating gastruloids from human embryonic stem cells (hESCs) [Moris et al. *Nature* 2020a; Moris et al. *Res. Sq*. 2020b]. Approximately 400 cells were aggregated in round-bottom wells and stimulated with the Wnt agonist CHIR99021 (Chir) to induce gastruloid formation (Figure 1A). While control aggregates without Wnt activation remained spherical, Chir stimulated aggregates reproducibly elongated across and within independent experimental batches within 72 hours (Figure 1B; Figure S1; Movie S1).

**Figure 1.**
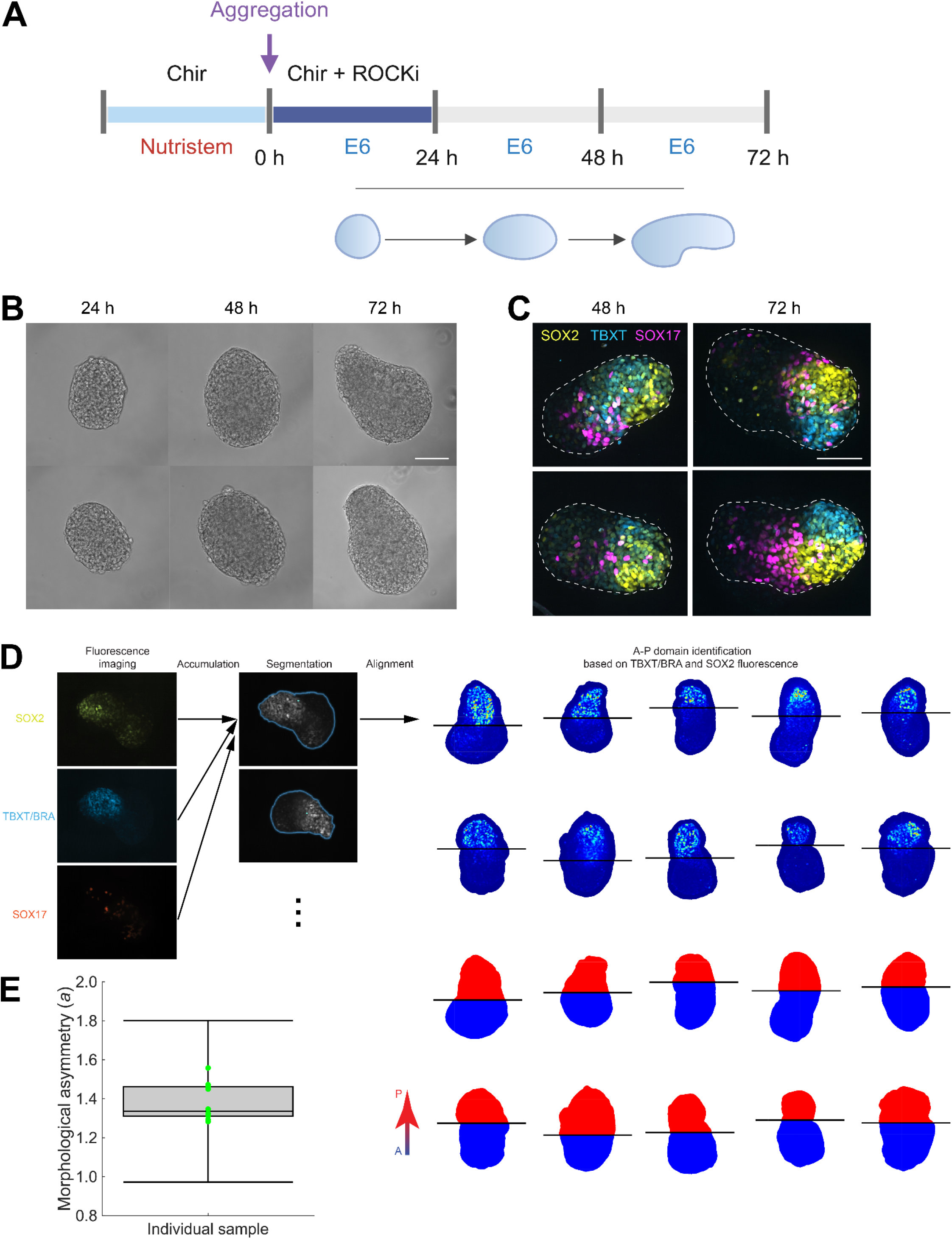
Human stem cell-derived 3D gastruloids exhibit robust A-P axis formation. (A)Schematic of the human gastruloid protocol. hESCs were pretreated with CHIR99021 for 24 h. Cells were then dissociated, and approximately 400 cells were aggregated in each 96-well U-bottom well with CHIR99021 and ROCK inhibitor. After 24 h, aggregates were cultured in E6 (basal) medium without additional patterning factors until 72 h. Chir, CHIR99021; ROCKi, ROCK inhibitor. (B)Representative brightfield images of two gastruloids imaged every 24 h. Chir-treated aggregates elongated by 72 h forming an A-P axis. Scale bar, 100 µm. (C)Representative fluorescence images of two gastruloids at 48 and 72 h made from RUES2-GLR germ layer reporter hES cells. SOX2 labels neuroectoderm-like cells, TBXT/BRA labels primitive-streak/mesoderm-like cells, and SOX17 labels endoderm-like cells. Cells of the three germ layers form spatially organized domains. Scale bar, 100 µm. (D)Quantification workflow. Fluorescence images were combined to segment each aggregate. The aggregate was aligned along its long axis. Anterior and posterior domains were assigned from the SOX2 and TBXT/BRA fluorescence patterns, demonstrating their polarized separation. (E)A-P separation score, a, across 10 gastruloids. The score was computed from binary masks, with each pixel weighted equally. For each gastruloid, a is the distance between the centroids of the posterior and anterior domains divided by the average distance of all pixels in the aggregate mask from the aggregate centroid. Green points indicate individual gastruloids. Box, interquartile range; center line, median; whiskers, 1.5× interquartile range.

At the molecular level, symmetry breaking was also evident in the germ layer organization of gastruloids. We generated human gastruloids using a previously characterized hES germ layer reporter cell line, RUES2-GLR, which labels cells expressing markers of mesoderm (TBXT/BRA), neuroectoderm (SOX2), and endoderm (SOX17). While SOX17^+^ endodermal cells were initially more dispersed, TBXT/BRA^+^ and SOX2^+^ cells rapidly co-localized as a single cluster at the extending distal pole, marking the posterior (Figure 1C). By 72 hours, distinct domains of all three germ layers were consistently found adjacent to one another at this posterior pole. We segmented the TBXT/BRA^+^ and SOX2^+^ domains from fluorescence images using *Segment Anything* (Figure 1D) [Kirillov et al. *ICCV* 2023]. We then computed an image-based A-P separation score, *a*. This score treats the segmented masks as uniform-density regions. It does not weight pixels by fluorescence intensity. For each gastruloid, we calculated the centroid of the posterior domain, the centroid of the opposite anterior domain, and the centroid of the whole aggregate. We defined *a* as the distance between the posterior and anterior centroids divided by the average distance of all pixels in the aggregate mask from the aggregate centroid. Thus, *a* is near zero when the two domains are mixed or concentric, and larger when they occupy opposite sides of the aggregate. Across 10 gastruloids, *a* = 1. 38±0. 10, showing that the domains were reproducibly separated (Figure 1E). To interpret this value, we computed *a* for idealized two-domain geometries with different domain sizes and centroid distances (Figure S2). This comparison shows that the measured value corresponds to a polarized geometry in which one domain is displaced from the other, rather than to mixed or concentric domains. This stereotypical polarized geometry provides a reproducible experimental reference for the modeling below, and stands in sharp contrast to the variable and uncontrolled cluster numbers reported for synthetic organoids that rely solely on differential adhesion [Toda et al. *Science* 2018; Yamada et al. *Cell* 2025].

### A two-population model tests which interactions convert radial patterning into one axis

Mouse and human gastruloids form an A-P axis without extraembryonic tissues [van den Brink et al. *Development* 2014; Turner et al. *Development* 2017; Beccari et al. *Nature* 2018; Moris et al. *Nature* 2020]. Recent work suggests that this axis does not form directly from a uniform aggregate. Instead, gastruloids first establish an inside-outside difference in cell state. In mouse gastruloids, core cells remain more pluripotent, whereas peripheral cells acquire primitive-streak-like features. These populations later break radial symmetry and elongate [Suppinger et al. *Cell Stem Cell* 2023; Anlas et al. Development 2024]. Signal-recording and perturbation experiments further show that Wnt and Nodal activities are organized during this transition and help determine the final A-P pattern [Turner et al. *Development* 2017; McNamara et al. *Nat. Cell Biol*. 2024; Dias et al. *bioRxiv* 2025]. These studies motivate a reduced mechanical problem: given two radially arranged populations, what cell-cell interactions convert that radial pattern into one stable axis?

To answer this question, we built a coarse-grained agent-based model (Figure 2). Each cell was represented as a point agent in an overdamped medium [Fickentscher et al. *Phys. Rev. Lett*. 2016; Nissen et al. *PLoS Biol*. 2017]. This abstraction allowed us to simulate aggregates with cell numbers similar to human gastruloids during axis formation. We initialized 1,500 cells in a compact spherical aggregate. We assigned 25% of the cells to an outer population (marked as “o”) and the remaining 75% to an inner population (marked as “i”), based on the relative domain sizes observed in human gastruloids (Figure 1D). The outer cells formed a peripheral shell. The inner cells formed the core. Thus, the initial condition had radial patterning but no preferred A-P direction.

**Figure 2.**
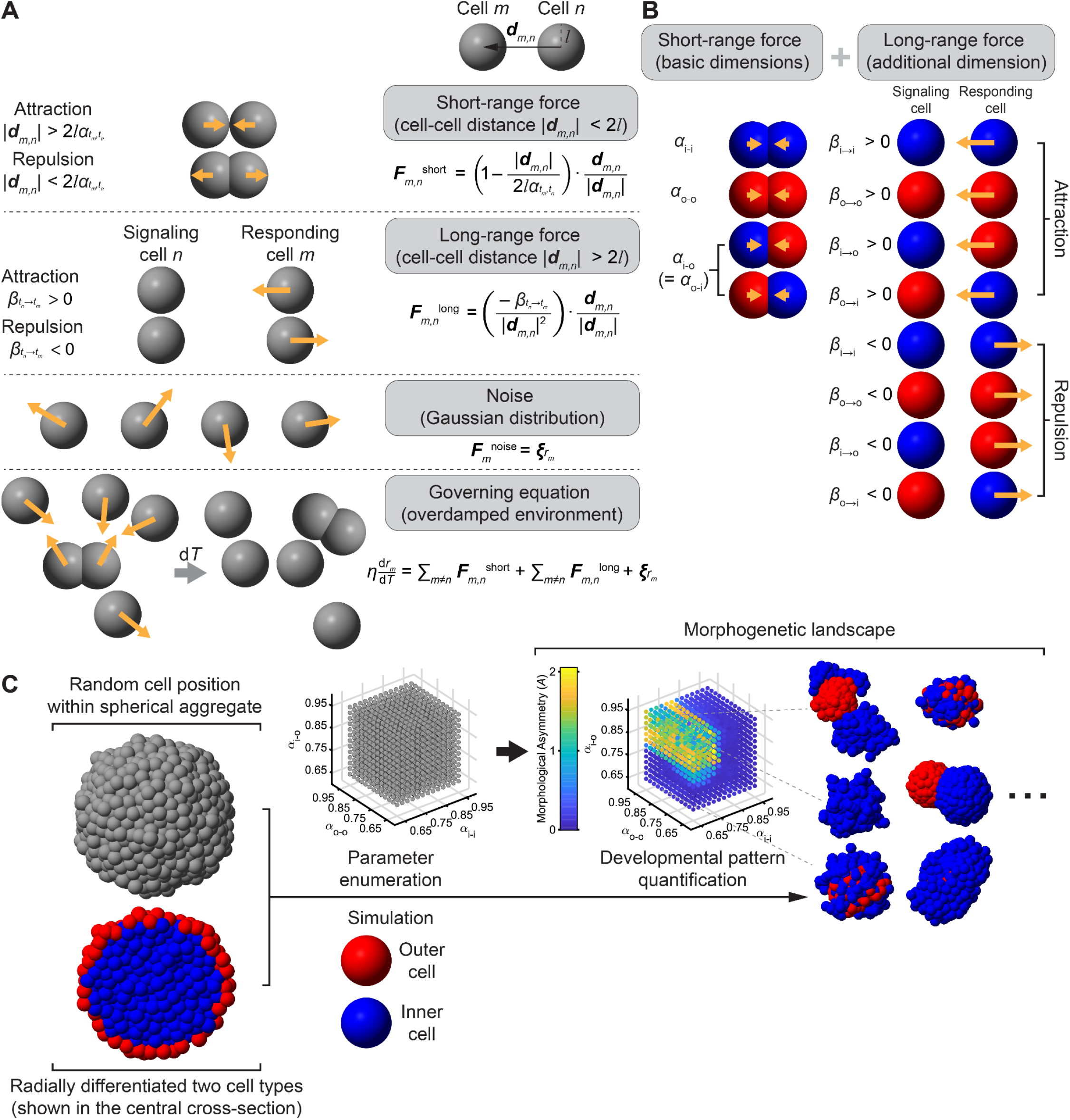
A two-population agent-based model for testing how radial patterning becomes one axis. Each cell is represented as a point agent in an overdamped medium. Short-range contact forces capture effective adhesion between neighboring cells. These forces depend only on the types of the two cells: inner (i) or outer (o). The three short-range parameters are α _i−i_, α _o−o_, and α _i−o_ = α _o−i_. The parameter α sets the distance at which the net contact force is zero; smaller α values correspond to stronger effective adhesion. Beyond the contact range, an optional long-range, chemotaxis-like force acts with strength 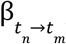, which depends on the signaling cell’s type (t _n_) and the responding cell’s type (t _m_); β > 0 denotes attraction, repulsion, and β = 0 denotes no long-range interaction. Gaussian positional noise is added at each time step. (B)Interaction channels. Short-range interactions define the baseline model through three symmetric adhesion terms: α_i−i_, α_o−o_, and the cross-type term α _i−o_ = α _o−i_. Long-range interactions add four additional channels: inner→inner (β _i_→_i_), outer→outer (β _o_→_o_), inner→outer (β _i_→_o_), and outer→inner (β _o_→_i_). The upper rows show attraction. The lower rows show repulsion. Red cells are outer cells, and blue cells are inner cells. (C)Parameter search and readouts. Simulations start from a compact spherical aggregate with a radial prepattern. Outer cells form a peripheral shell, and inner cells form the core (Figure S3). For each model architecture, we scan the short-range adhesion parameters and, when present, the long-range force parameters. We then quantify the final pattern with two readouts. The asymmetry score, A (computed considering non-uniform density) measures separation of the two cell populations while penalizing fragmentation. The cell loss score, L, measures the fraction of cells excluded from the largest connected aggregate. Representative final configurations show the patterns produced by different parameter choices.

Cells interacted through two classes of forces (Figure 2A). The first was a short-range contact force [Petridou et al. *Cell* 2021]. This force captured volume exclusion and adhesion between neighboring cells. It depended only on the types of the two interacting cells. We used three short-range parameters: α_*i*−*i*_, α_*o*−*o*_, and α_*i*−*o*_. In this force law, α sets the distance at which the net force between two cells is zero. Smaller α values correspond to stronger adhesion, because cells settle closer together. This contact term was motivated by previous models of E-cadherin-mediated cell arrangement and by evidence that cadherins contribute to gastruloid polarization [Yamamoto et al. *Development* 2017; Guan et al. *Commun. Nonlinear Sci. Numer. Simul*. 2022; Mayran et al. *bioRxiv* 2023; McNamara et al. *Nat. Cell Biol*. 2024].

The second class of force was an optional long-range interaction [Pani et al. *eLife* 2018]. This term is phenomenological. It represents any process that biases one cell type to move toward or away from another cell type over distances longer than direct contact. We parameterized this interaction by β. Positive β values represent attraction. Negative β values represent repulsion. A value of zero means that the long-range interaction is absent. Because the interaction can depend on both the signaling cell type and the responding cell type, there are four possible long-range channels: β _*i*_→_*i*_, β _*o*_→_*o*_, β _*i*_→_*o*_, and β _*o*_→_*i*_ (Figure 2B).

We then searched the space of possible interaction rules. For each model architecture, we scanned the short-range adhesion parameters and, when present, the long-range force parameters. We measured whether the aggregate formed one stable axis while keeping most cells in one connected structure. We used two readouts. The first was the simulation asymmetry score, *A*. It is the cell-based version of the image score *a* used in Figure 1: it measures the distance between the centers of the two populations, normalized by aggregate size. We first identified the largest connected aggregate. We then multiplied the normalized center separation by the fraction of inner cells and the fraction of outer cells that remained in that aggregate: 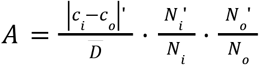. Here, *c*_*i*_ and *c*_*o*_ are the centers of the inner and outer populations in the largest aggregate, 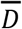 is the average distance of cells from the aggregate center, and 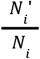 and 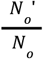 are the retained fractions of each cell type. Thus, a simulation received a large *A* only if the two populations separated and both populations stayed in one connected structure. The second readout was cell loss, *L*, which measured the fraction of cells excluded from the largest connected aggregate. Together, *A* and *L* distinguished productive axis formation from fragmentation. We first tested the simplest model, with short-range adhesion alone. We then added one long-range interaction channel at a time to ask which interaction, if any, made axis formation reliable (Figure 2C).

### Differential adhesion is not sufficient for robust axis formation

We first asked whether short-range adhesion alone could convert radial patterning into one A-P axis. For each adhesion setting, we ran five simulations with independent noise seeds. At the final time point, we identified the largest connected aggregate. We then calculated *A* and *L* for each run and averaged these values across runs. Cells outside the largest aggregate increased *L* and reduced A through the retained-cell penalty defined above.

We scanned the three adhesion parameters: α_*i*−*i*_, α_*o*−*o*_, and α_*i*−*o*_ (Figure 3A-C; Table S1). Smaller α values correspond to stronger adhesion. The top-scoring parameter sets followed the expected logic for cell sorting: inner-inner and outer-outer adhesion were stronger than inner-outer adhesion (Figure 3B). However, this was not enough to produce a single stable axis. Many simulations showed weak separation, local clustering, or fragmentation rather than one global A-P axis (Figure 3C; Figure S4; Movie S2).

**Figure 3.**
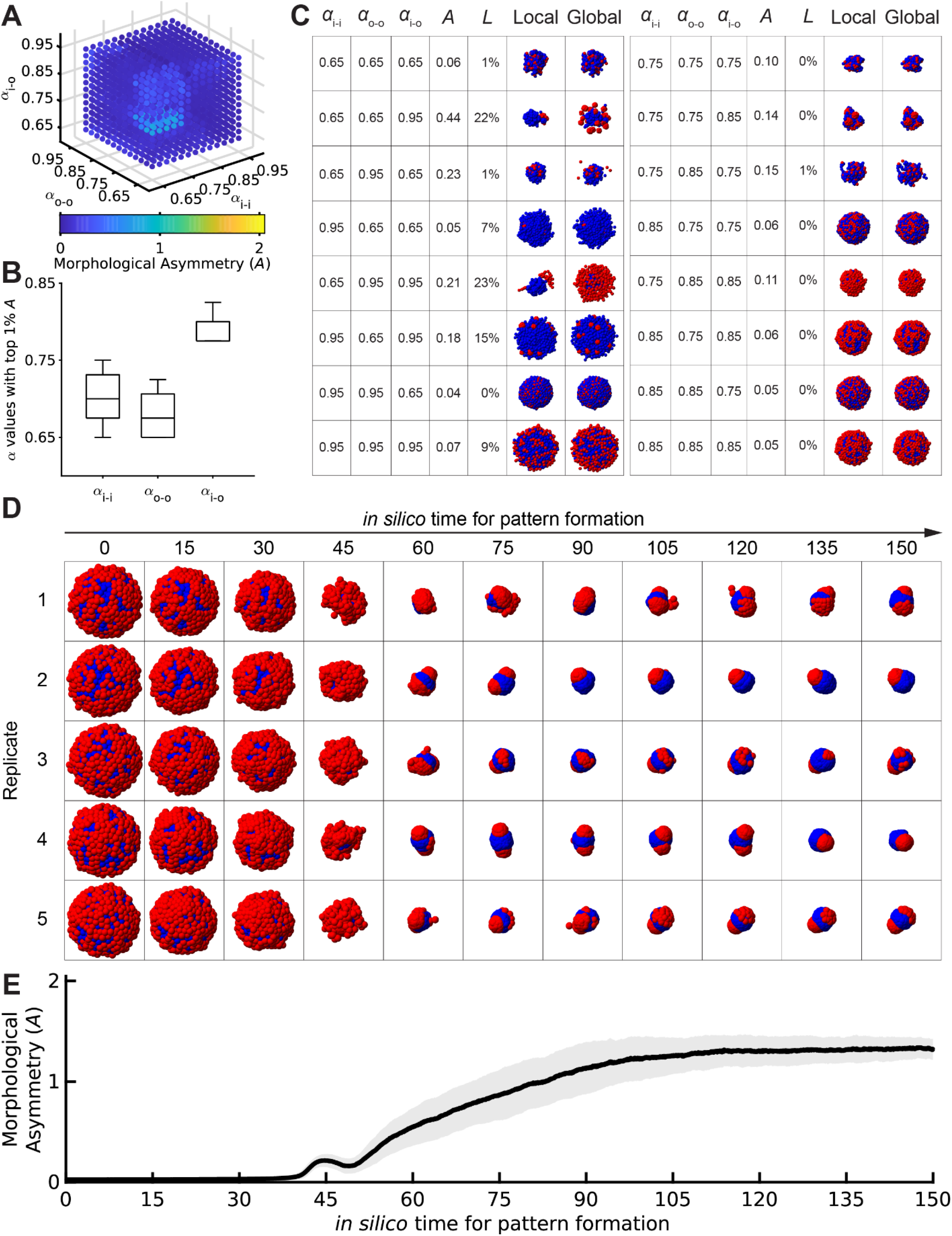
Short-range adhesion alone does not reliably convert radial patterning into one axis. (A)Morphological asymmetry score, A, across the three short-range adhesion parameters α _i−i_, α _o−o_, and α _i−o_. Each point is one adhesion-only parameter setting. Blue indicates low A. Yellow indicates high A. Smaller α values correspond to stronger adhesion. (B)Distribution of α _i−i_, α _o−o_, and α _i−o_ among the top 1% of adhesion-only simulations ranked by A. The top-scoring simulations tend to have stronger same-type adhesion than cross-type adhesion. Box, interquartile range; center line, median; whiskers, 1.5× interquartile range. (C)Representative final states from selected adhesion-only simulations. The table shows the three adhesion parameters, the morphological asymmetry score A, and the cell loss score L. A measures separation of the inner and outer populations while penalizing fragmentation. L measures the fraction of cells outside the largest connected aggregate. “Local” shows the largest connected aggregate. “Global” shows all cells in the simulation. Red cells are outer cells, and blue cells are inner cells. (D)Time course of five representative simulations using the best adhesion-only parameter setting from the scan: (α _i−i_, α _o−o_ α _i−o_) = (0. 750, 0. 725, 0. 800). Even under this setting, the final structures remain variable and do not consistently form one stable axis. (E)Morphological asymmetry score A over time for 100 independent simulations using the same adhesion-only parameter setting as in (D). Black line, mean. Gray shading, standard deviation.

We then examined the best adhesion-only parameter set in more detail. Across 100 independent simulations, the aggregate often began to separate, but the final structures remained variable. Some runs formed partial separation. Others produced small displaced clusters or unstable geometries (Figure 3D-E; Figure S4). Thus, even when adhesion parameters were chosen from the best region of the scan, differential adhesion did not reliably produce the one-axis geometry seen in human gastruloids.

We also varied the magnitude of stochastic noise (Figure S5). At low noise, cells remained trapped near the initial radial configuration. At high noise, adhesion could not keep the aggregate intact. Intermediate noise allowed more rearrangement, but it did not create a broad regime of reliable axis formation. Thus, the failure of the adhesion-only model was not caused by one arbitrary noise choice.

These results are consistent with synthetic multicellular systems in which local adhesion or local cell-cell programs can sort cells but often produce variable cluster numbers rather than one global axis [Toda et al. *Science* 2018; Yamada et al. *Cell* 2025]. They are also consistent with previous computational studies showing that adhesion-driven sorting depends strongly on model details and parameter choice [Osborne et al. *PLoS Comput. Biol*. 2017; Kuang et al. *npj Syst. Biol. Appl*. 2023]. Together, these simulations show that differential adhesion can help cells sort, but it does not by itself make axis formation reliable.

### Long-range attraction between outer cells produces one stable axis

We next asked whether one long-range interaction could rescue the adhesion-only model. We tested eight possibilities: each of the four directed channels, β_*i*_→_*i*_, β_*o*_→_*o*_, β_*i*_→_*o*_, and β_*o*_→_*i*_, as either attraction or repulsion. In each scan, we set one long-range term to a nonzero value and set all other β terms to zero. We then scanned the full grid of adhesion parameters, α_*i*−*i*_, α_*o*−*o*_, and α_*i*−*o*_. Across these scans, we simulated 177,957 parameter settings (Figures S6-S13; Table S1).

Some long-range interactions improved axis formation, but one interaction stood out. Long-range attraction between outer cells, β_*o*_→_*o*_ > 0, dominated the top-scoring solutions (Figure 4A). In these simulations, outer cells collected into one pole. Inner cells formed the opposite domain. Most cells remained in one connected aggregate (Figure 4B-D; Figure S14; Movies S3-S4). This result was not specific to one simulation setting. The same qualitative behavior persisted when we shortened the interaction range, changed the number or fraction of outer cells, or varied the time step and noise level (Figures S15-S17). The successful simulations reached cell-based asymmetry scores of *A*~2 (Figure 4A-D). This value is not directly comparable to the experimental image-based score *a*, because the two scores weigh the same geometry differently. *A* is computed from cell positions and therefore weights regions by their local cell density. In contrast, *a* treats the segmented anterior and posterior domains as uniform-density masks, as in Figure 1. To compare simulations with experiments, we rescored the simulated final patterns using the same uniform-density definition used for a. Simulations with *A*~2 gave *a*~1.46, close to the experimental value of *a* = 1. 38±0. 10 across 10 gastruloids (Figure 1D-E). We then quantified how β_*o*→*o*_ changed the adhesion landscape. In the full scan, 99.94% of the top 1% parameter sets included outer-to-outer attraction. When β_*o*→*o*_ = 0, no adhesion setting met our criterion for axis formation, defined as *A* > 1 and *L* < 0. 1. As β_*o*→*o*_ increased, the fraction of adhesion settings that met this criterion rose to 18% of the full adhesion parameter space (Figure 4E). The mean and maximum values of *A* also increased and then saturated (Figure 4F). For 58.6% of adhesion settings, adding β_*o*→*o*_ increased *A* relative to adhesion alone across the tested values (Figure 4G).

**Figure 4.**
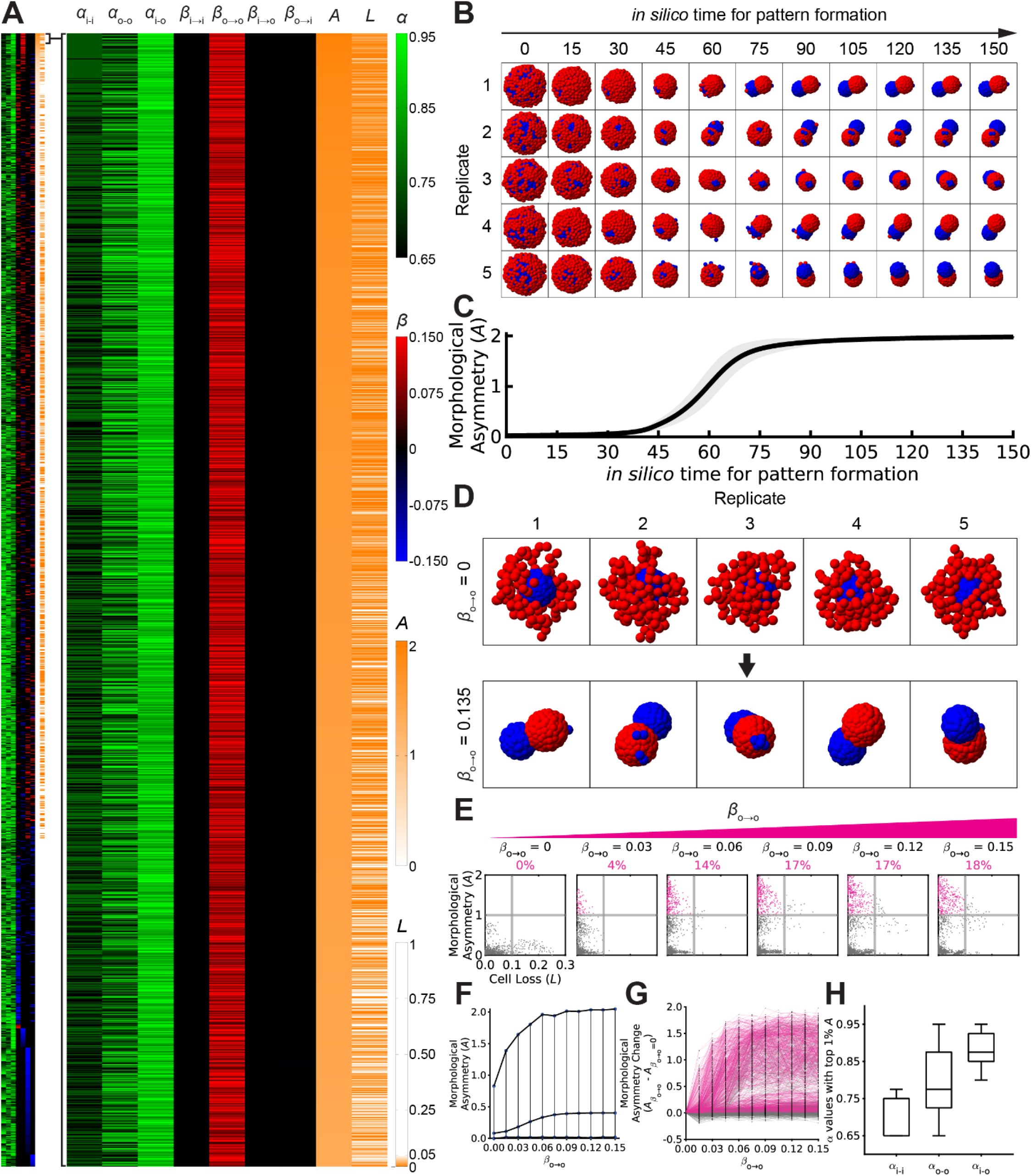
Outer-to-outer long-range attraction expands the parameter region that forms one axis. Parameter map for all 177,957 simulated conditions. Each row is one parameter setting. Rows are ranked by the morphological asymmetry score, A (computed considering non-uniform density). The left map shows the full scan. The right map enlarges the top 1% of conditions ranked by A. Columns show the three short-range adhesion parameters, α_i−i_, α _o−o_, and α _i−o_; the four long-range interaction parameters, β _i→i_, β_o→o_, β_i→o_, and β_o→i_; the asymmetry score, A; and the cell loss score, L. For α, black indicates smaller values and stronger adhesion, and green indicates larger values and weaker adhesion. For β, red indicates attraction, blue indicates repulsion, and black indicates no long-range interaction. For A and L, orange indicates larger values. Outer-to-outer attraction, β_o→o_ > 0, is strongly enriched among the top-scoring simulations. (B)Time course of five representative simulations with outer-to-outer attraction. Simulations used α _i−i_ = 0. 775, α_o−o_ = 0. 95, α_i−o_ = 0. 875, and β_o→o_ = 0. 135, a parameter setting from the top 1% of the scan. Red cells are outer cells. Blue cells are inner cells. (C)Morphological asymmetry score, A, over time for 100 independent simulations using the same parameters as in (B). Black line, mean. Gray shading, standard deviation. (D)Final states for five representative simulations with the same adhesion parameters as in (B). Top row: no outer-to-outer long-range attraction, β_o→o_ = 0. Bottom row: outer-to-outer attraction, β_o→o_ = 0. 135. Adding β_o→o_ converts weak or local separation into one stable axis. (E)Scatter plots of A and L for all adhesion settings at increasing values of β_o→o_. Pink points mark simulations that satisfy the axis-formation criterion, A > 1 and L < 0. 1. The percentage above each plot gives the fraction of adhesion settings that satisfy this criterion. As β_o→o_ increases, the successful region expands from 0% to 18% of the adhesion parameter space. (F)Mean and maximum A across all adhesion settings as a function of β_o→o_. Both increase with β_o→o_ and then saturate. (G)Change in A relative to the adhesion-only case for each adhesion setting. Each line corresponds to one α_i−i_, α_o−o_, α_i−o_ combination. For 58.6% of adhesion settings, adding β_o→o_ increases A across the tested range. (H)Distribution of α_i−i_, α_o−o_, and α_i−o_ among the top 1% of simulations ranked by A. Box, interquartile range; center line, median; whiskers, 1.5 times interquartile range.

These results show that outer-to-outer long-range attraction changes the behavior of the model. Adhesion alone can sort cells, but only in a narrow and unstable region of parameter space. Adding long-range attraction between outer cells creates a broad region in which the model forms one axis while keeping most cells in the aggregate. In the model, this interaction acts as a global coupling among outer cells. It lets the outer population collect into one pole instead of forming multiple local clusters.

### Adhesion provides local cohesion while long-range attraction provides global coordination

The top-scoring simulations showed a consistent adhesion pattern (Figure 4H). Inner-inner adhesion was strong (α_*i*−*i*_ = 0. 72±0. 05). This kept the inner core together. Inner-outer adhesion was weak (α_*i*−*o*_ = 0. 89±0. 04). This allowed the inner and outer populations to separate. Outer-outer adhesion was intermediate (α_*o*−*o*_ = 0. 79±0. 09). This kept outer cells associated, but still allowed them to rearrange along the surface.

These short-range interactions helped sort and stabilize the two populations. But they did not by themselves select one global direction. Outer-to-outer long-range attraction provided this missing global coupling. It biased outer cells toward one another over distances longer than direct cell-cell contact. As a result, the outer population collected into one pole while the inner population remained cohesive on the opposite side.

Adding β_*o*→*o*_ also changed which adhesion values worked. In the adhesion-only model, axis formation was rare and depended on a narrow range of adhesion strengths. With β_*o*→*o*_, a much broader range of α_*i*−*i*_, α_*o*−*o*_, and α_*i*−*o*_ values produced separated domains with low cell loss. In particular, outer-outer adhesion no longer had to be strong. Moderate outer-outer adhesion often worked best, because the outer cells could remain associated while still moving around the aggregate. Thus, outer-to-outer attraction did not replace adhesion. It made axis formation less dependent on fine-tuned adhesion.

### The result does not require a perfect initial outer shell

The simulations above started with outer cells arranged around an inner core. We therefore asked whether axis formation depended on this exact initial geometry. It did not. When half or all outer cells were displaced from a continuous shell, the model still formed one axis (Figure S18). Stronger perturbations had a clearer effect. When inner and outer cell positions were swapped or randomized, axis formation became less reliable and the simulations often produced layered or multi-cluster structures (Figure S19). This failure was rescued when long-range attraction was assigned to the cells positioned at the periphery, even when those cells were originally labeled as inner cells (Figure S20). Thus, a perfect initial shell is not required. What matters is that cells at the periphery can coordinate over distances longer than direct cell-cell contact.

### Long-range attraction predicts posterior-pole fusion in merging gastruloids

The model makes a direct prediction beyond single-gastruloid morphologies. If outer-to-outer long-range attraction helps form one axis, then two separated posterior domains should be able to find each other after two gastruloids merge. This prediction is most clear when two polarized gastruloids meet in opposite orientations. Adhesion alone should leave the two posterior domains at opposite ends of the fused aggregate. Long-range attraction should instead bring the two posterior domains around the surface until they form one pole.

We tested this prediction first in simulations. We placed two polarized model gastruloids side by side, either in the same A-P orientation or in opposite A-P orientations (Figure 5A). The simulated gastruloid shape matched the elongated shape measured in experiments (Figure 1D) [Schmidbauer. *Elem. Math*. 1948; Narushin et al. *Ann. N*.*Y. Acad. Sci*. 2021; Narushin et al. *Biosyst. Eng*. 2022]. When the two gastruloids had the same orientation, their anterior domains and posterior domains merged rapidly, with or without long-range attraction (Movies S5-S6). When the two gastruloids had opposite orientations, the outcome depended on long-range attraction. Without long-range attraction, the anterior domains merged, but the two posterior domains stayed separated at the distal ends of the fused aggregate (Movie S7) [Toda et al. *Science* 2018; Wauford et al. *Cell Systems* 2023; Yamada et al. *Cell* 2025]. With outer-to-outer long-range attraction, the two posterior domains moved along the periphery of the fused aggregate and converged into one domain (Movie S8). Across 100 independent simulations, the fraction of pairs that ended with one continuous posterior domain increased from 0% without long-range attraction to 97% with long-range attraction (Figure S21).

**Figure 5.**
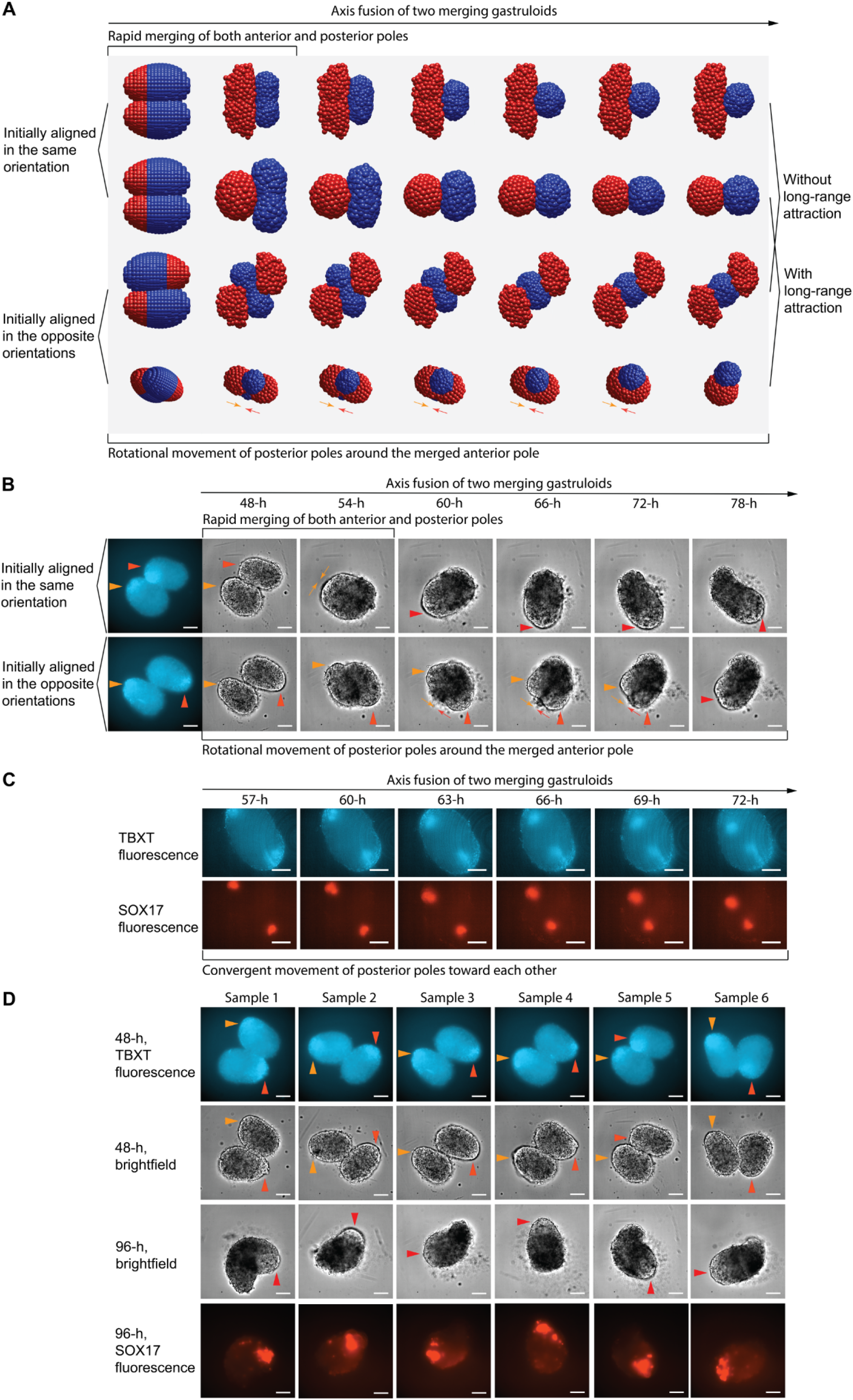
Long-range attraction predicts that merging gastruloids resolve two posterior poles into one. (A)Simulated merging of two polarized gastruloids. Pairs were initialized with either the same A-P orientation or opposite A-P orientations. When the two gastruloids started in the same orientation, the anterior domains and posterior domains merged rapidly. When the two gastruloids started in opposite orientations, the outcome depended on long-range attraction. Without long-range attraction, the two posterior domains remained separated. With long-range attraction, the posterior domains moved around the fused aggregate and converged into one domain. (B)Representative time course imaging of two merging human gastruloids imaged every 6 h. The first column shows TBXT/BRA fluorescence at 48 h, before merging, to mark the initial posterior poles. Subsequent columns show brightfield images during merging. Orange and red triangles mark the two posterior poles. Same-orientation pairs merged rapidly. Opposite-orientation pairs merged more slowly as the posterior poles moved around the periphery of the fused aggregate. Scale bars, 100 µm. (C)Representative fluorescence time course imaging of an opposite-orientation pair during merging. The top row shows TBXT/BRA fluorescence. The bottom row shows SOX17 fluorescence. Both reporters mark posterior-associated domains that move progressively closer to each other as the two gastruloids merge. Scale bars, 100 µm. (D)Six independent merging experiments under the standard CHIR dose. Rows show, from top to bottom: TBXT/BRA fluorescence at 48 h, brightfield at 48 h, brightfield at 96 h, and SOX17 fluorescence at 96 h. TBXT/BRA fluorescence marks the two initial posterior poles before merging. SOX17 fluorescence marks the final posterior-associated domain after merging. In all six samples, the fused aggregate formed one final posterior domain. Six additional samples cultured with reduced CHIR are shown in Figure S22. Scale bars, 100 µm.

We then tested this prediction experimentally, with details given in the METHOD DETAILS - Gastruloid merging experiment. We cultured human gastruloids for 48 h, when they had already formed polarized domains. We then transferred one gastruloid into a U-bottom well that already contained another gastruloid and imaged the pair for another 48 h (Figure 5B). TBXT/BRA fluorescence marked the initial posterior region at 48 h. Pairs that began in the same orientation fused rapidly, usually within 6 h. Pairs that began in opposite orientations fused more slowly, over about 30 h. In these opposite-orientation pairs, the posterior regions moved around the periphery of the fused aggregate and approached each other, matching the simulated trajectory (Figure 5B; Movies S9-S10).

Fluorescence imaging supported the same conclusion. During fusion, SOX17-positive domains became concentrated and moved progressively closer to each other, in concordance with TBXT/BRA fluorescence (Figure 5C). By the end of the experiment, the merged aggregate contained one SOX17-positive domain. This occurred in 6/6 pairs cultured with the standard Chir dose (0.5 µM) and in 6/6 pairs cultured with a reduced Chir dose (0.25 µM) (Figure 5D; Figure S22; Movies S11-S12).

Thus, the model predicted two features of merging gastruloids: orientation-dependent fusion time and peripheral convergence of separated posterior domains. The experiments confirmed both features. These results support the idea that an effective long-range attraction-like interaction helps coordinate posterior domains during axis formation.

### A minimal gene-mechanical circuit can generate radial states and activate axis formation

Until this point, the model assumed that inner and outer cell states already existed. We next asked whether a simple regulatory circuit could generate these states from an initially uniform aggregate and then activate the mechanical interactions needed for axis formation.

We built a three-gene regulatory model. Each simulated cell carried the same three genes, G1, G2, and G3. Gene expression changed through intracellular regulation, distance-dependent intercellular signaling, degradation, leakage, and noise. Mechanical interactions were then read out from the gene state. One readout changed short-range adhesion. A second readout changed long-range attraction. Thus, cells were not assigned fixed inner or outer identities. Their mechanical behavior emerged from their gene-expression state.

The circuit had three modules (Figure 6A-B). G1 acted as a timer. It started high and then decreased. This created an early establishment stage and a later mechanical stage. The timer did not break spatial symmetry by itself. Instead, spatial information came from the aggregate geometry. Each cell received a distance-decaying signal from other cells. Cells in the center received signals from many directions. Cells at the surface received less signal because there were no cells outside the aggregate [Warmflash et al. *Nat. Methods* 2014]. This created an inside-outside difference in G2 expression.

**Figure 6.**
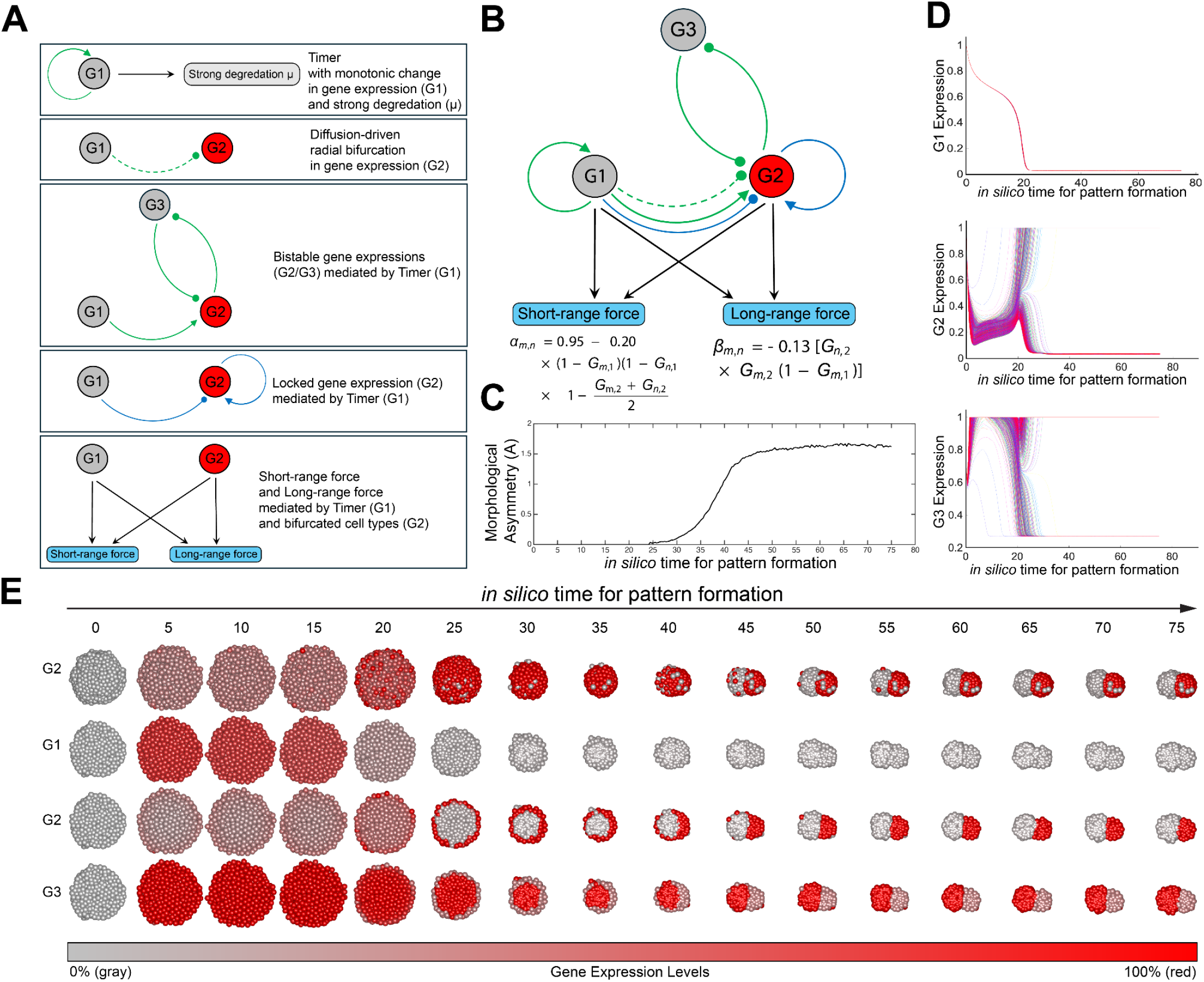
A minimal gene-mechanical circuit generates radial cell states and converts them into one axis. (A)Design logic of the three-gene circuit. G1 acts as a timer. It starts high and then decreases, creating an early establishment stage and a later maintenance stage. During the establishment stage, signaling from G1 creates a radial difference in G2. G2 and G3 then form a mutually inhibitory switch that separates cells into two gene-expression states. During the maintenance stage, a lock-in module preserves these states [Wolpert. J. Theor. Biol. 1969]. The final gene state controls short-range adhesion and long-range attraction. (B)Schematic of the full gene-mechanical network. Each cell contains the same three-gene circuit. Gene regulation includes intracellular interactions and distance-dependent intercellular signaling. Mechanical parameters are read out from gene state. One readout controls the short-range adhesion parameter. A second readout controls the long-range attraction parameter. In this abstract circuit, the G2/G3 switch can be viewed as a Wnt/Nodal-like opposition, but the genes are not assigned to specific molecular pathways. (C)Morphological asymmetry score, A, over time. A increases as the gene circuit activates the mechanical interactions and the aggregate forms one axis. (D)Gene-expression trajectories over time. G1 decreases as the timer turns off. G2 and G3 then bifurcate into two stable expression states. Each trace shows one simulated cell. Colors distinguish cells and do not encode additional variables. (E)Time course of pattern formation from an initially uniform spherical aggregate. Red indicates high gene expression, and gray indicates low gene expression. Row 1 shows the whole aggregate colored by G2 expression. Rows 2–4 show cross-sections colored by G1, G2, and G3 expressions. The circuit first generates radial gene-expression differences. It then activates mechanical interactions that move the outer population along the surface and into one pole. This produces a single polarized axis without pre-assigned cell types or an imposed A-P direction.

G2 and G3 then formed a mutually inhibitory switch [Elowitz and Leibler, *Nature* 2000; Gardner et al. *Nature* 2000]. This switch converted the radial difference in G2 into two stable gene-expression states [Zhu et al. *Nat. Chem. Biol*. 2023; Chen et al. *eLife* 2025]. This G2/G3 switch can be viewed as an abstract version of the Wnt/Nodal opposition observed in gastruloids [Dias et al., *bioRxiv* 2025; Wehmeyer et al. *bioRxiv* 2025; Arias et al. *Cells Dev*. 2025]. We do not assign G2 and G3 to specific molecular pathways. Rather, one state is Wnt-like and the other is Nodal-like. The point is that a mutually inhibitory signaling circuit can convert a graded radial signal into two stable cell states.

As G1 decreased, the gene states began to control mechanics. Cells in one state acquired the attraction-like interaction that coordinates cells over distances longer than direct contact. Cells in the other state remained cohesive and formed the opposite domain. Small angular fluctuations in the initially spherical aggregate then selected a direction. Long-range attraction amplified this fluctuation, causing the peripheral population to move along the surface and collect into one pole. The simulation followed this sequence. G1 decreased over time. G2 and G3 bifurcated into two cell populations. The morphological asymmetry score increased. The outer population moved along the periphery and converged into one pole (Figure 6C-E; Figure S23; Movies S13-S14). Thus, the mechanical design principle identified above can be embedded in a small regulatory circuit. The model does not require pre-assigned cell types or an externally imposed A-P axis, but it does require intercellular signaling, aggregate geometry, and mechanical amplification of fluctuations.

### *DevSim* enables interactive exploration of gene-mechanical patterning models

The three-gene circuit is one example of a larger design space. Other circuits could use different numbers of genes, different signaling ranges, different adhesion rules, or different long-range interactions. To make this design space easier to explore, we built *DevSim*, a *MATLAB*-based platform for simulating gene-mechanical models of multicellular patterning (Figure 7A; Movie S15; Supplemental Text 1) [The MathWorks Inc. 2024].

**Figure 7.**
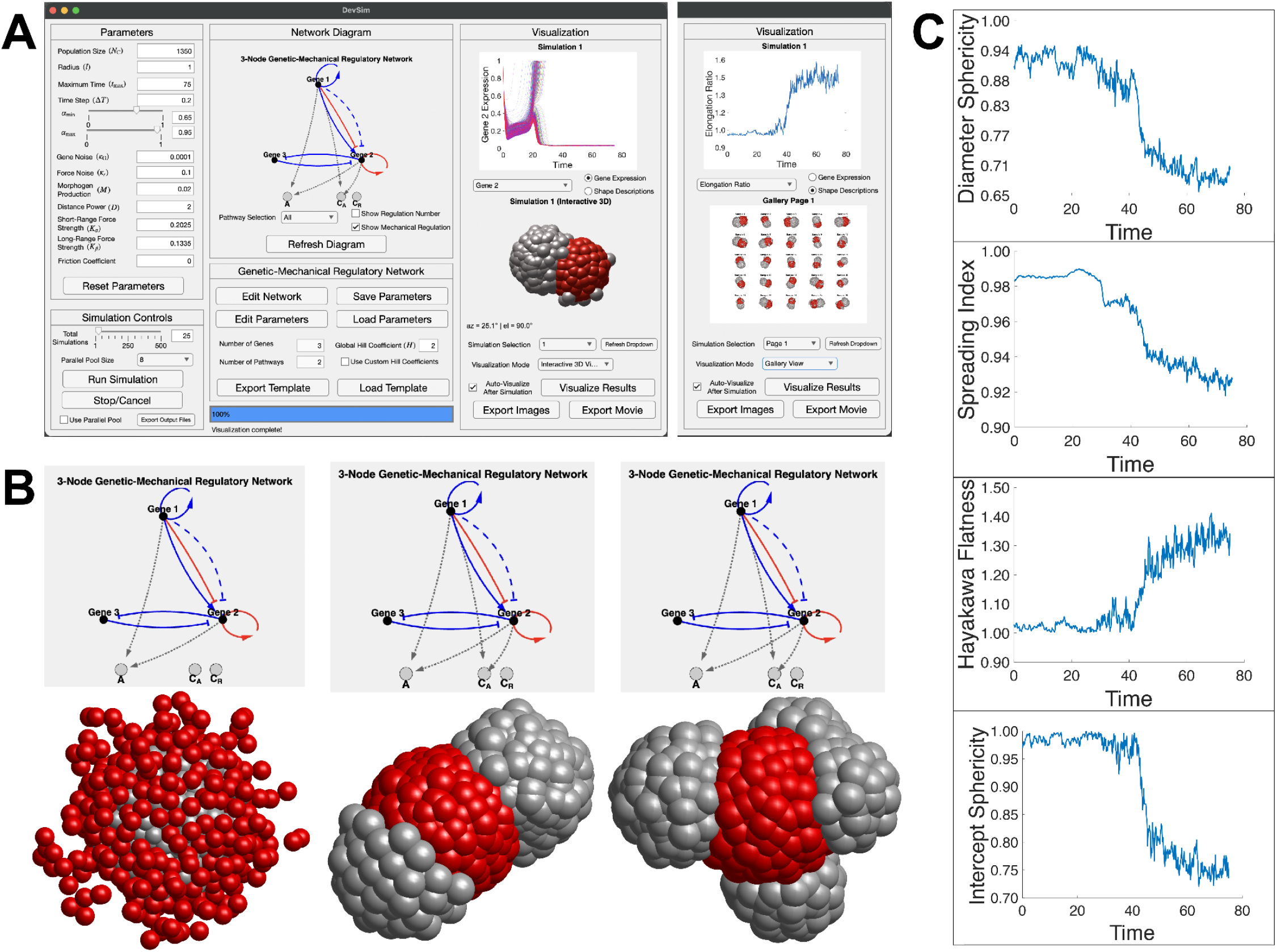
DevSim enables interactive simulation and analysis of gene-mechanical patterning models. (A)DevSim interface for the three-gene model from Figure 6. The interface includes parameter controls, simulation controls, a network diagram, gene-expression plots, shape-dynamics plots, 3D renderings, and a gallery of independent simulations. The left visualization column shows Gene 2 expression over time and a 3D rendering of a single-pole pattern. The right visualization column shows elongation ratio over time and final 3D patterns from 25 independent simulations. A user guide is provided in Supplementary Text 1. (B)Example patterns generated by changing the network topology and parameters in DevSim. Network diagrams are shown above the corresponding final structures. From left to right: layered, bilobed, and multilobed patterns. Corresponding simulations are shown in Movies S16-S18. Cell color indicates simulated gene-expression state. (C)Shape descriptors measured over time from DevSim output. From top to bottom: diameter sphericity, spreading index, Hayakawa flatness, and intercept sphericity (see Supplemental Text 1 for a detailed description). These descriptors quantify how the aggregate changes shape during pattern formation.

*DevSim* implements the same model class used above (see Supplemental Text 1 for a detailed description of the model). Each cell carries a user-defined gene network. Gene expression changes through intracellular regulation, distance-dependent intercellular signaling, degradation, leakage, and noise. Gene state is then read out into mechanical parameters that control short-range adhesion and long-range attraction or repulsion. Users can change the cell population, gene network, signaling parameters, mechanical parameters, and noise levels. Simulations can be run on a personal computer, a computing cluster, or through *MATLAB* Online.

*DevSim* also provides tools for visualizing and analyzing the simulations. It displays the gene network, cell positions, gene-expression patterns, and single-cell gene-expression trajectories. It also measures whole-aggregate shape over time using 3D shape descriptors (Figures S23-S24; Table S2) [Wadell. *J. Geol*. 1932; Wadell. *J. Geol*. 1933; Zingg. *Schweiz. Mineral. Petrogr. Mitt*. 1935; Krumbein. *J. Sediment. Res*. 1941; Sneed et al. *J. Geol*. 1958; Wilson et al. *Earth Planet. Sci. Lett*. 1979; Hayakawa et al. *Comput. Geosci*. 2005; Yu et al. *Adv. Healthc. Mater*. 2013; Guan et al. *Membranes* 2024;]. In the three-gene circuit from Figure 6, *DevSim* tracks the rise in morphological asymmetry and the accompanying changes in aggregate shape (Figure 7A,C).

We then used *DevSim* to test whether the same framework could produce other multicellular patterns. By changing the network topology and parameters, the model generated layered, bilobed, multilobed, and striped patterns (Figure 7B; Movies S16-S19). Thus, *DevSim* provides an executable framework for testing how genetic regulation and mechanical interactions together shape self-organizing multicellular structures.

## DISCUSSION

A central problem in development is how a group of cells forms one body axis. In gastruloids, this problem has two steps. First, cells form an inside-outside difference in state. Outer cells tend to acquire primitive-streak-like features, while inner cells remain more pluripotent. Second, this radial pattern must be converted into one anterior-posterior axis. The second step is not automatic. A radial pattern has no preferred direction on the surface of the aggregate.

We asked which cell-cell interactions can make this conversion reliable. A two-population model showed that short-range adhesion alone rarely formed one stable axis. Adhesion could sort the two populations locally, but it often produced weak separation, unstable structures, or multiple clusters. Adding long-range attraction between outer cells changed the behavior. Outer cells collected into one pole. Inner cells remained in the opposite domain. Most cells stayed in one aggregate. This occurred across a broad range of adhesion strengths.

The merging experiments provided a direct test of this idea. The model predicted that two gastruloids with opposite orientations should behave differently depending on whether long-range attraction is present. Without long-range attraction, the two posterior domains should remain separated. With long-range attraction, the two posterior domains should move around the fused aggregate and converge into one pole. Human gastruloid merging experiments showed the predicted behavior. Same-orientation pairs fused rapidly. Opposite-orientation pairs fused more slowly, as their posterior domains moved around the aggregate and formed one final posterior pole. This does not prove the molecular identity of the long-range interaction. But it supports the idea that an effective attraction-like interaction helps coordinate posterior domains over distances longer than direct cell-cell contact.

This result changes how we think about adhesion in axis formation. Adhesion is clearly important. It can keep cell populations cohesive and help different cell states separate. But adhesion is local. It acts through contact between neighboring cells. Local sorting rules do not necessarily select one global direction. In our model, long-range attraction provides the missing global coupling. It allows cells in the outer population to coordinate across the aggregate and collect into one pole. Thus, short-range adhesion and long-range attraction play different roles. Adhesion provides local cohesion and sorting. Long-range attraction selects one global axis.

The long-range term in the model is phenomenological. It should not be interpreted as a physical force with a known molecular mechanism. It could represent several biological processes. Cells may secrete and respond to diffusible signals. They may migrate directionally through chemotaxis. They may polarize and crawl along the tissue surface. Tissue-scale flows, actomyosin contractility, or signaling-dependent changes in motility could also produce an effective attraction between posterior cells. In gastruloids, Wnt, Nodal, BMP, and FGF signaling are natural candidates to test, because these pathways regulate early patterning and cell-state divergence [McNamara et al. *Nat. Cell Biol*. 2024; Dias et al. *bioRxiv* 2025; Liu et al. *Stem Cell Rep*. 2021; Gattiglio et al. *Biol. Open* 2023].

Several experiments could distinguish these possibilities. First, 3D live imaging could track posterior SOX2^+^ or TBXT/BRA^+^ cells during axis formation and merging [Gros et al. *eLife* 2025]. This would show whether their velocities contain a directional component toward other posterior cells, beyond passive tissue relaxation. Second, perturbing Wnt, Nodal, BMP, FGF, cadherins, or actomyosin during the merging assay could test which pathways are required for posterior-pole convergence [Mayran et al. *Cell Rep*. 2025]. Third, placing polarized gastruloids near each other before physical contact could test whether posterior domains influence each other through secreted signals [Anlaş et al. *bioRxiv* 2021; Anand et al. *Cell* 2023]. These experiments would move the model from an effective interaction toward a molecular mechanism.

The results also have implications for synthetic developmental biology. Many synthetic multicellular systems use engineered adhesion or local cell-cell signaling to build structures [Toda et al. *Science* 2018; Stevens et al. *Nature* 2023; Yamada et al. *Cell* 2025]. These approaches can sort cells and create local organization. But our results suggest that local rules may not be enough to build one reproducible axis. A synthetic aggregate may also need a way for cells to coordinate over longer distances. This could be implemented through engineered morphogen secretion, chemotaxis, motility control, or attraction-like interactions coupled to cell state. In this sense, long-range coordination may be an important design rule for building reproducible multicellular forms.

We also showed that the mechanical design can be embedded in a minimal gene-mechanical circuit. The three-gene model starts from an initially uniform aggregate. G1 acts as a timer. It does not provide spatial information by itself. Instead, it separates an early pattern-establishment stage from a later mechanical stage. Spatial information comes from the geometry of the aggregate: cells near the center and cells near the surface receive different levels of distance-dependent intercellular signal. The G2/G3 switch converts this radial difference into two stable cell states. This switch can be viewed as an abstract version of the opposing Wnt/Nodal activities reported in gastruloids and mammalian embryos [Dias et al. *bioRxiv* 2025; Wehmeyer et al. *bioRxiv* 2025; Arias et al., *Cells Dev*. 2025], although we do not assign G2 and G3 to specific molecular pathways. As G1 decreases, these cell states begin to control adhesion and long-range attraction. Long-range attraction then amplifies small angular fluctuations and converts the radial pattern into one axis. Thus, a regulatory circuit can generate cell states, and those states can control the mechanical interactions that shape the aggregate.

*DevSim* was built to explore this larger design space. The three-gene circuit is only one example. Other circuits could use different numbers of genes, different signaling ranges, different adhesion rules, or different long-range interactions. *DevSim* lets users change these rules and simulate the resulting multicellular patterns. It also measures gene dynamics, cell positions, and aggregate shape over time. This makes it possible to test how genetic regulation and mechanical interactions combine to produce layered, bilobed, multilobed, striped, or polarized structures.

The current model has clear limitations. Cells are represented as point agents. The short-range force approximates adhesion and volume exclusion, but it does not represent cell shape, cell-cell contact area, cortical tension, or tissue rheology explicitly [Kajita et al. *Bioinformatics* 2003; Kuang et al. *PLoS Comput. Biol*. 2022]. The long-range force is instantaneous and pairwise. It does not model diffusion, ligand production, receptor binding, sensing, polarization, or adaptation [Hu et al. *PLoS Comput. Biol*. 2014; Nakamura et al. *Phys. Rev. Res*. 2022; Yaman et al. *Cell* 2023; Al-Hilal et al. *Nat. Cell Biol*. 2025]. Cell number is fixed, although real gastruloids grow and their patterning depends on size [Fiuza et al. *Cells Dev*. 2025; Bennabi et al. *Sci. Adv*. 2025]. Future models should include explicit morphogen fields, cell polarity, cell division, and more realistic tissue mechanics [Nissen et al. *eLife* 2018; Cao et al. *eLife* 2019; Nielsen et al. *iScience* 2020; Tian et al. *Phys. Biol*. 2020]. They should also be calibrated to experimental length, time, and force scales. Despite these limitations, the model identifies a simple principle. Radial patterning does not automatically create one axis. Adhesion can sort cells locally, but reliable axis formation may require a second interaction that coordinates cells across the aggregate. In human gastruloids, the convergence of posterior domains during merging supports this principle. More broadly, the work suggests that self-organizing tissues may use both local adhesion and longer-range coordination to turn cell-state differences into reproducible form.

## STAR★METHODS

Detailed methods are provided in the online version of this paper and include the following:

- KEY RESOURCES TABLE
- RESOURCE AVAILABILITY Lead Contact

Materials availability Data and code availability

- RESOURCE AVAILABILITY
- EXPERIMENTAL MODEL AND SUBJECT DETAILS
- METHOD DETAILS

Confocal fluorescence microscopy Brightfield microscopy

Gastruloid merging experiment Biophysical models

Platform construction

- QUANTIFICATION AND STATISTICAL ANALYSIS

1. Developmental pattern description

## SUPPLEMENTAL INFORMATION

Supplemental Information can be found along this paper.

## ACKNOWLEDGEMENTS

We thank all members of Hormoz Lab for their support, with special gratitude to Senjuti Gayen and Tianhua Zhao for their experimental assistance, and to Yitong Yang, Andrea Perry, Alvaro Tello-Rodriguez, and Audric Adonteng for their computational assistance. We also thank the Core for Imaging Technology & Education at Harvard Medical School for help with light microscopy, with special gratitude to Dr. Jennifer Waters, Dr. Talley Lambert, Dr. Eva de la Serna, Dr. Anna Payne-Tobin Jost, Dr. Federico Gasparoli, and Dr. Asemare Taddesse, for their technical assistance. We are grateful to Prof. Alfonso Martinez Arias for his constructive suggestion on project progress. Thanaphone Glenn Shields is funded by Undergraduate Project & Harvard College Research Program (HCRP); Juns Ye is funded by Continuing Umbrella of Research Experiences Program (CURE). Portions of this research were conducted on the O2 High Performance Compute Cluster supported by the Research Computing Group, at Harvard Medical School; portions of this research were conducted on the High Performance Compute Cluster supported by the Data Science Systems Group, at Dana-Farber Cancer Institute. This work was supported by the National Institutes of Health, National Heart, Lung, and Blood Institute (NIH/NHLBI R01 HL158269) and by the Barry Family HSCI Innovation Award for Early Investigators.

## AUTHOR CONTRIBUTION

Biophysical modeling and genetic-mechanical regulatory network design: G.G.; Parameter space exploration and quantitative statistical analysis: G.G., S.W.; *DevSim* platform construction: G.G., T.G.S.; Experiments: G.G., C.X.S., C.B.; Manuscript writing and revision: G.G., S.W., T.G.S., C.B., S.H.; Supervision: S.H.

## DECLARATION OF INTERESTS

The authors declare no competing interests.

## STAR★METHODS

Detailed methods are provided in the online version of this paper and include the following:

## RESOURCE AVAILABILITY

| REAGENT or RESOURCE | SOURCE | IDENTIFIER |
| --- | --- | --- |
| **Reagents** |  |  |
| NutriStem hPSC XF Medium | Sartorius | Catalog # 05-100-1B |
| Vitronectin (VTN-N) | Thermo Scientific | Catalog # A14700 |
| Essential 6 (E6) Medium | Thermo Scientific | Catalog # A1516401 |
| 96W Cell-Repellent Plate, Round (U) Bottom | Greiner Bio-One | Catalog # 650970 |
| Y-27632 dihydrochloride | Tocris | Catalog # 1254 |
| CHIR99021 | Tocris | Catalog # 4423 |
| **Cell lines** |  |  |
| RUES2-GLR | [Martyn et al. *Nature* 2018] | RUESe002-A-6 (RRID:CVCL_E9CV) |
| H9 SOX17<sup>mCHERRY/w</sup>, RUNX1C<sup>GFP/w</sup> | [Ng et al. *Nat. Biotechnol.* 2016] | Parental cell line: H9 (WA09), (RRID: CVCL_9773) |
| **Software and algorithms** |  |  |
| MATLAB (R2023b, R2024b, R2025b) | The MathWorks | RRID:SCR_001622<br>https://www.mathworks.com/products/matlab.html |

### Lead Contact

Further information and requests for resources and reagents should be directed to and will be fulfilled by the Lead Contact, Sahand Hormoz.

### Materials availability

This study did not generate new unique reagents.

### Data availability

The data is available at https://github.com/hormoz-lab/Symmetry-Breaking.git.

### Code availability

The code is available at https://github.com/hormoz-lab/Symmetry-Breaking.git.

## EXPERIMENTAL MODEL AND SUBJECT DETAILS

Human ES cells were maintained in NutriStem hPSC XF medium (Sartorius, 05-100-1B) on Vitronectin-coated plates (VTN-N; Thermo Fisher Scientific, A14700). To prepare cells for generating gastruloids, 2 × 10^4^ RUES2-GLR hES cells [Martyn et al. *Nature* 2018] were seeded onto Vitronectin-coated 6-well multi-well plates in Nutristem and supplemented with 10 µM Y-27632 dihydrochloride (ROCK inhibitor; Tocris) for the first 24h after seeding. Media changes were performed daily. On Day 4, cells were pre-treated with 3.25 µM Chir (Tocris). On Day 5, cells were harvested and dissociated into single cells. 400 cells were transferred into 96-well U-bottom multi-well plates in 40 µL E6 media supplemented with ROCKi and 0.5 µM Chir. After 24h, 150 µL per well E6 media was added. E6 media was refreshed subsequently every 24h by removing 150 µL media and adding 150 µL fresh E6 media.

## METHOD DETAILS

### Confocal fluorescence microscopy

Live imaging of human gastruloids was performed on a Yokogawa CSU-W1 spinning-disk confocal mounted on a Nikon Ti2 motorized inverted microscope equipped with Perfect Focus System (PFS), controlled by NIS-Elements (Nikon). The system includes a Nikon LUN-F XL solid-state laser combiner (405/445/488/514/561/640 nm), a Hamamatsu ORCA-Fusion BT sCMOS camera, and an Okolab stage-top incubator for on-stage environmental control. All imaging was done using a 20×/0.75 Plan Apo DIC objective.

For fluorescence imaging, we imaged RUES2-GLR hESC-derived gastruloids (containing fluorescent reporters as follows: BRA/TBXT-mCerulean, SOX2-mCitrine, SOX17-tdTomato), using the following laser and Chroma Enhanced Transmission (ET) filter sets: mCerulean (445 nm laser; emission 480/40), mCitrine (514 nm laser; emission 535/30), and tdTomato (561 nm laser; emission 620/60). Acquisition proceeded in the order 445→514→561 to minimize cross-excitation and photobleaching. For time-lapse experiments, gastruloids were maintained at 37 °C, 5% CO_2_, and high humidity in the Okolab enclosure. Laser power and exposure times were adjusted to the minimum required to achieve adequate signal-to-noise ratio (SNR) while avoiding phototoxicity.

### Brightfield microscopy

Live imaging of human gastruloids was performed on a Nikon Ti2 motorized inverted microscope equipped with Perfect Focus System (PFS), controlled by NIS-Elements (Nikon). The system includes a Hamamatsu Flash4.0 sCMOS camera (6.5 µm^2^ photodiode), Nikon motorized stage and emission filter wheel, and an Okolab stage-top incubator for on-stage environmental control. All imaging was done using a 10x/0.45 Plan Apo λ objective.

For brightfield-only experiments, we imaged H9 SOX17mCHERRY/w;RUNX1CGFP/w hESC-derived gastruloids. Gastruloids were maintained at 37 °C, 5% CO_2_, and high humidity in the Okolab enclosure.

### Gastruloid merging experiment

For the merging experiment, gastruloids were generated as described above and in [Moris et al. *Nature* 2020a; Moris et al. *Res. Sq*. 2020b] using GLR hES cells. At 48 hours after cell seeding, two gastruloids were transferred into a new U-bottom well containing 190 µL E6 medium. Live imaging of merging process was performed using the time-lapse microscope system described above. BRA/TBXT-mCerulean and SOX17-TdTomato fluorescence and brightfield imaging were used in combination to track the merging process.

### Biophysical model

The biophysical model is detailed in the Supplemental Information.

### Platform construction

The *DevSim* (Development Simulator) platform was constructed for simulating genetic-mechanical regulatory networks and developmental dynamics, as a numerical implementation following exactly the descriptions in [METHOD DETAILS – Biophysical model]. It is based on *MATLAB R2024b* [The MathWorks Inc. 2024], leveraging MATLAB App Designer for the user interface and Parallel Computing Toolbox for efficient large-scale simulations. The platform provides a fully executable framework for the symmetry-breaking mechanisms reported in this study. After extensive testing, it has been customized to run seamlessly on personal computers (Windows 11 and macOS Sequoia 15.6.1), online webpage (https://www.mathworks.com/products/matlab-online), and high-performance servers (O2 High Performance Compute Cluster at Harvard Medical School, https://harvardmed.atlassian.net/wiki/spaces/O2). A complete user guidebook is provided in the Supplemental Text 1.

## QUANTITATIVE AND STATISTICAL ANALYSIS

### Quantifying cell loss and domain separation

We quantified two-cell-type patterns with two related readouts: cell loss, *L*, and morphological asymmetry, *A*.

For simulations, we first identified the largest connected aggregate. To do this, we built a graph in which each cell was a node. Two cells were connected if their centers were separated by less than 2*l*, where *l* is the cell radius. The largest connected component of this graph was taken as the main aggregate.

Let the full simulation contain 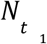 cells of type *t*_1_ and 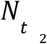 cells of type *t*_2_. Let the largest aggregate contain 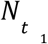‘ and 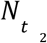 cells of these two types. Cell loss was defined as the fraction of cells outside the largest aggregate:

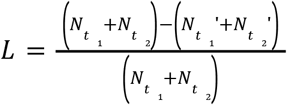

Let 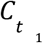 and 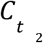 be the sets of type *t*_1_ and type *t*_2_ cells in the largest aggregate. We computed the centroid of each cell type:

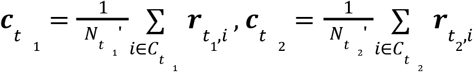

The distance between the two cell-type centroids was:

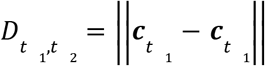

We normalized this distance by the size of the aggregate. Let 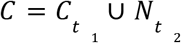 and let 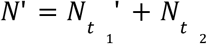. The centroid of the whole aggregate was:

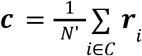

The mean distance of cells from this centroid was:

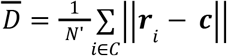

We then defined morphological asymmetry as:

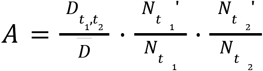

Thus, *A* is large only when the two cell types are separated and both cell types remain in the main aggregate. The retained-cell factors prevent fragmented simulations from receiving a large score simply because two small clusters drift apart.

For experimental images, individual cell positions were not available. We therefore computed a related image-based score, *a*. First, we segmented 2D anterior and posterior domains from the fluorescence images. We then rotated each 2D domain around the midline and filled the resulting 3D domain with uniformly spaced pseudo-cells. We computed *a* using the same centroid-distance normalization as above, but without a cell-loss factor:

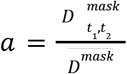

Here, 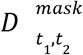 is the distance between the centroids of the two uniformly filled image masks, and *D*^*mask*^ is the mean distance of pseudo-cells from the centroid of the reconstructed aggregate.

Thus, *A* is computed from simulated cell positions and preserves non-uniform cell density. It also penalizes fragmentation. The image-based score *a* is computed from uniformly filled masks and is used to compare simulated final patterns with segmented experimental images.

## Notes

https://github.com/hormoz-lab/Symmetry-Breaking.git

### Competing Interest Statement

The authors have declared no competing interest.

### Summary of Updates

Manuscript rewritten; Figure 1 revised; Figure 5 added; Supplemental files updated.

