## Supplemental Information for "Robust axis formation requires both short-range adhesion and long-range attraction"

**Supplemental Figure**

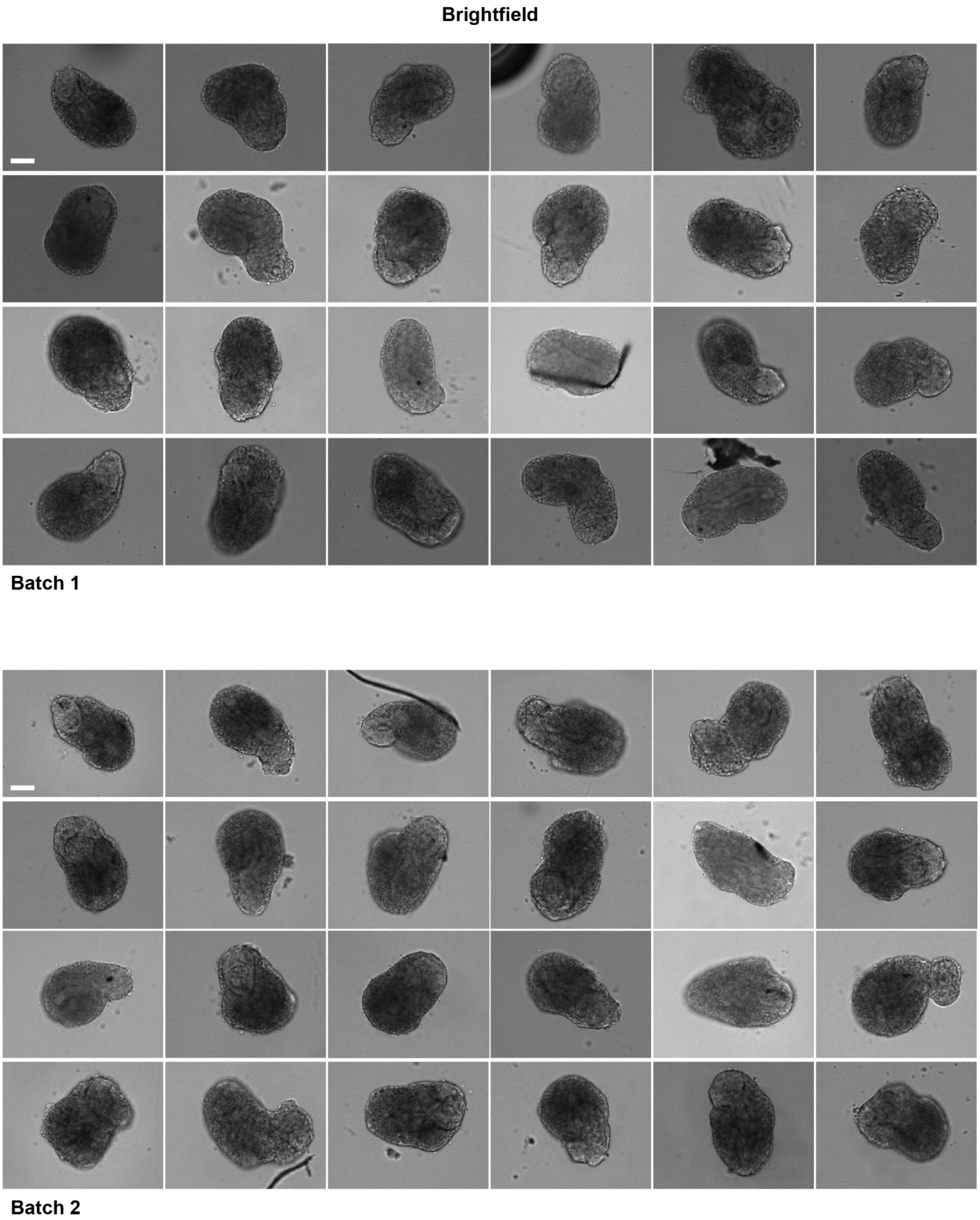


*Figure S1. Human gastruloids reproducibly elongate across independent batches.*

*Brightfield images of RUES2-GLR hESC-derived human gastruloids from two independent experimental batches at 72 h after seeding. Each batch contains 24 gastruloids. Scale bar, 100 µm.*

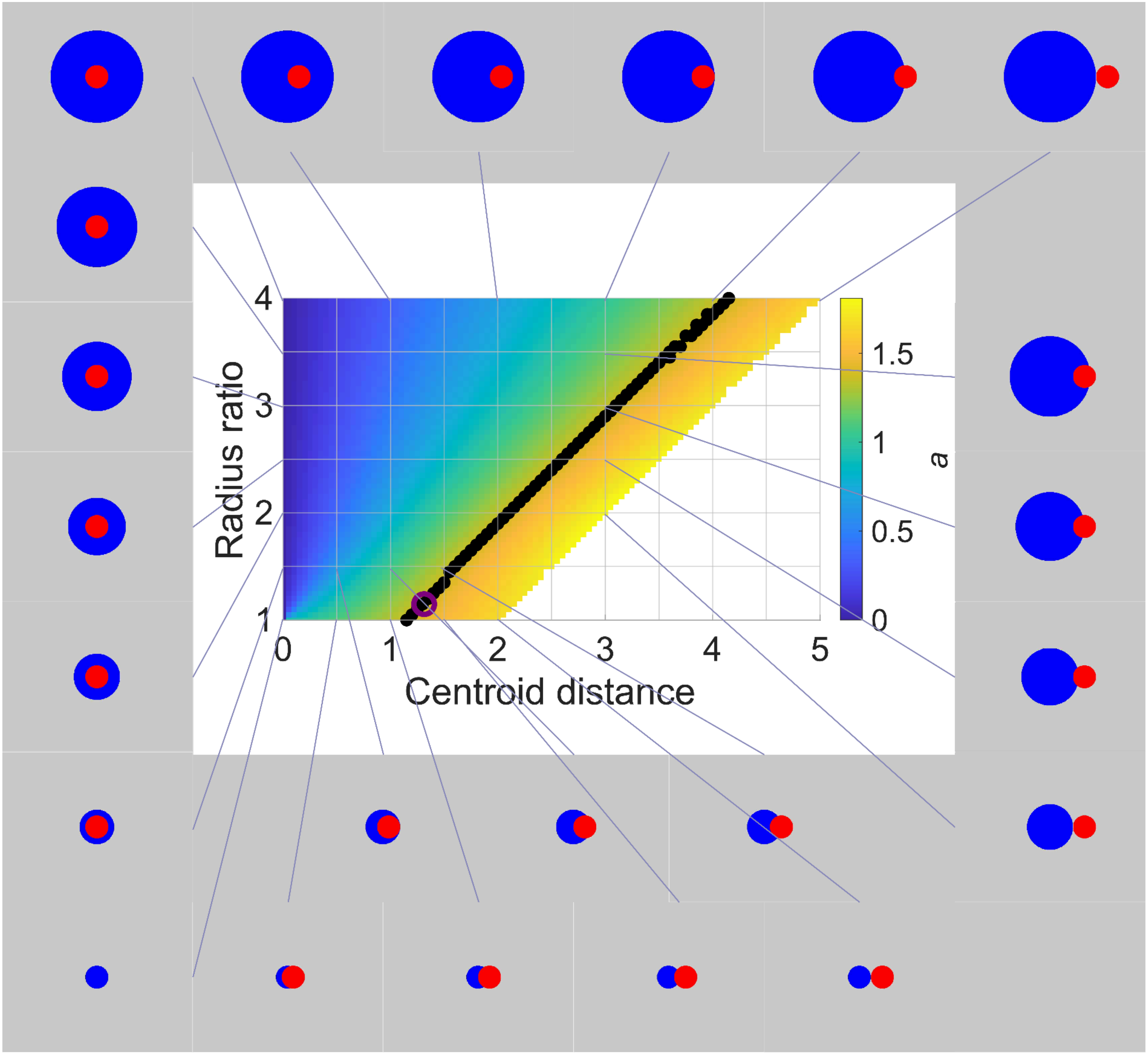


***Figure S2. Interpreting the image-based A-P separation score, α.***

*Heat map showing α for idealized two-domain geometries as a function of domain radius ratio and centroid distance. The score α was computed assuming uniform density within each domain, as in the experimental image analysis. Surrounding schematics show representative geometries. Black points mark idealized geometries with* $\alpha = 1.38$, *the mean value measured in human gastruloids. The purple circle marks the geometry with the mean radius ratio and mean centroid distance measured in human gastruloids.*

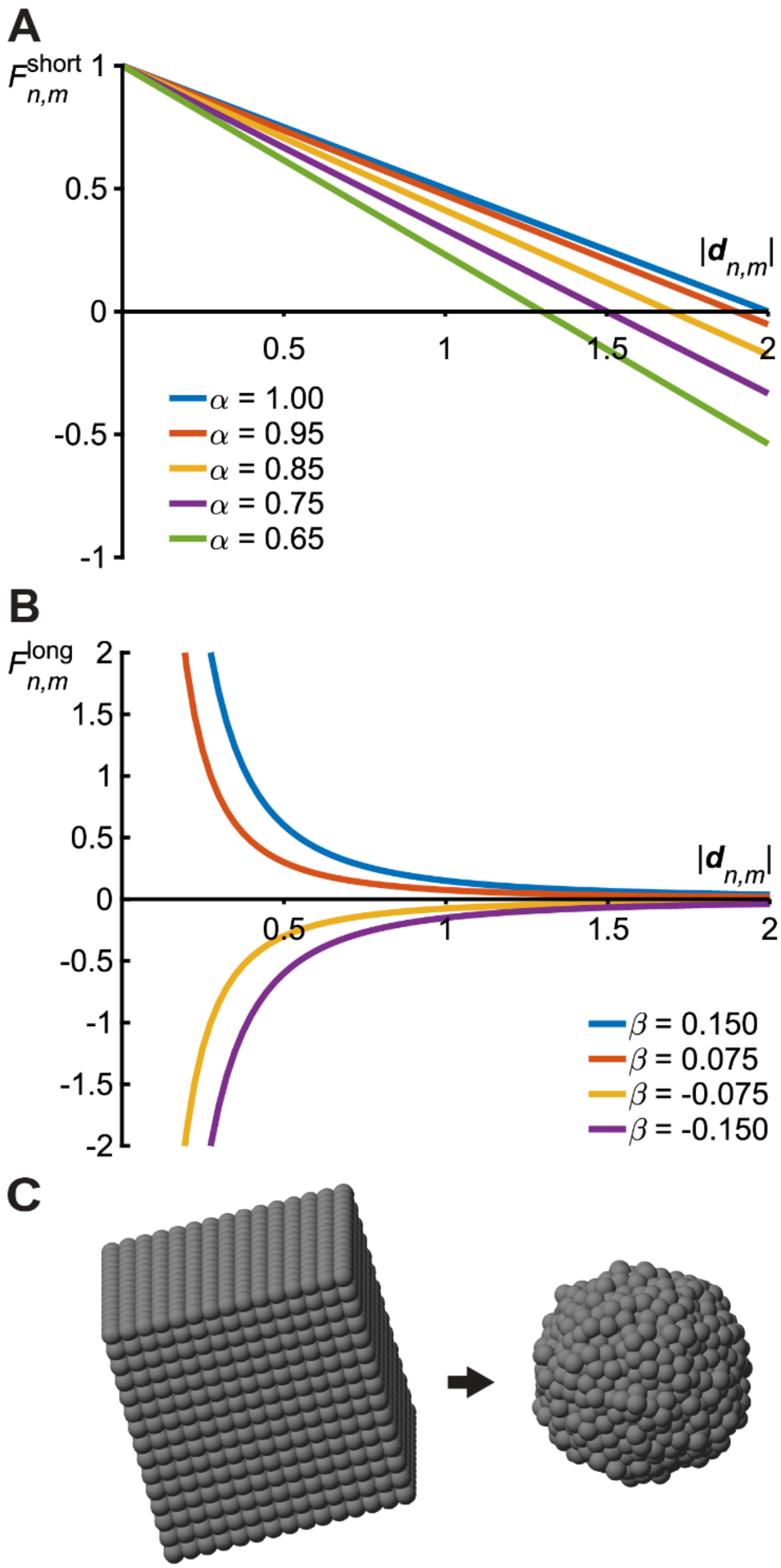


***Figure S3. Mechanical force laws and initialization of spherical aggregates.***

*(A) Short-range contact force as a function of cell-cell distance,* $d_{m,n}$*, for* $\alpha = 0.65$*,* $0.75$*,* $0.85$*,* $0.95$*, and* $1.0$*. The parameter* $\alpha$ *sets the zero-force distance,* $d_{m,n} = 2l\alpha$*. Distances below this value are repulsive. Distances above this value, within the contact range, are attractive.*

*(B) Long-range force as a function of cell-cell distance,* $d_{m,n}$*, for* $\beta = -0.150$*,* $-0.075$*,* $0.075$*, and* $0.150$*. Positive* $\beta$ *denotes attraction. Negative* $\beta$ *denotes repulsion. The force magnitude decays with distance.*

*(C) Initialization of a compact spherical aggregate. Cells were first placed on a regular* $15\times15\times15$ *cubic lattice. The positions were randomized by simulating homogeneous short-range interactions with* $\alpha = 0.80$*, time step* $\Delta T = 0.01$*, and total time* $T_{total} = 100$*. The* $1,500$ *cells closest to the aggregate centroid were retained for subsequent simulations.*

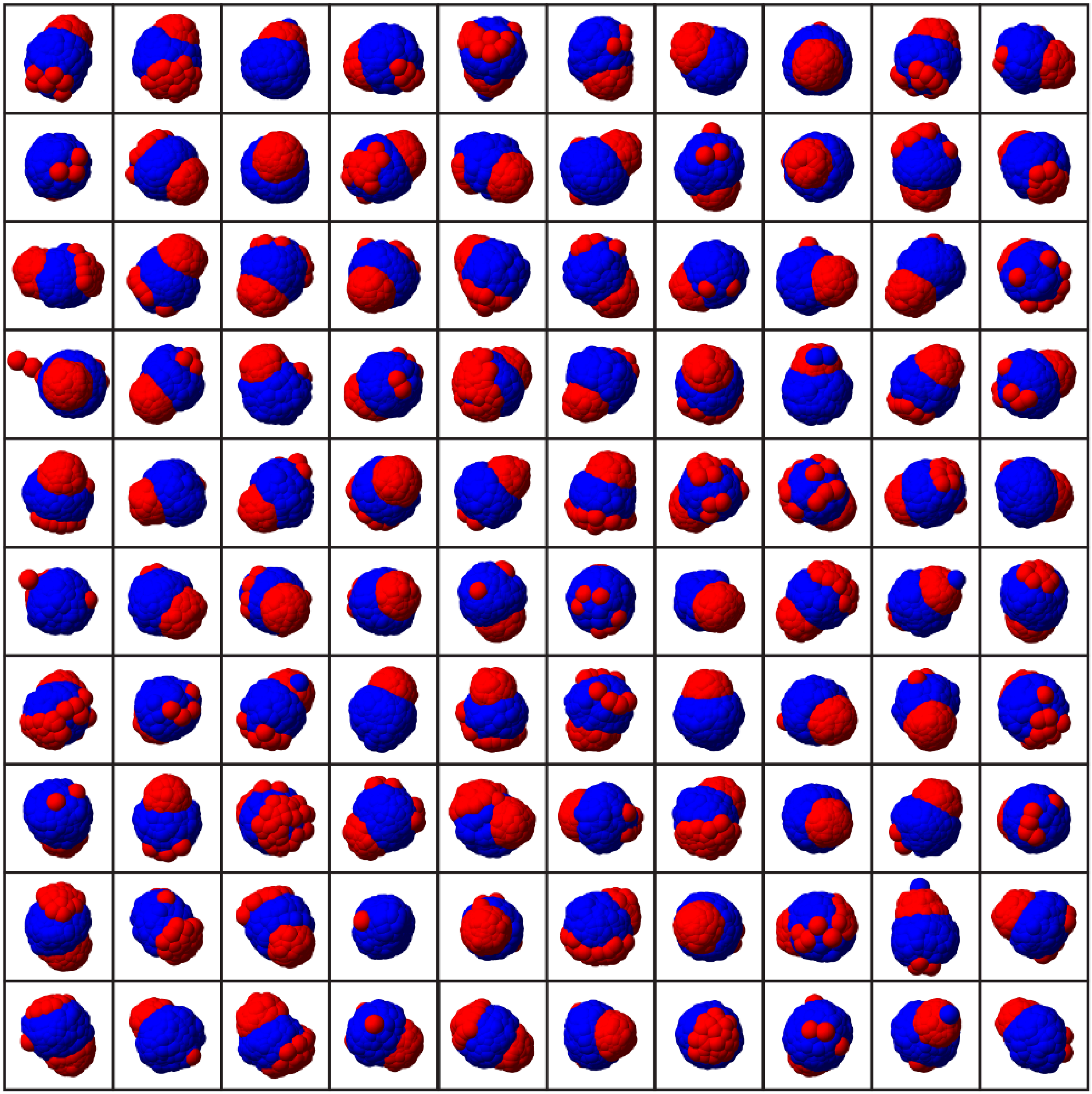

***Figure S4. Final states from 100 independent adhesion-only simulations.***

*Final states from 100 independent simulations using the best adhesion-only parameter set from Figure 3A:* $\alpha_{i-i} = 0.750$, $\alpha_{o-o} = 0.725$, *and* $\alpha_{i-o} = 0.800$. *These simulations typically show weak separation, local clustering, or unstable final structures rather than one stable axis. Five representative trajectories from this set are shown in Figure 3D.*

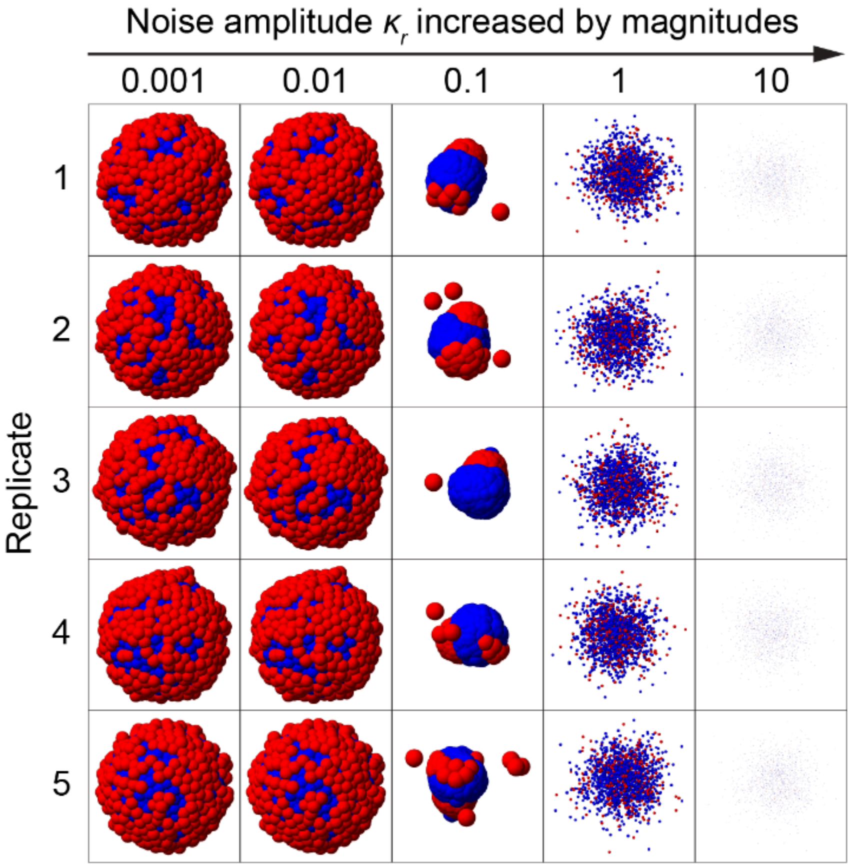


***Figure S5. Effect of noise on adhesion-only simulations.***

*Final states from simulations with different noise levels, using the adhesion parameters* $\alpha_{i-i} = 0.750$, $\alpha_{o-o} = 0.725$, *and* $\alpha_{i-o} = 0.800$. *At low noise, cells remain trapped near the initial radial configuration. At high noise, the aggregate becomes unstable. Intermediate noise allows rearrangement but does not produce a broad regime of reliable one-axis formation.*

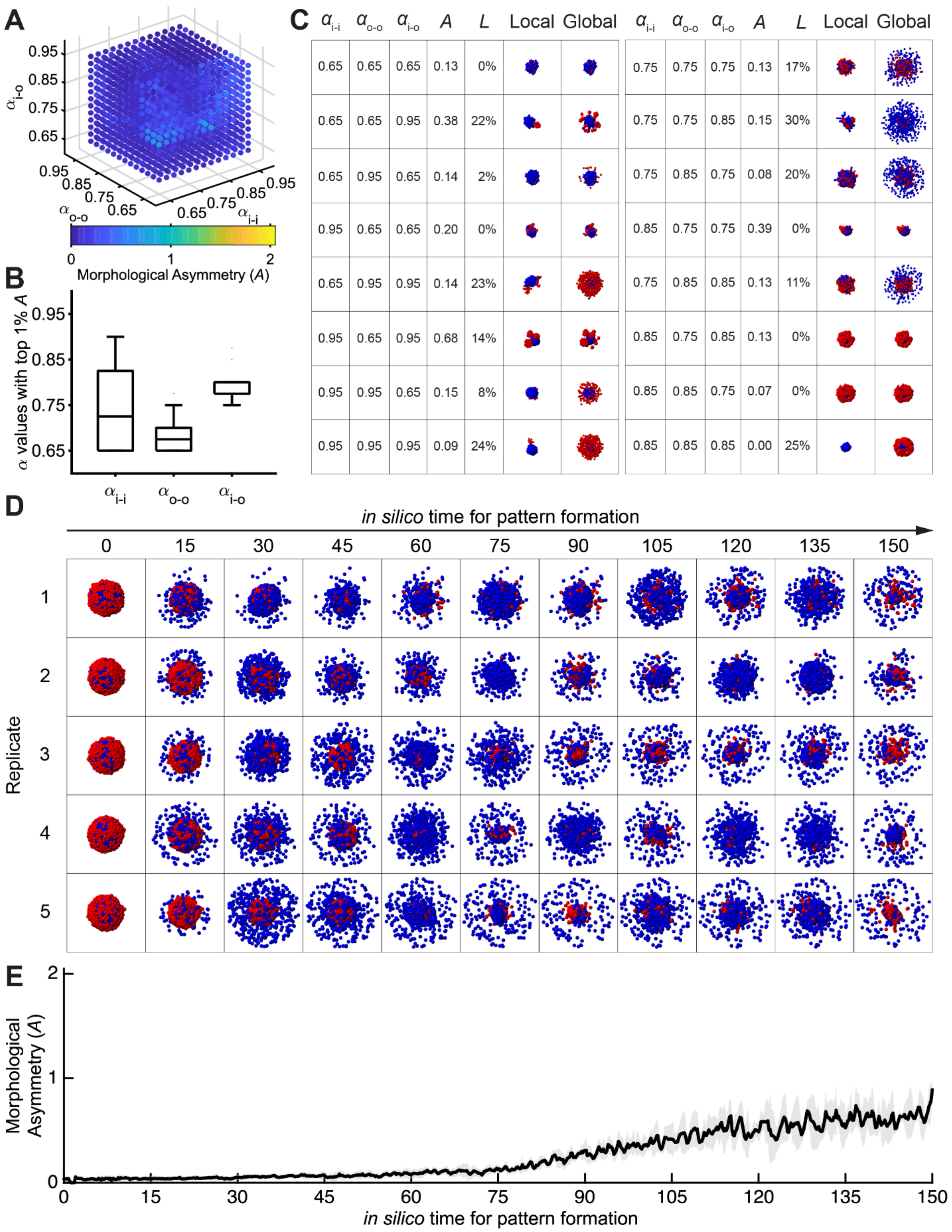

A
$\alpha_{i\text{-}o}$
0.95
0.85
0.75
0.65
$\alpha_{o\text{-}o}$
$\alpha_{i\text{-}i}$
0 1 2
Morphological Asymmetry (A)
B
α values with top 1% A
0.95
0.85
0.75
0.65
$\alpha_{i\text{-}i}$ $\alpha_{o\text{-}o}$ $\alpha_{i\text{-}o}$
C
$\alpha_{i\text{-}i}$ $\alpha_{o\text{-}o}$ $\alpha_{i\text{-}o}$ A L Local Global
0.65 0.65 0.65 0.13 0%
0.65 0.65 0.95 0.38 22%
0.65 0.95 0.65 0.14 2%
0.95 0.65 0.65 0.20 0%
0.65 0.95 0.95 0.14 23%
0.95 0.65 0.95 0.68 14%
0.95 0.95 0.65 0.15 8%
0.95 0.95 0.95 0.09 24%
$\alpha_{i\text{-}i}$ $\alpha_{o\text{-}o}$ $\alpha_{i\text{-}o}$ A L Local Global
0.75 0.75 0.75 0.13 17%
0.75 0.75 0.85 0.15 30%
0.75 0.85 0.75 0.08 20%
0.85 0.75 0.75 0.39 0%
0.75 0.85 0.85 0.13 11%
0.85 0.75 0.85 0.13 0%
0.85 0.85 0.75 0.07 0%
0.85 0.85 0.85 0.00 25%
D
in silico time for pattern formation
0 15 30 45 60 75 90 105 120 135 150
Replicate
1
2
3
4
5
E
Morphological Asymmetry (A)
2
1
0
0 15 30 45 60 75 90 105 120 135 150
in silico time for pattern formation

***Figure S6. Morphogenetic landscape with inner-to-inner long-range attraction.***

*Simulations were performed with* $\beta_{i \to i} = 0.15$ *and all other long-range interaction parameters set to zero.*

*(A) Morphological asymmetry score, A, across the three short-range adhesion parameters* $\alpha_{i-i}$, $\alpha_{o-o}$, *and* $\alpha_{i-o}$. *Blue indicates low A. Yellow indicates high A. The maximum value in this scan is* $A = 0.891$.

*(B) Distribution of* $\alpha_{i-i}$, $\alpha_{o-o}$, *and* $\alpha_{i-o}$ *among the top* $1\%$ *of simulations ranked by A. Box, interquartile range; center line, median; whiskers,* $1.5\times$ *interquartile range.*

*(C) Representative final states from selected parameter settings. The table shows the three adhesion parameters, the asymmetry score A, and the cell loss score L. "Local" shows the largest connected aggregate. "Global" shows all cells in the simulation.*

*(D) Time course of five representative simulations using the best parameter setting from this scan:* $\alpha_{i-i} = 0.800$, $\alpha_{o-o} = 0.750$, *and* $\alpha_{i-o} = 0.800$. *These simulations show weak, absent, or unstable axis formation.*

*(E) Morphological asymmetry score, A, over time for the five simulations shown in (D). Black line, mean. Gray shading, standard deviation.*

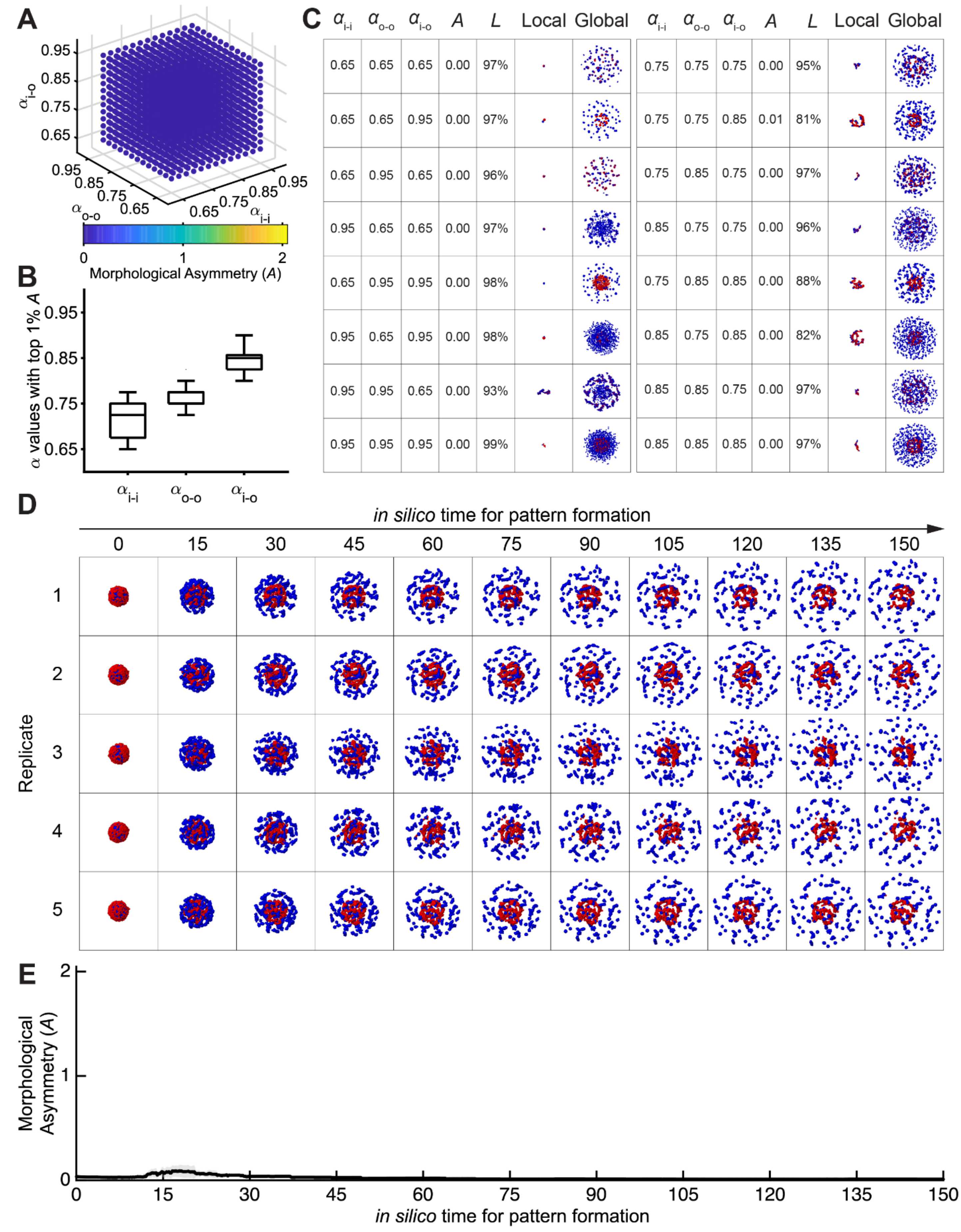


***Figure S7. Morphogenetic landscape with inner-to-inner long-range repulsion.***

*Simulations were performed with* $\beta_{i \to i} = -0.15$ *and all other long-range interaction parameters set to zero.*

*(A) Morphological asymmetry score, A, across the three short-range adhesion parameters* $\alpha_{i-i}$, $\alpha_{o-o}$, *and* $\alpha_{i-o}$. *Blue indicates low A. Yellow indicates high A. The maximum value in this scan is* $A = 0.010$.

*(B) Distribution of* $\alpha_{i-i}$, $\alpha_{o-o}$, *and* $\alpha_{i-o}$ *among the top* 1% *of simulations ranked by A. Box, interquartile range; center line, median; whiskers,* 1.5× *interquartile range.*

*(C) Representative final states from selected parameter settings. The table shows the three adhesion parameters, the asymmetry score A, and the cell loss score L. "Local" shows the largest connected aggregate. "Global" shows all cells in the simulation.*

*(D) Time course of five representative simulations using the best parameter setting from this scan:* $\alpha_{i-i} = 0.675$, $\alpha_{o-o} = 0.750$, *and* $\alpha_{i-o} = 0.825$. *These simulations show weak, absent, or unstable axis formation.*

*(E) Morphological asymmetry score, A, over time for the five simulations shown in (D). Black line, mean. Gray shading, standard deviation.*

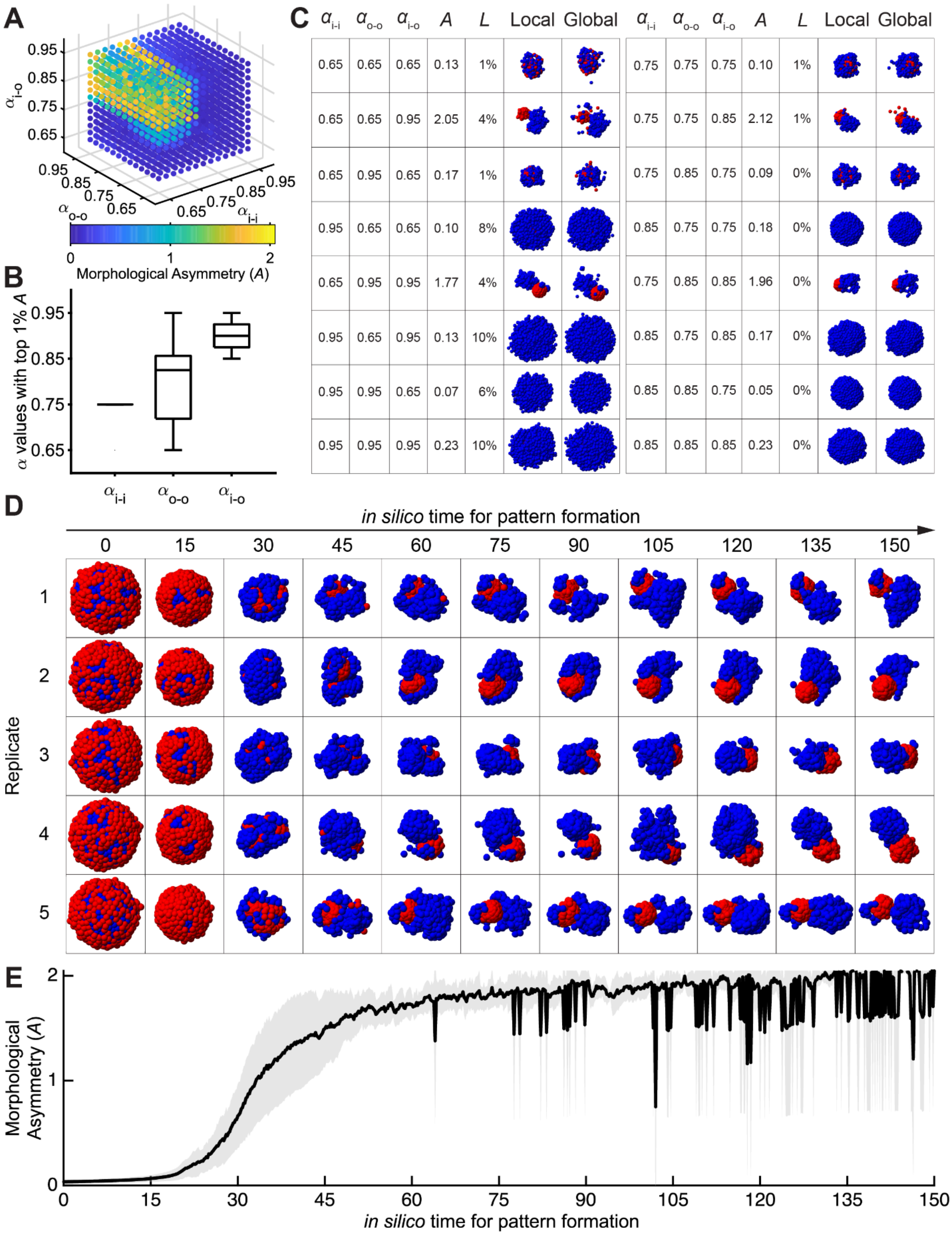


*Figure S8. Morphogenetic landscape with outer-to-outer long-range attraction.*

*Simulations were performed with* $\beta_{o \to o} = 0.15$ *and all other long-range interaction parameters set to zero.*

*(A) Morphological asymmetry score, A, across the three short-range adhesion parameters* $\alpha_{i-i}$, $\alpha_{o-o}$, *and* $\alpha_{i-o}$. *Blue indicates low A. Yellow indicates high A. The maximum value in this scan is* $A = 2.047$.

*(B) Distribution of* $\alpha_{i-i}$, $\alpha_{o-o}$, *and* $\alpha_{i-o}$ *among the top* $1\%$ *of simulations ranked by A. Box, interquartile range; center line, median; whiskers,* $1.5\times$ *interquartile range.*

*(C) Representative final states from selected parameter settings. The table shows the three adhesion parameters, the asymmetry score A, and the cell loss score L. "Local" shows the largest connected aggregate. "Global" shows all cells in the simulation.*

*(D) Time course of five representative simulations using the best parameter setting from this scan:* $\alpha_{i-i} = 0.750$, $\alpha_{o-o} = 0.825$, *and* $\alpha_{i-o} = 0.950$. *These simulations form one stable axis.*

*(E) Morphological asymmetry score, A, over time for the five simulations shown in (D). Black line, mean. Gray shading, standard deviation.*

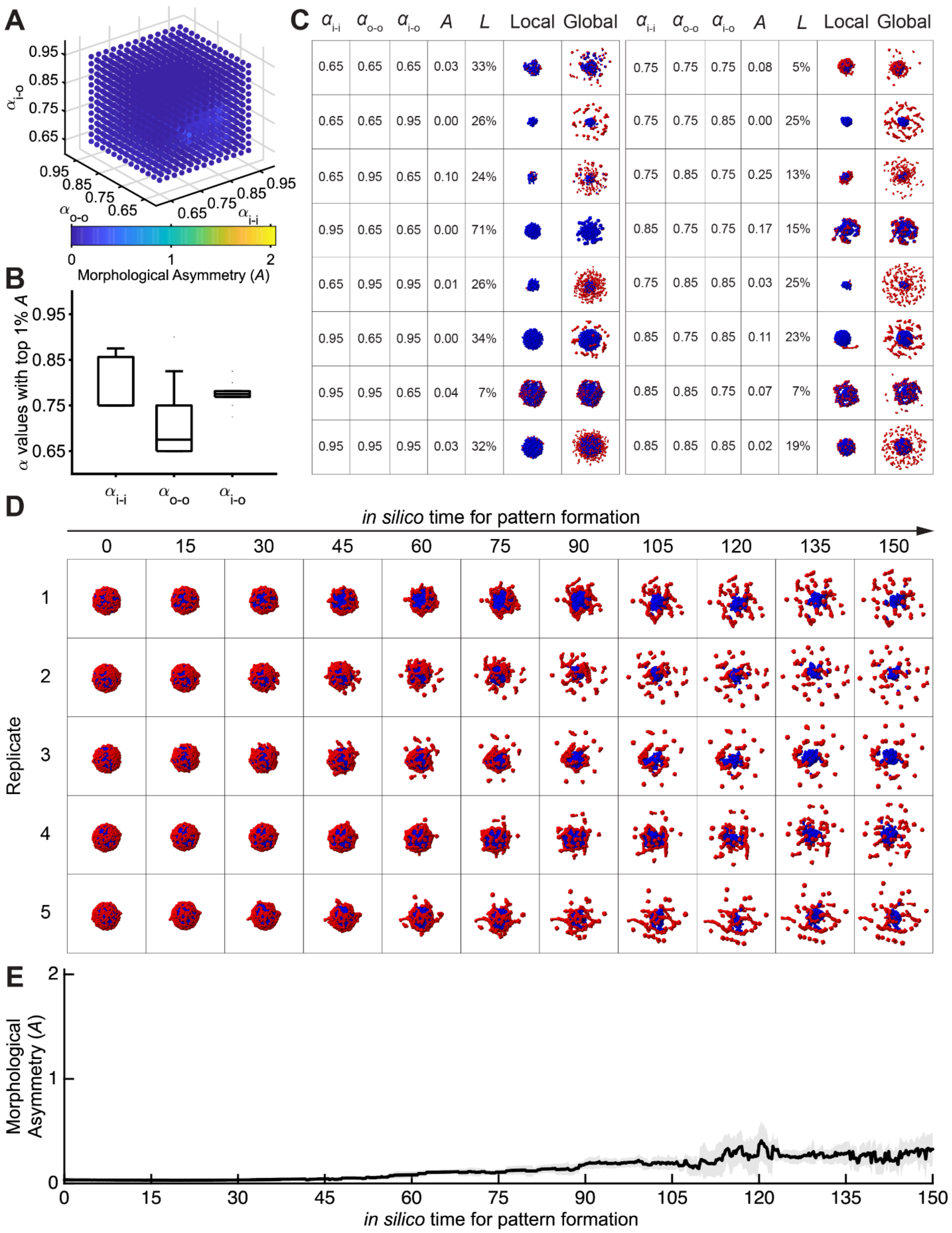

A
0.95
0.85
0.75
0.65
$\alpha_{i-o}$
0.95
0.85
0.75
0.65
$\alpha_{o-o}$
0.65
0.75
0.85
0.95
$\alpha_{i-i}$
0
1
2
Morphological Asymmetry (A)
B
α values with top 1% A
0.95
0.85
0.75
0.65
$\alpha_{i-i}$
$\alpha_{o-o}$
$\alpha_{i-o}$
C
$\alpha_{i-i}$ $\alpha_{o-o}$ $\alpha_{i-o}$ A L Local Global $\alpha_{i-i}$ $\alpha_{o-o}$ $\alpha_{i-o}$ A L Local Global
0.65 0.65 0.65 0.03 33% 0.75 0.75 0.75 0.08 5%
0.65 0.65 0.95 0.00 26% 0.75 0.75 0.85 0.00 25%
0.65 0.95 0.65 0.10 24% 0.75 0.85 0.75 0.25 13%
0.95 0.65 0.65 0.00 71% 0.85 0.75 0.75 0.17 15%
0.65 0.95 0.95 0.01 26% 0.75 0.85 0.85 0.03 25%
0.95 0.65 0.95 0.00 34% 0.85 0.75 0.85 0.11 23%
0.95 0.95 0.65 0.04 7% 0.85 0.85 0.75 0.07 7%
0.95 0.95 0.95 0.03 32% 0.85 0.85 0.85 0.02 19%
D
in silico time for pattern formation
0
15
30
45
60
75
90
105
120
135
150
Replicate
1
2
3
4
5
E
Morphological Asymmetry (A)
2
1
0
0
15
30
45
60
75
90
105
120
135
150
in silico time for pattern formation

***Figure S9. Morphogenetic landscape with outer-to-outer long-range repulsion.***

*Simulations were performed with* $\beta_{o\to o} = -0.15$ *and all other long-range interaction parameters set to zero.*

*(A) Morphological asymmetry score, A, across the three short-range adhesion parameters* $\alpha_{i-i}$, $\alpha_{o-o}$, *and* $\alpha_{i-o}$. *Blue indicates low A. Yellow indicates high A. The maximum value in this scan is* $A = 0.328$.

*(B) Distribution of* $\alpha_{i-i}$, $\alpha_{o-o}$, *and* $\alpha_{i-o}$ *among the top* $1\%$ *of simulations ranked by A. Box, interquartile range; center line, median; whiskers,* $1.5\times$ *interquartile range.*

*(C) Representative final states from selected parameter settings. The table shows the three adhesion parameters, the asymmetry score A, and the cell loss score L. "Local" shows the largest connected aggregate. "Global" shows all cells in the simulation.*

*(D) Time course of five representative simulations using the best parameter setting from this scan:* $\alpha_{i-i} = 0.750$, $\alpha_{o-o} = 0.650$, *and* $\alpha_{i-o} = 0.775$. *These simulations show weak, absent, or unstable axis formation.*

*(E) Morphological asymmetry score, A, over time for the five simulations shown in (D). Black line, mean. Gray shading, standard deviation.*

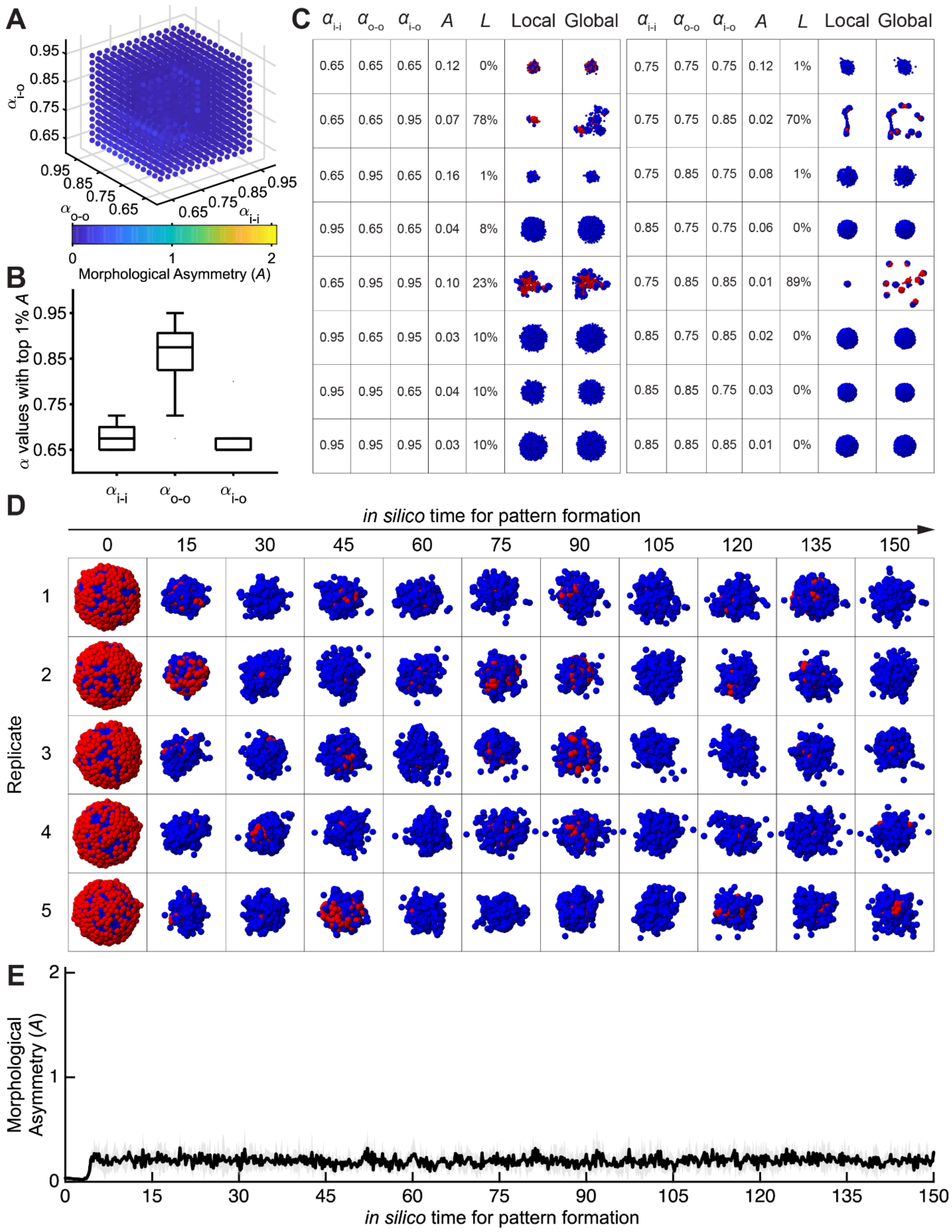
A
$\alpha_{i-o}$
0.95
0.85
0.75
0.65
0.95
0.85
0.75
$\alpha_{o-o}$
0.65
0.65
0.75
0.85
0.95
$\alpha_{i-i}$
0
1
2
Morphological Asymmetry ($A$)
B
$\alpha$ values with top 1% $A$
0.95
0.85
0.75
0.65
$\alpha_{i-i}$
$\alpha_{o-o}$
$\alpha_{i-o}$
C
$\alpha_{i-i}$ $\alpha_{o-o}$ $\alpha_{i-o}$ $A$ $L$ Local Global
0.65 0.65 0.65 0.12 0%
0.65 0.65 0.95 0.07 78%
0.65 0.95 0.65 0.16 1%
0.95 0.65 0.65 0.04 8%
0.65 0.95 0.95 0.10 23%
0.95 0.65 0.95 0.03 10%
0.95 0.95 0.65 0.04 10%
0.95 0.95 0.95 0.03 10%
$\alpha_{i-i}$ $\alpha_{o-o}$ $\alpha_{i-o}$ $A$ $L$ Local Global
0.75 0.75 0.75 0.12 1%
0.75 0.75 0.85 0.02 70%
0.75 0.85 0.75 0.08 1%
0.85 0.75 0.75 0.06 0%
0.75 0.85 0.85 0.01 89%
0.85 0.75 0.85 0.02 0%
0.85 0.85 0.75 0.03 0%
0.85 0.85 0.85 0.01 0%
D
*in silico* time for pattern formation
0 15 30 45 60 75 90 105 120 135 150
Replicate
1
2
3
4
5
E
Morphological Asymmetry ($A$)
2
1
0
0 15 30 45 60 75 90 105 120 135 150
*in silico* time for pattern formation

***Figure S10. Morphogenetic landscape with inner-to-outer long-range attraction.***

*Simulations were performed with* $\beta_{i \to o} = 0.15$ *and all other long-range interaction parameters set to zero.*

*(A) Morphological asymmetry score, A, across the three short-range adhesion parameters* $\alpha_{i-i}$*,* $\alpha_{o-o}$*, and* $\alpha_{i-o}$*. Blue indicates low A. Yellow indicates high A. The maximum value in this scan is* $A = 0.282$.

*(B) Distribution of* $\alpha_{i-i}$*,* $\alpha_{o-o}$*, and* $\alpha_{i-o}$ *among the top* $1\%$ *of simulations ranked by A. Box, interquartile range; center line, median; whiskers,* $1.5\times$ *interquartile range.*

*(C) Representative final states from selected parameter settings. The table shows the three adhesion parameters, the asymmetry score A, and the cell loss score L. "Local" shows the largest connected aggregate. "Global" shows all cells in the simulation.*

*(D) Time course of five representative simulations using the best parameter setting from this scan:* $\alpha_{i-i} = 0.650$, $\alpha_{o-o} = 0.875$*, and* $\alpha_{i-o} = 0.650$*. These simulations show weak, absent, or unstable axis formation.*

*(E) Morphological asymmetry score, A, over time for the five simulations shown in (D). Black line, mean. Gray shading, standard deviation.*

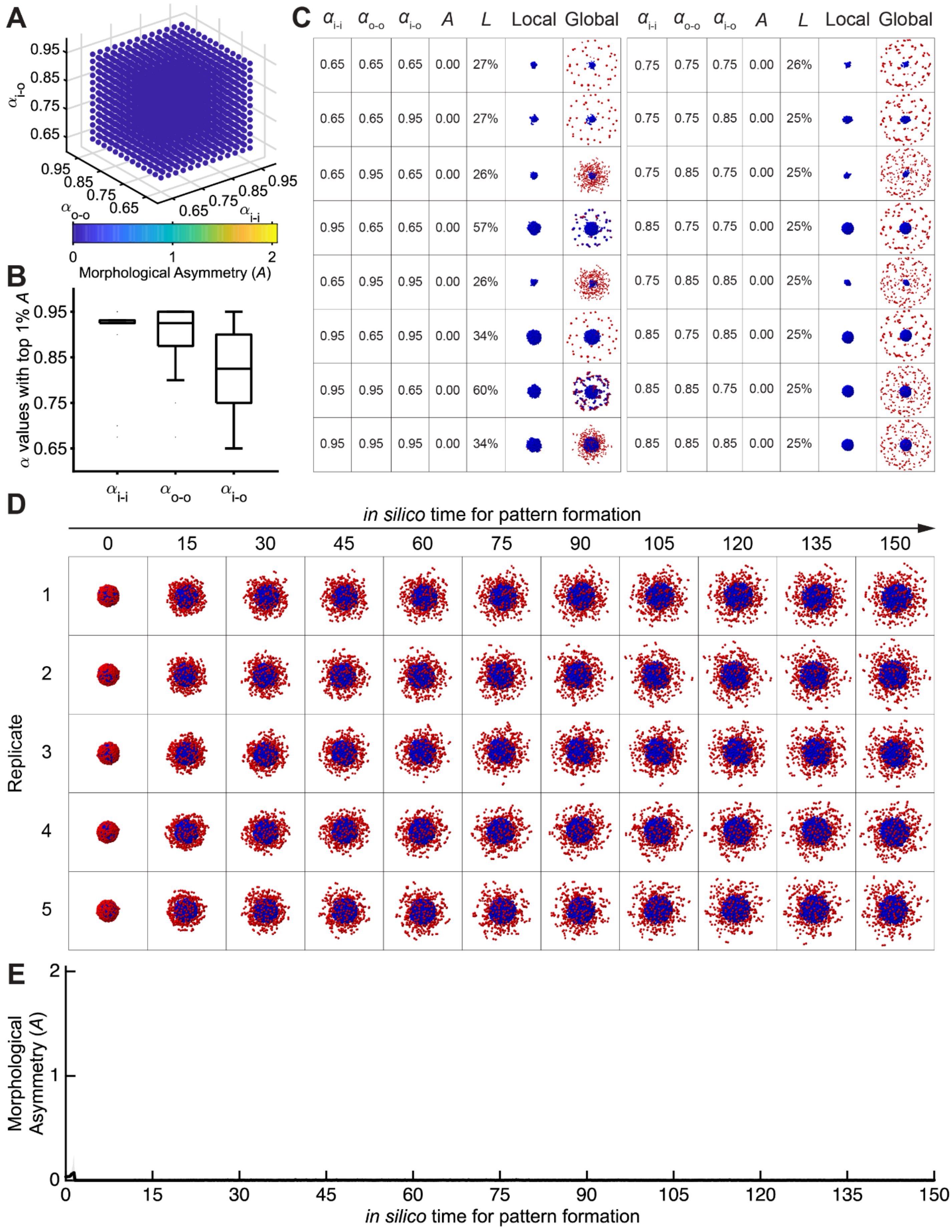


| $\alpha_{i\text{-}i}$ | $\alpha_{o\text{-}o}$ | $\alpha_{i\text{-}o}$ | A | L | Local | Global | $\alpha_{i\text{-}i}$ | $\alpha_{o\text{-}o}$ | $\alpha_{i\text{-}o}$ | A | L | Local | Global |
|---|---|---|---|---|---|---|---|---|---|---|---|---|---|
| 0.65 | 0.65 | 0.65 | 0.00 | 27% | | | 0.75 | 0.75 | 0.75 | 0.00 | 26% | | |
| 0.65 | 0.65 | 0.95 | 0.00 | 27% | | | 0.75 | 0.75 | 0.85 | 0.00 | 25% | | |
| 0.65 | 0.95 | 0.65 | 0.00 | 26% | | | 0.75 | 0.85 | 0.75 | 0.00 | 25% | | |
| 0.95 | 0.65 | 0.65 | 0.00 | 57% | | | 0.85 | 0.75 | 0.75 | 0.00 | 25% | | |
| 0.65 | 0.95 | 0.95 | 0.00 | 26% | | | 0.75 | 0.85 | 0.85 | 0.00 | 25% | | |
| 0.95 | 0.65 | 0.95 | 0.00 | 34% | | | 0.85 | 0.75 | 0.85 | 0.00 | 25% | | |
| 0.95 | 0.95 | 0.65 | 0.00 | 60% | | | 0.85 | 0.85 | 0.75 | 0.00 | 25% | | |
| 0.95 | 0.95 | 0.95 | 0.00 | 34% | | | 0.85 | 0.85 | 0.85 | 0.00 | 25% | | |

***Figure S11. Morphogenetic landscape with inner-to-outer long-range repulsion.***

*Simulations were performed with* $\beta_{i \to o} = -0.15$ *and all other long-range interaction parameters set to zero.*

*(A) Morphological asymmetry score, A, across the three short-range adhesion parameters* $\alpha_{i-i}$, $\alpha_{o-o}$, *and* $\alpha_{i-o}$. *Blue indicates low A. Yellow indicates high A. The maximum value in this scan is* $A = 0.003$.

*(B) Distribution of* $\alpha_{i-i}$, $\alpha_{o-o}$, *and* $\alpha_{i-o}$ *among the top* $1\%$ *of simulations ranked by A. Box, interquartile range; center line, median; whiskers,* $1.5\times$ *interquartile range.*

*(C) Representative final states from selected parameter settings. The table shows the three adhesion parameters, the asymmetry score A, and the cell loss score L. "Local" shows the largest connected aggregate. "Global" shows all cells in the simulation.*

*(D) Time course of five representative simulations using the best parameter setting from this scan:* $\alpha_{i-i} = 0.950$, $\alpha_{o-o} = 0.950$, *and* $\alpha_{i-o} = 0.900$. *These simulations show weak, absent, or unstable axis formation.*

*(E) Morphological asymmetry score, A, over time for the five simulations shown in (D). Black line, mean. Gray shading, standard deviation.*

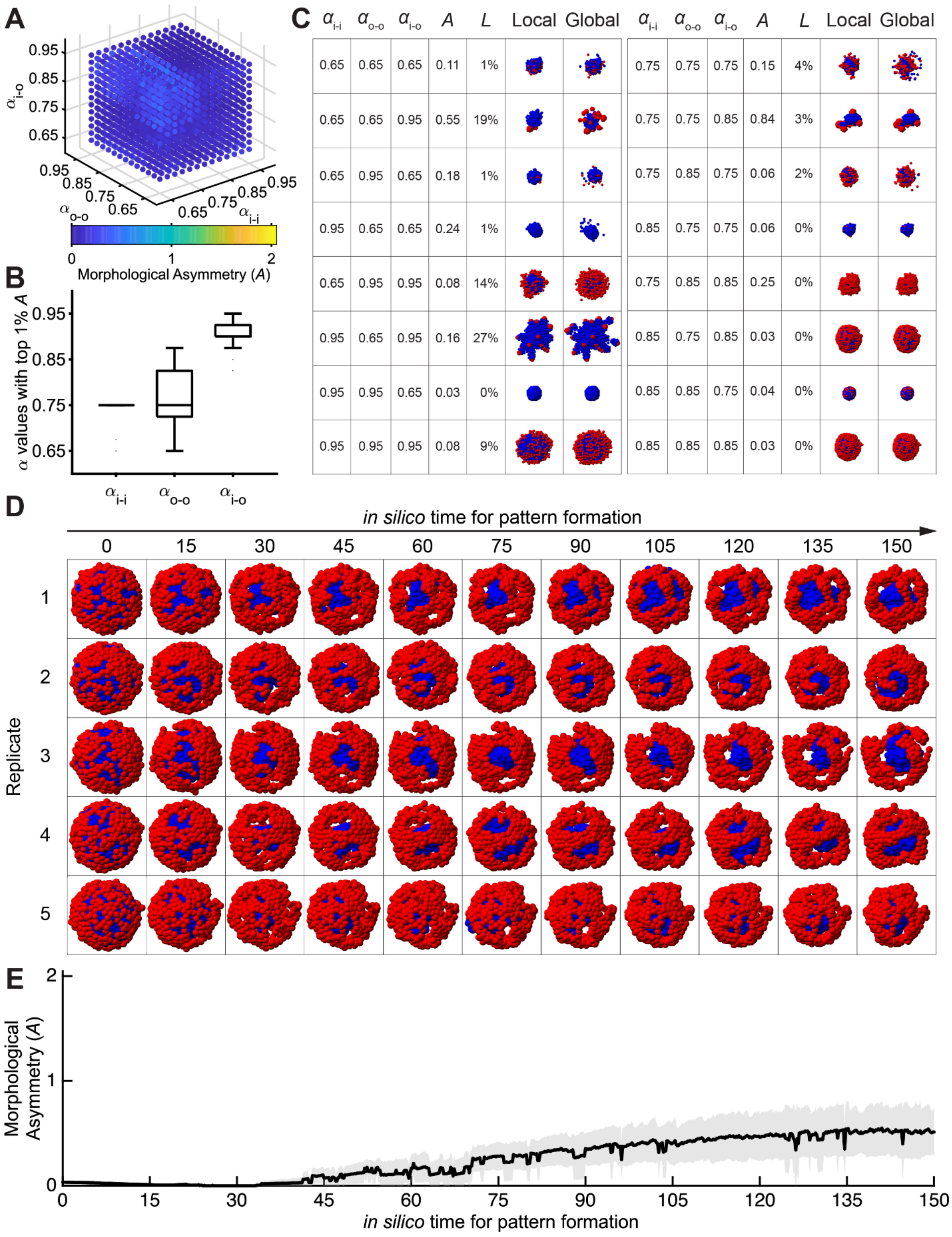


*Figure S12. Morphogenetic landscape with outer-to-inner long-range attraction.*

*Simulations were performed with* $\beta_{o \to i} = 0.15$ *and all other long-range interaction parameters set to zero.*

*(A) Morphological asymmetry score, A, across the three short-range adhesion parameters* $\alpha_{i-i}$*,* $\alpha_{o-o}$*, and* $\alpha_{i-o}$*. Blue indicates low A. Yellow indicates high A. The maximum value in this scan is* $A = 0.513$.

*(B) Distribution of* $\alpha_{i-i}$*,* $\alpha_{o-o}$*, and* $\alpha_{i-o}$ *among the top* 1% *of simulations ranked by A. Box, interquartile range; center line, median; whiskers,* $1.5\times$ *interquartile range.*

*(C) Representative final states from selected parameter settings. The table shows the three adhesion parameters, the asymmetry score A, and the cell loss score L. "Local" shows the largest connected aggregate. "Global" shows all cells in the simulation.*

*(D) Time course of five representative simulations using the best parameter setting from this scan:* $\alpha_{i-i} = 0.750$*,* $\alpha_{o-o} = 0.750$*, and* $\alpha_{i-o} = 0.950$*. These simulations show weak, absent, or unstable axis formation.*

*(E) Morphological asymmetry score, A, over time for the five simulations shown in (D). Black line, mean. Gray shading, standard deviation.*

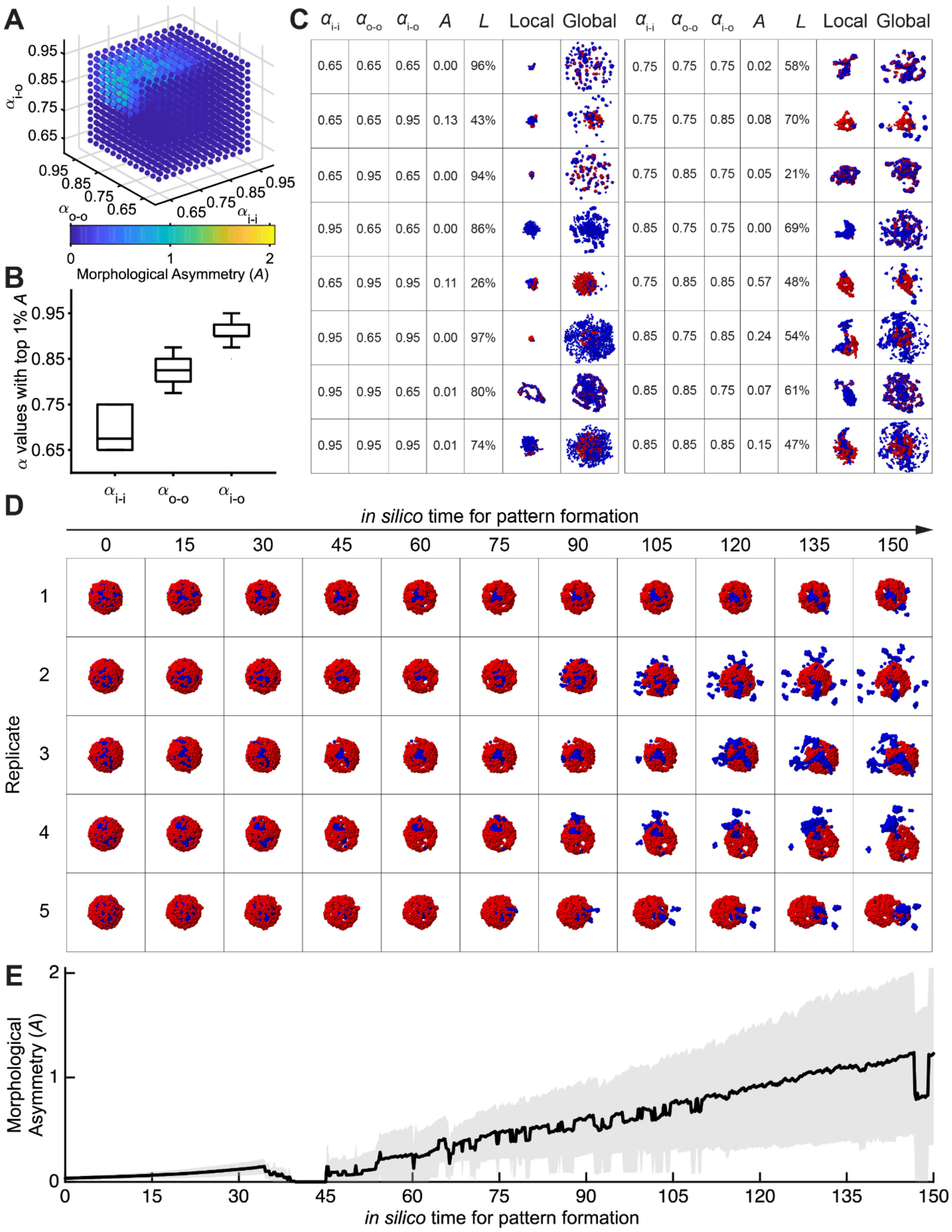
A
$\alpha_{i-o}$
0.95
0.85
0.75
0.65
$\alpha_{o-o}$
$\alpha_{i-i}$
0
1
2
Morphological Asymmetry (A)
B
α values with top 1% A
$\alpha_{i-i}$
$\alpha_{o-o}$
$\alpha_{i-o}$
C
$\alpha_{i-i}$ $\alpha_{o-o}$ $\alpha_{i-o}$ A L Local Global
0.65 0.65 0.65 0.00 96%
0.65 0.65 0.95 0.13 43%
0.65 0.95 0.65 0.00 94%
0.95 0.65 0.65 0.00 86%
0.65 0.95 0.95 0.11 26%
0.95 0.65 0.95 0.00 97%
0.95 0.95 0.65 0.01 80%
0.95 0.95 0.95 0.01 74%
0.75 0.75 0.75 0.02 58%
0.75 0.75 0.85 0.08 70%
0.75 0.85 0.75 0.05 21%
0.85 0.75 0.75 0.00 69%
0.75 0.85 0.85 0.57 48%
0.85 0.75 0.85 0.24 54%
0.85 0.85 0.75 0.07 61%
0.85 0.85 0.85 0.15 47%
D
in silico time for pattern formation
0 15 30 45 60 75 90 105 120 135 150
Replicate
1
2
3
4
5
E
Morphological Asymmetry (A)
0
1
2
0 15 30 45 60 75 90 105 120 135 150
in silico time for pattern formation

***Figure S13. Morphogenetic landscape with outer-to-inner long-range repulsion.***

*Simulations were performed with* $\beta_{o\to i} =- 0.15$ *and all other long-range interaction parameters set to zero.*

*(A) Morphological asymmetry score, A, across the three short-range adhesion parameters* $\alpha_{i-i}$, $\alpha_{o-o}$, *and* $\alpha_{i-o}$. *Blue indicates low A. Yellow indicates high A. The maximum value in this scan is* $A = 1.229$.

*(B) Distribution of* $\alpha_{i-i}$, $\alpha_{o-o}$, *and* $\alpha_{i-o}$ *among the top* 1% *of simulations ranked by A. Box, interquartile range; center line, median; whiskers,* 1.5× *interquartile range.*

*(C) Representative final states from selected parameter settings. The table shows the three adhesion parameters, the asymmetry score A, and the cell loss score L. "Local" shows the largest connected aggregate. "Global" shows all cells in the simulation.*

*(D) Time course of five representative simulations using the best parameter setting from this scan:* $\alpha_{i-i} = 0.750$, $\alpha_{o-o} = 0.800$, *and* $\alpha_{i-o} = 0.925$. *These simulations show partial axis formation, but less reliable separation than outer-to-outer attraction.*

*(E) Morphological asymmetry score, A, over time for the five simulations shown in (D). Black line, mean. Gray shading, standard deviation.*

A

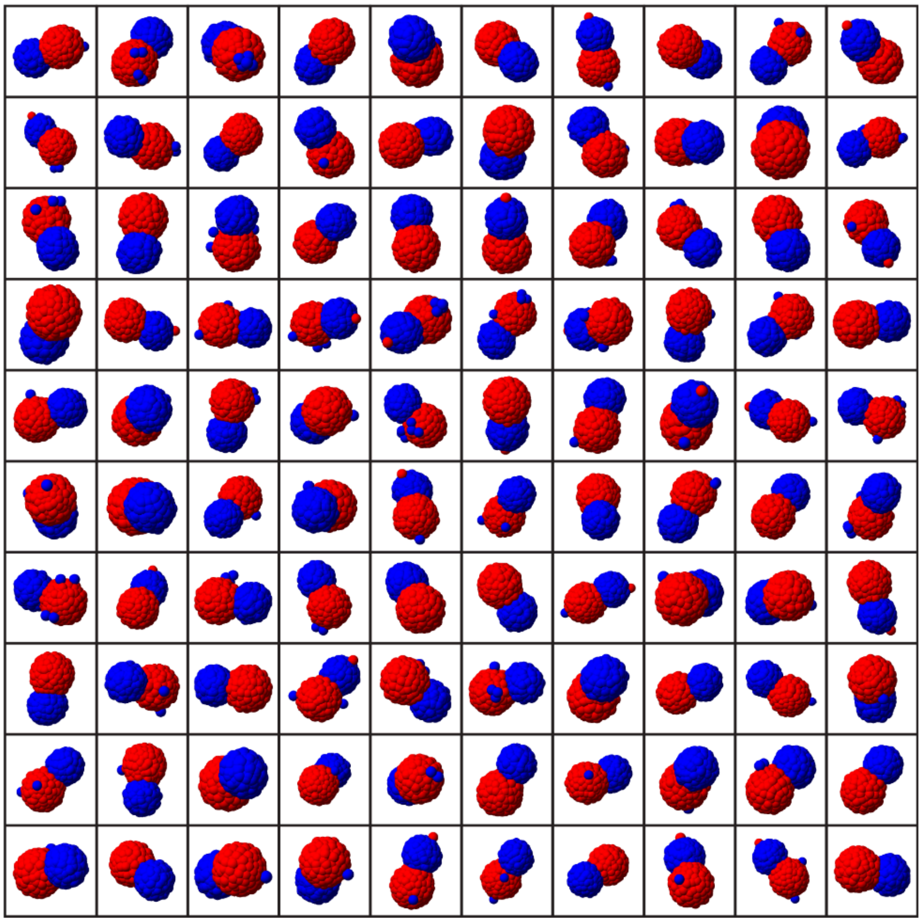

B

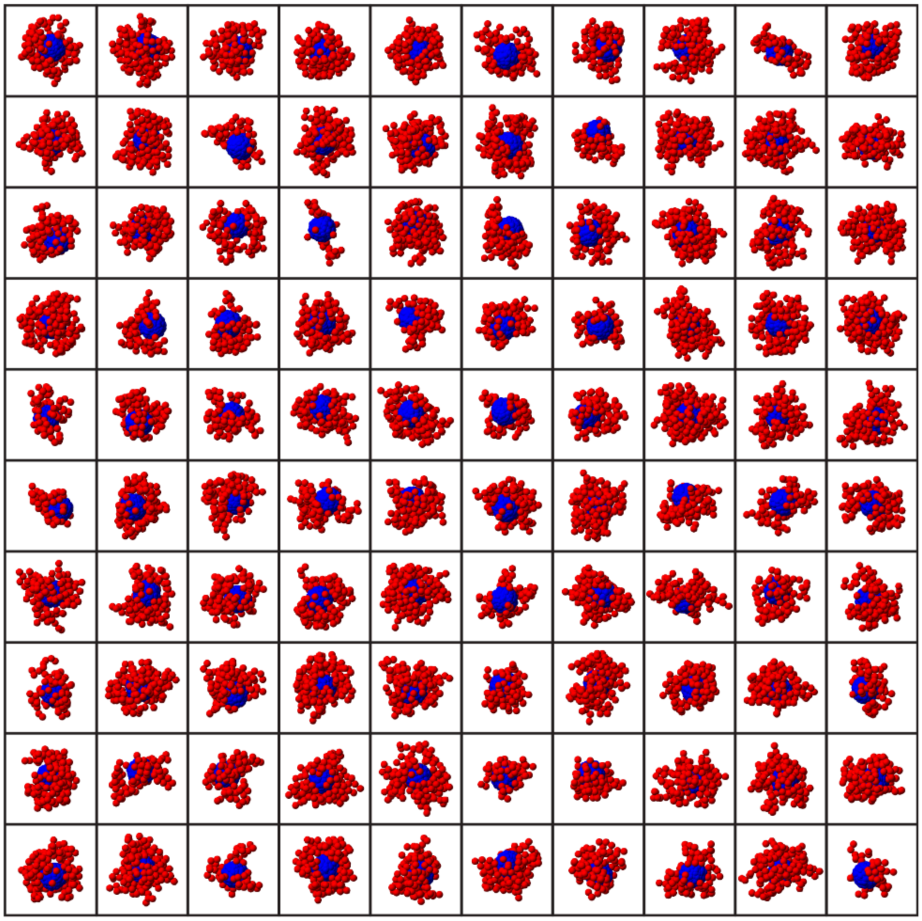

***Figure S14. Final states from 100 independent simulations with and without outer-to-outer long-range attraction.***

*Final states from 100 independent simulations using the adhesion parameters* $\alpha_{i-i} = 0.775$, $\alpha_{o-o} = 0.950$, *and* $\alpha_{i-o} = 0.875$. *This adhesion setting is from the top* 1% *of simulations ranked by A in Figure 4A.*

*(A) Simulations with outer-to-outer long-range attraction,* $\beta_{o\to o} = 0.135$. *Most simulations form one stable axis. Five representative trajectories from this set are shown in Figure 4B, and their final states are shown in Figure 4D.*

*(B) Simulations with the same adhesion parameters but without outer-to-outer long-range attraction,* $\beta_{o\to o} = 0$. *These simulations show weak separation or local clustering rather than one stable axis.*

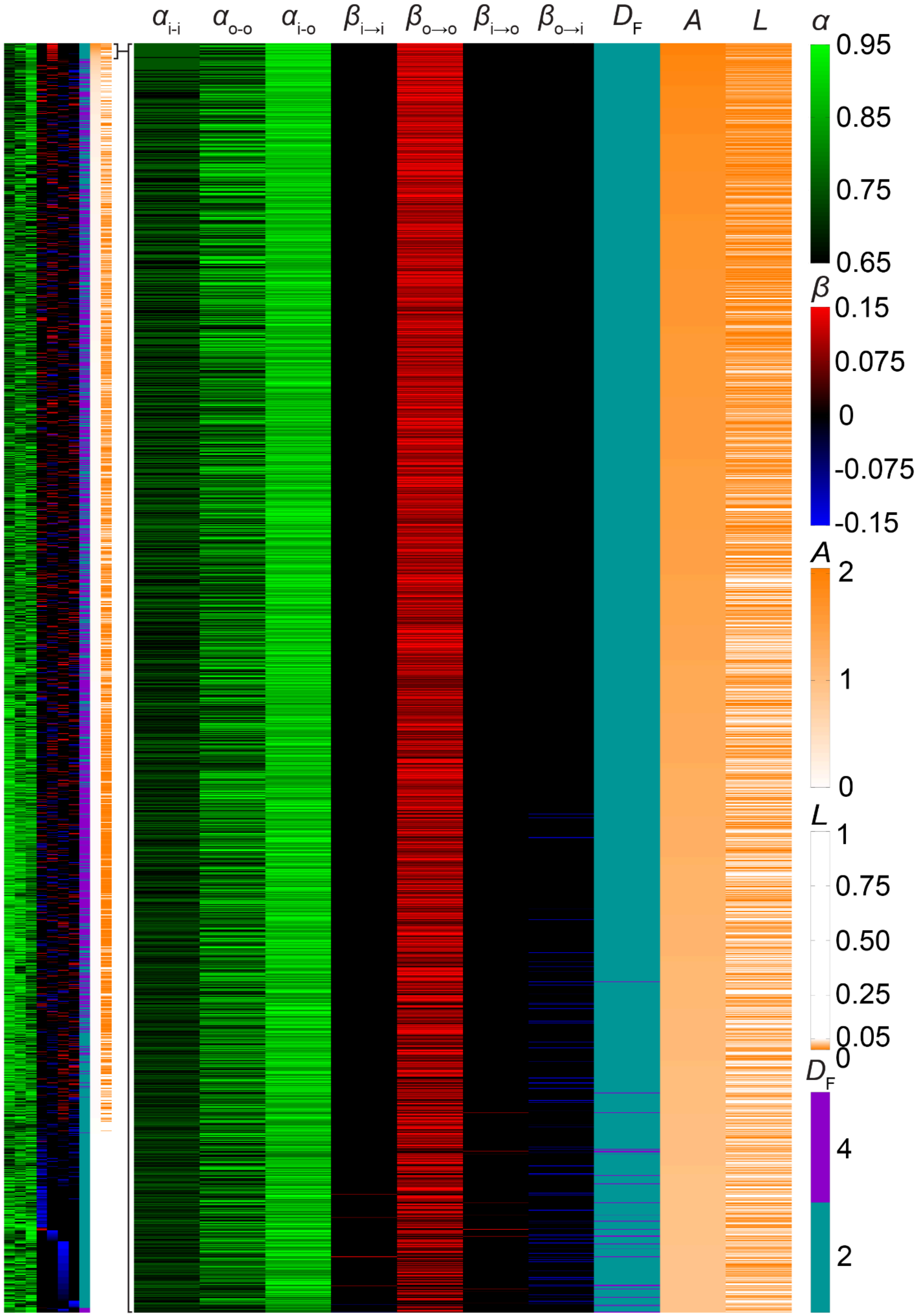

$\alpha_{i-i}$
$\alpha_{o-o}$
$\alpha_{i-o}$
$\beta_{i\to i}$
$\beta_{o\to o}$
$\beta_{i\to o}$
$\beta_{o\to i}$
$D_F$
$A$
$L$
$\alpha$
0.95
0.85
0.75
0.65
$\beta$
0.15
0.075
0
-0.075
-0.15
$A$
2
1
0
$L$
1
0.75
0.50
0.25
0.05
0
$D_F$
4
2

***Figure S15. Outer-to-outer attraction remains enriched when long-range forces decay more slowly.***

*Parameter map for simulations that include two distance-decay exponents for the long-range force,* $D_F = 2$ *and* $D_F = 4$*. Each row is one parameter setting. Rows are ranked by the morphological asymmetry score, A. The left map shows all* $355,914$ *simulated conditions. The right map enlarges the top* $1\%$ *of conditions ranked by A. Columns show the three short-range adhesion parameters,* $\alpha_{i-i}$*,* $\alpha_{o-o}$*, and* $\alpha_{i-o}$*; the four long-range interaction parameters,* $\beta_{i\to i}$*,* $\beta_{o\to o}$*,* $\beta_{i\to o}$*, and* $\beta_{o\to i}$*; the decay exponent,* $D_F$*; the asymmetry score, A; and the cell loss score, L. For* $\alpha$*, black indicates smaller values and stronger adhesion, and green indicates larger values and weaker adhesion. For* $\beta$*, red indicates attraction, blue indicates repulsion, and black indicates no long-range interaction. For A and L, orange indicates larger values. Outer-to-outer attraction,* $\beta_{o\to o} > 0$*, is strongly enriched among the top-scoring simulations.*

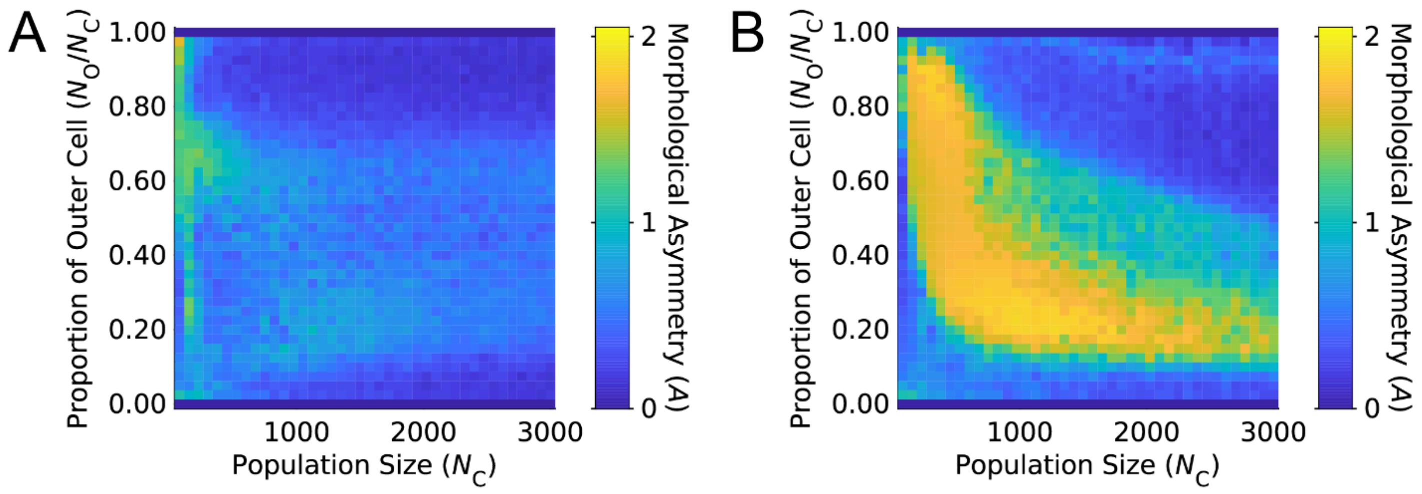


***Figure S16. Outer-to-outer attraction promotes axis formation across cell numbers and outer-cell fractions.***

*Morphological asymmetry score, A, across simulations with different total cell numbers,* $N_C = 75{:}75{:}3000$*, and different outer-cell fractions,* $\frac{N_O}{N_C} = 0{:}0.025{:}1$*.*

*(A) Simulations without long-range attraction, using the best adhesion-only parameter set:* $\alpha_{i-i} = 0.750$*,* $\alpha_{o-o} = 0.725$*,* $\alpha_{i-o} = 0.800$*, and* $\beta_{o\to o} = 0$*.*

*(B) Simulations with outer-to-outer long-range attraction, using* $\alpha_{i-i} = 0.775$*,* $\alpha_{o-o} = 0.950$*,* $\alpha_{i-o} = 0.875$*, and* $\beta_{o\to o} = 0.135$*.*

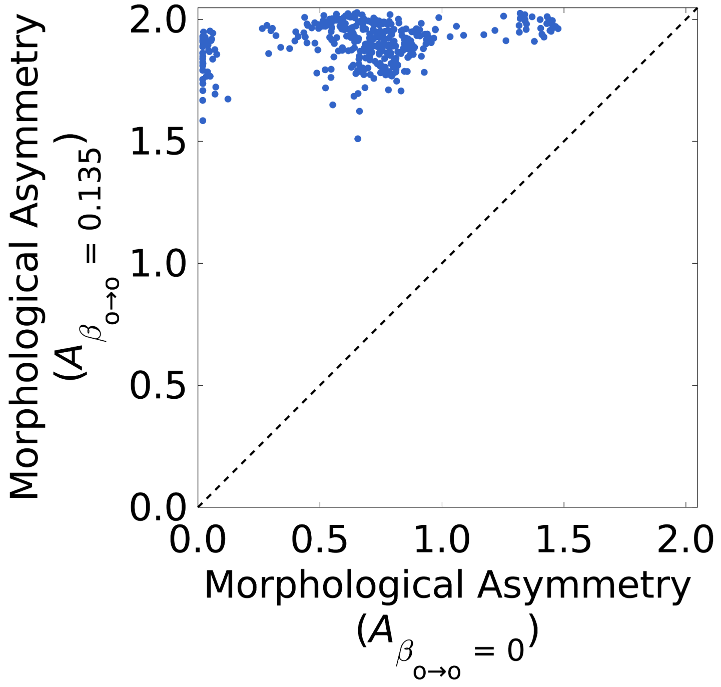


***Figure S17. Outer-to-outer attraction increases*** *A* ***across time steps and noise levels.***

*Comparison of simulations without and with outer-to-outer long-range attraction across time step lengths,* $\Delta T = 0.02{:}\,0.01{:}\,0.2$*, and noise levels,* $\kappa_F = 0.01{:}\,0.01{:}\,0.19$*. Each point corresponds to one* $(\Delta T, \kappa_F)$ *parameter pair. The horizontal axis shows A for simulations without long-range attraction, using* $\alpha_{i-i} = 0.750$*,* $\alpha_{o-o} = 0.725$*,* $\alpha_{i-o} = 0.800$*, and* $\beta_{o\to o} = 0$*. The vertical axis shows A for simulations with outer-to-outer attraction, using* $\alpha_{i-i} = 0.775$*,* $\alpha_{o-o} = 0.950$*,* $\alpha_{i-o} = 0.875$*, and* $\beta_{o\to o} = 0.135$*. Points above the diagonal indicate conditions in which outer-to-outer attraction increases A.*

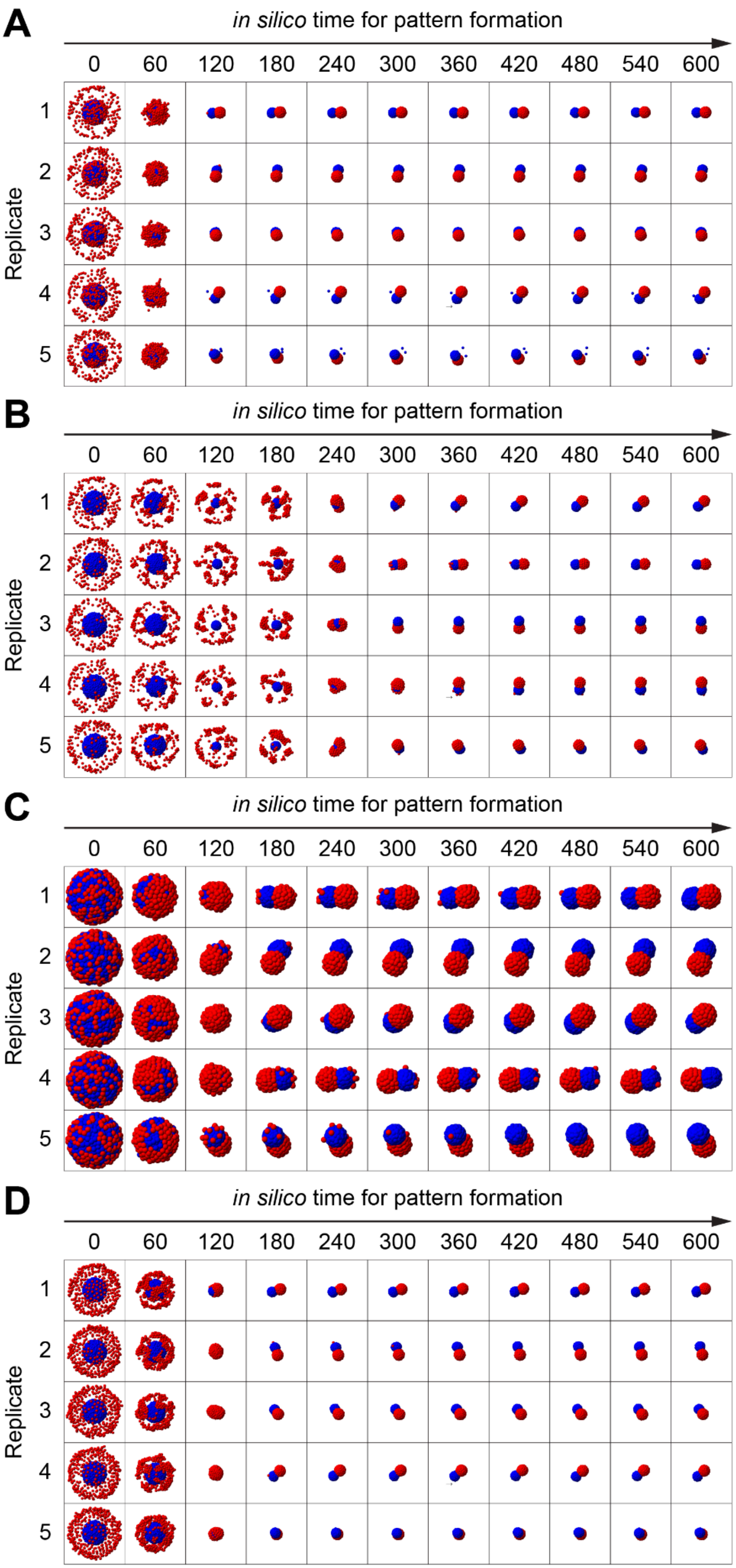
A
in silico time for pattern formation
0 60 120 180 240 300 360 420 480 540 600
Replicate
1 2 3 4 5
B
in silico time for pattern formation
0 60 120 180 240 300 360 420 480 540 600
Replicate
1 2 3 4 5
C
in silico time for pattern formation
0 60 120 180 240 300 360 420 480 540 600
Replicate
1 2 3 4 5
D
in silico time for pattern formation
0 60 120 180 240 300 360 420 480 540 600
Replicate
1 2 3 4 5

***Figure S18. Axis formation does not require a perfect initial outer shell.***

*Time courses of simulations in which the initial positions of outer cells were perturbed before the simulation began. Simulations used* $\alpha_{i-i} = 0.775$, $\alpha_{o-o} = 0.950$, $\alpha_{i-o} = 0.875$, *and* $\beta_{o \to o} = 0.135$. *In each case, displaced outer cells were moved outward by doubling their distance from the aggregate centroid.*

*(A) Half of the outer cells were moved beyond the periphery.*

*(B) Half of the outer cells were moved beyond the periphery, and the outer cells that remained in the original shell were removed.*

*(C) Half of the outer cells were moved beyond the periphery, and the displaced outer cells were removed.*

*(D) All outer cells were moved beyond the periphery.*

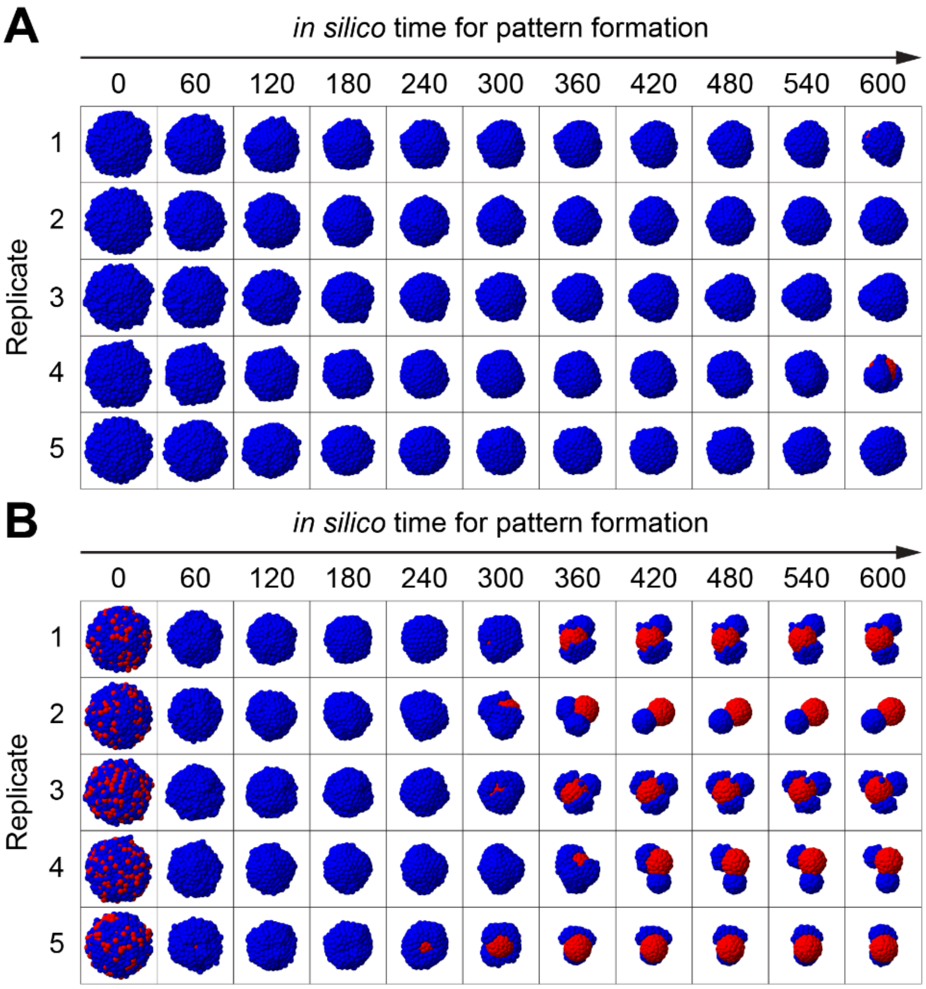


***Figure S19. Swapping or randomizing initial cell-type positions disrupts one-axis formation.***

*Time courses of simulations in which the initial positions of inner and outer cells were perturbed before the simulation began. Simulations used* $\alpha_{i-i} = 0.775$, $\alpha_{o-o} = 0.950$, $\alpha_{i-o} = 0.875$, *and* $\beta_{o \to o} = 0.135$. *In these simulations, long-range attraction remains assigned to the original outer-cell type. Swapping or randomizing cell positions reduces the alignment between peripheral position and long-range attraction and makes one-axis formation less reliable.*

*(A) Initial positions of inner and outer cell types were swapped.*

*(B) Initial positions of inner and outer cell types were randomized.*

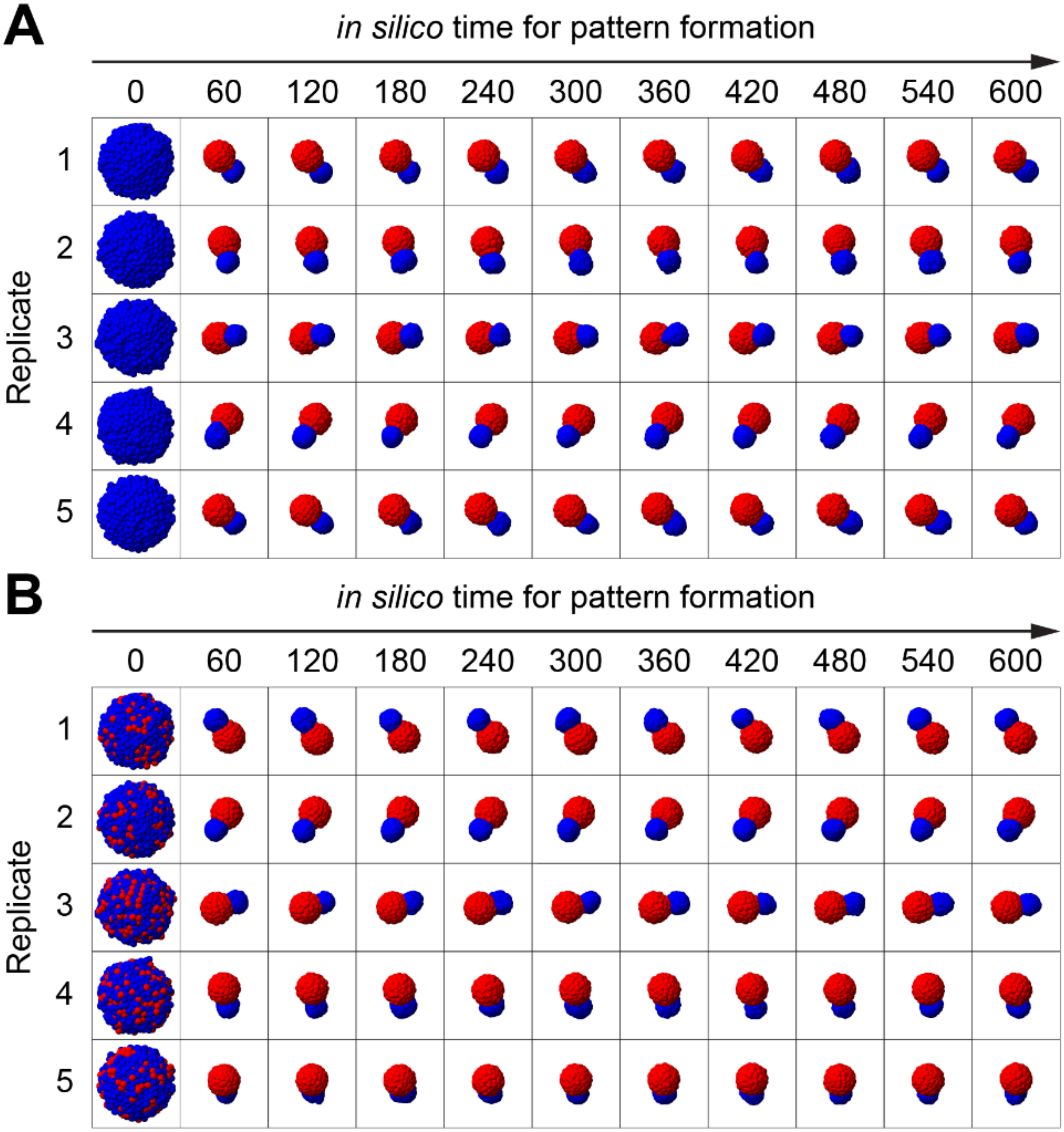


***Figure S20. Assigning long-range attraction to peripheral-positioned cells restores one-axis formation.***

*Time courses of simulations in which the initial positions of inner and outer cells were swapped or randomized, and long-range attraction was also assigned to the cells positioned at the periphery. Simulations used* $\alpha_{i-i} = 0.775$, $\alpha_{o-o} = 0.950$, $\alpha_{i-o} = 0.875$, $\beta_{o\to o} = 0.135$, *and* $\beta_{i\to i} = 0.135$. *This restores the alignment between peripheral position and long-range attraction and allows the aggregate to form one axis.*

*(A) Initial positions of inner and outer cell types were swapped.*

*(B) Initial positions of inner and outer cell types were randomized.*

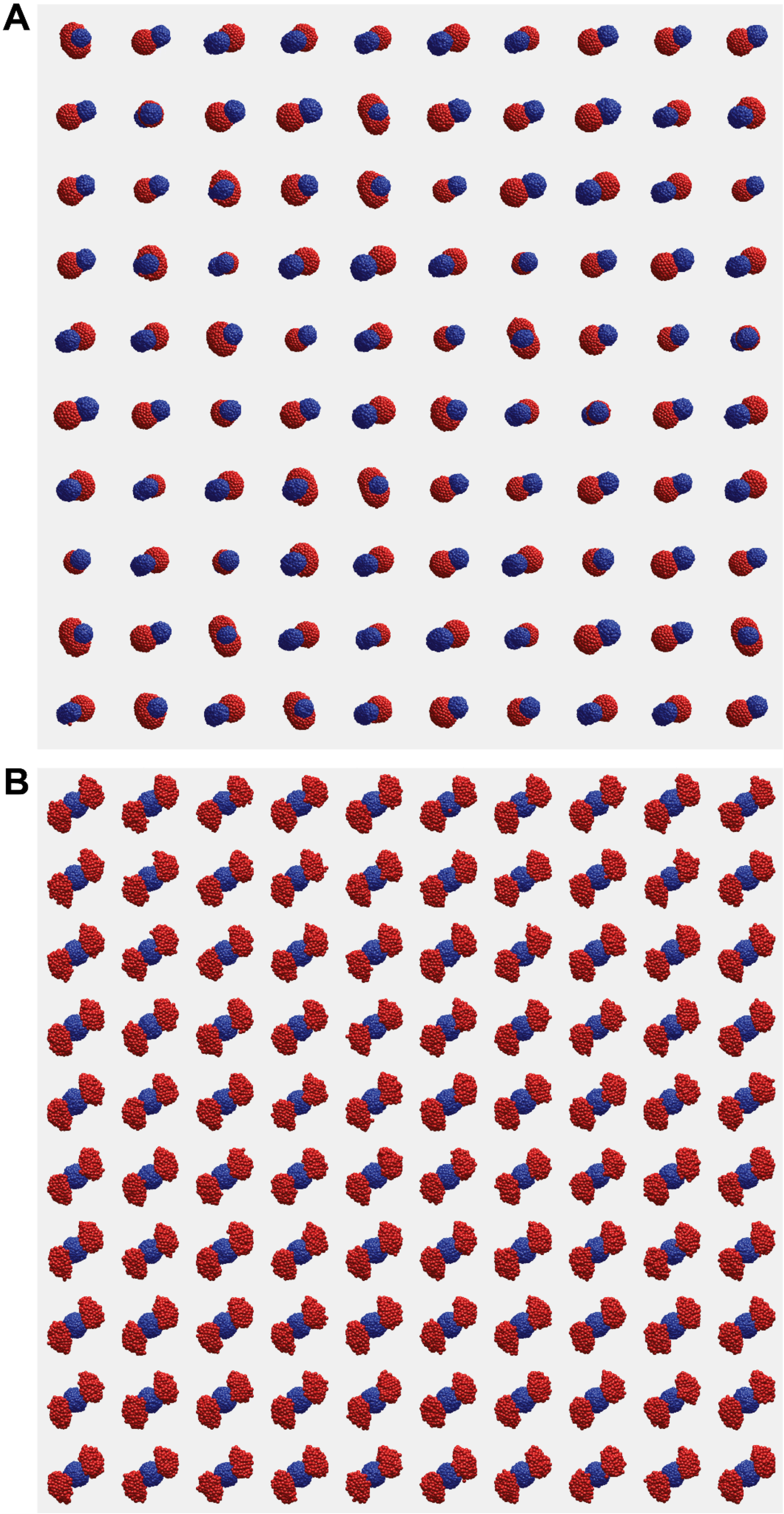

***Figure S21. Long-range attraction promotes single-pole resolution in opposite-orientation merging simulations.***

*Final states from* 100 *independent simulations of two polarized model gastruloids initialized in opposite A-P orientations.*

*(A) Simulations with outer-to-outer long-range attraction. Most pairs resolved the two posterior domains into one continuous posterior domain.*

*(B) Simulations without long-range attraction. The two posterior domains remained separated at opposite ends of the fused aggregate.*

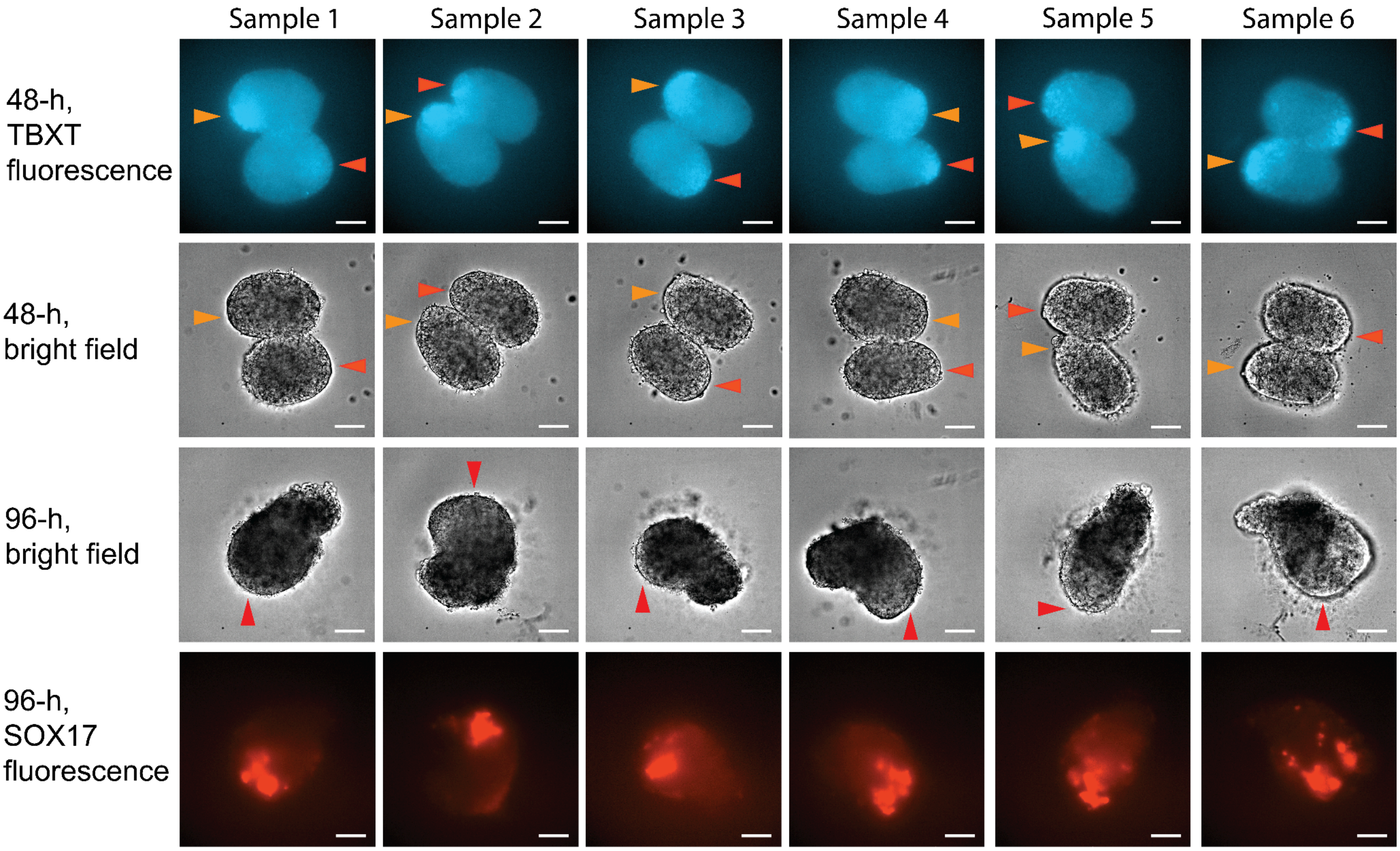


***Figure S22. Merging gastruloids form one posterior domain under reduced Chir.***

*Six independent merging experiments under the reduced CHIR dose. Rows show, from top to bottom: TBXT/BRA fluorescence at 48 h, brightfield at 48 h, brightfield at 96 h, and SOX17 fluorescence at 96 h. TBXT/BRA fluorescence marks the two initial posterior poles before merging. SOX17 fluorescence marks the final posterior-associated domain after merging. In all six samples, the fused aggregate formed one final posterior domain. Scale bars, 100 µm.*

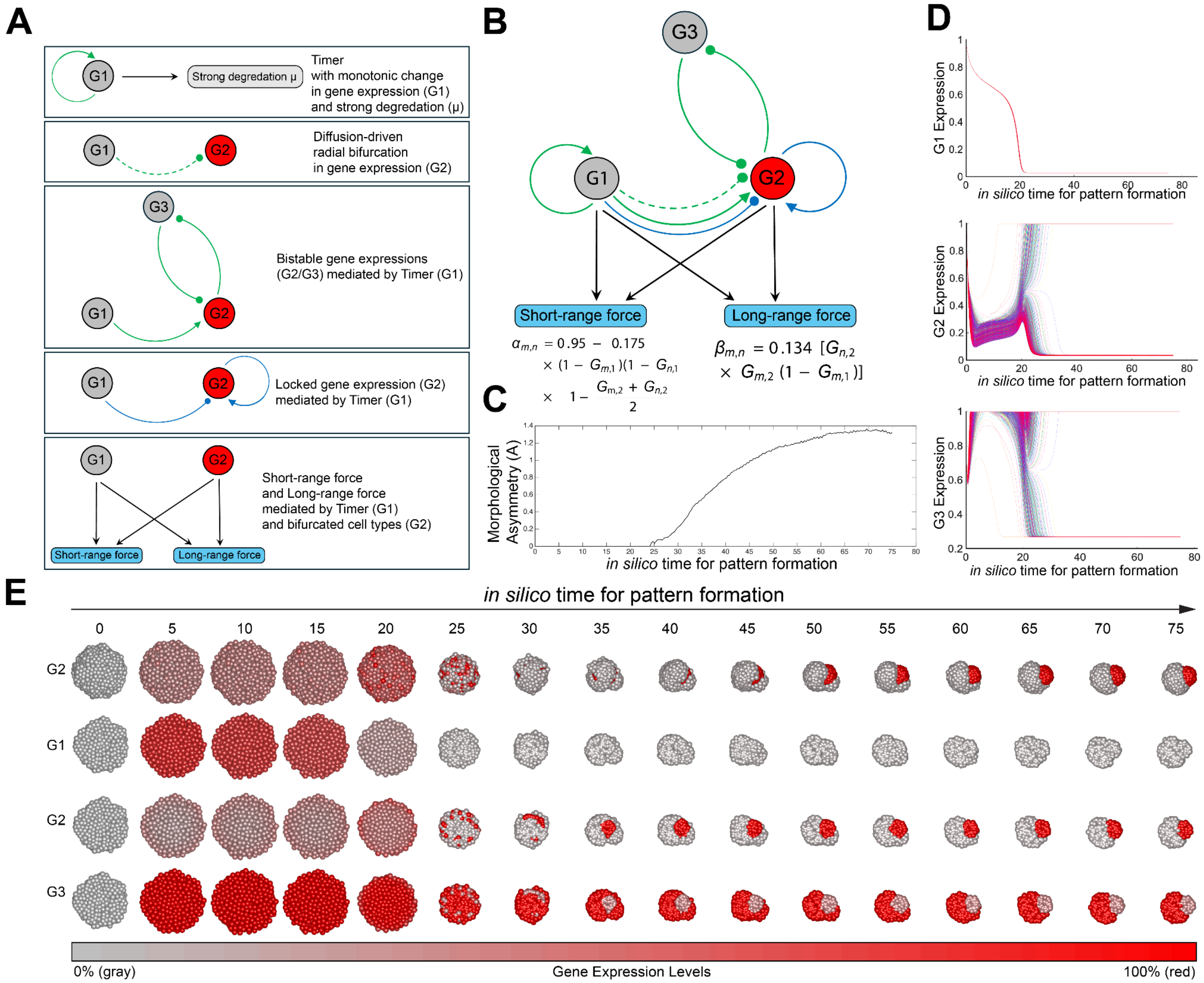


***Figure S23. An alternative gene-mechanical simulation produces core invasion and convergence.***

*(A) Design logic of the three-gene circuit. G1 acts as a timer. It starts high and then decreases, creating an early establishment stage and a later maintenance stage. During the establishment stage, signaling from G1 creates a radial difference in G2. G2 and G3 then form a mutually inhibitory switch that separates cells into two gene-expression states. During the maintenance stage, a lock-in module preserves these states [Wolpert. J. Theor. Biol. 1969]. The final gene state controls short-range adhesion and long-range attraction.*

*(B) Schematic of the full gene-mechanical network. Each cell contains the same three-gene circuit. Gene regulation includes intracellular interactions and distance-dependent intercellular signaling. Mechanical parameters are read out from gene state. One readout controls the short-range adhesion parameter. A second readout controls the long-range attraction parameter. In this abstract circuit, the G2/G3 switch can be viewed as a Wnt/Nodal-like opposition, but the genes are not assigned to specific molecular pathways.*

*(C) Morphological asymmetry score, A, over time. A increases as the gene circuit activates the mechanical interactions and the aggregate forms a polarized structure.*

*(D) Gene-expression trajectories over time. G1 decreases as the timer turns off. G2 and G3 then bifurcate into two stable expression states. Each trace shows one simulated cell. Colors distinguish cells and do not encode additional variables.*

*(E) Time course of pattern formation from an initially uniform spherical aggregate. Red indicates high gene expression, and gray indicates low gene expression. Row 1 shows the whole aggregate colored by G2 expression. Rows 2–4 show cross-sections colored by G1, G2, and G3 expression. In this simulation, the circuit first generates radial gene-expression differences. It then activates mechanical interactions that cause one population to move inward, invade the core, and converge. This produces a polarized final structure without pre-assigned cell types or an imposed A-P direction.*

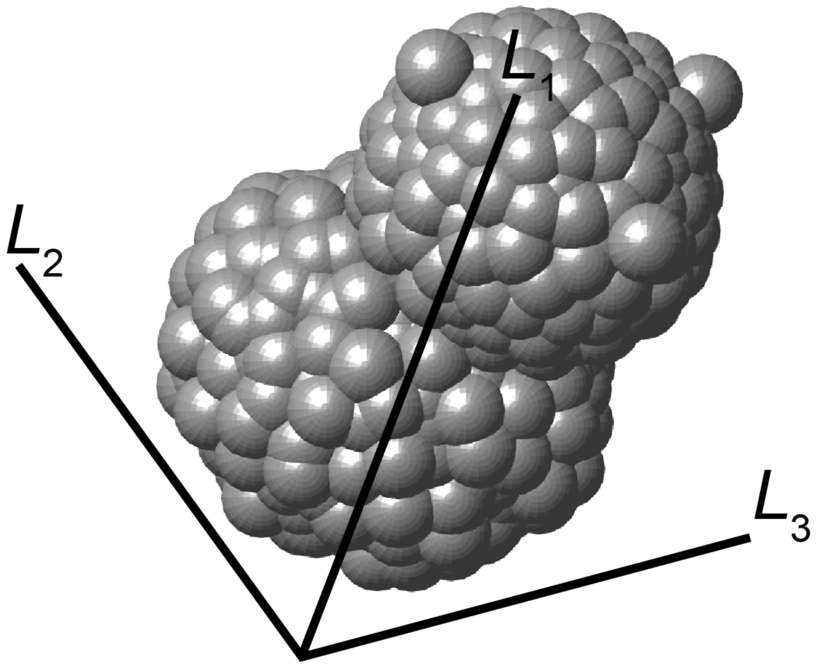


***Figure S24. Defining principal axes of a simulated 3D aggregate.***

*Schematic showing the three orthogonal principal axes of a DevSim-generated 3D aggregate. The axis directions were determined by principal component analysis of the cell-center positions. $L_1$, $L_2$, and $L_3$ denote the longest, intermediate, and shortest maximum-to-minimum extents of the projected coordinates, respectively. The illustrated axes are schematic and are not drawn to scale [Guan et al. Membranes 2024].*

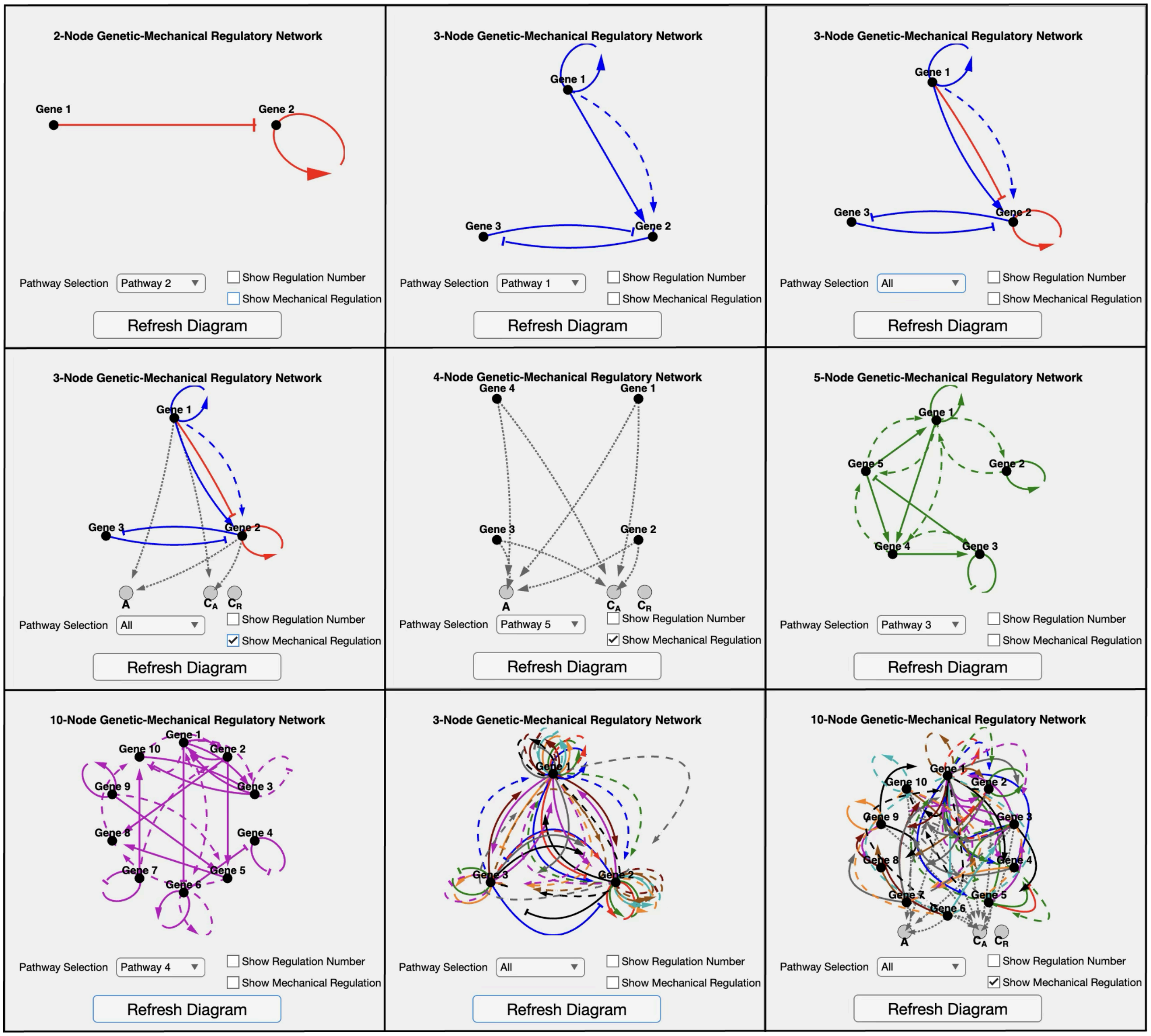


***Figure S25. DevSim visualizes arbitrary multi-node gene-mechanical networks.***

*Examples of multi-node gene-mechanical regulatory networks visualized in DevSim. The diagrams illustrate that the platform can display user-defined numbers of genes, parallel pathways, regulatory edges, and regulatory types.*

#### Supplemental Table

***Table S1. Simulated morphogenetic landscapes across short- and long-range interaction parameters.***

*Each row is one simulated parameter setting. Columns list the three short-range adhesion parameters,* $\alpha_{i-i}$*,* $\alpha_{o-o}$*, and* $\alpha_{i-o}$*; the four directed long-range interaction parameters,* $\beta_{i\to i}$*,* $\beta_{o\to o}$*,* $\beta_{i\to o}$*, and* $\beta_{o\to i}$*; the long-range distance-decay exponent,* $D_F$*; and the resulting morphological asymmetry score, A, and cell loss score, L. For A and L, the table reports the average across five independent simulations and the value from each replicate.*

***Table S2. Shape descriptors implemented in DevSim.***

*The table lists the 3D shape descriptors used to quantify simulated aggregates, their mathematical definitions, and the original references.* $L_1$*,* $L_2$*, and* $L_3$ *denote the longest, intermediate, and shortest principal lengths of the object, respectively. V is object volume, S is object surface area,* $S_{convex}$ *is the surface area of the 3D convex hull, and* $A_{convex}$ *is the projected area of the convex hull. Descriptors and notation are adapted from [Guan et al. Membranes 2024].*

#### Supplemental Movie

***Movie S1. Time-lapse brightfield imaging of human gastruloids with or without CHIR99021 treatment.***

*Time-lapse brightfield recording of four human gastruloids from the same batch. Gastruloids were cultured with or without CHIR99021 treatment from 0 to 24 h after seeding.*

***Movie S2. Adhesion-only simulations do not reliably form one axis.***

*Time course of five representative adhesion-only simulations starting from a spherical aggregate. Simulations used* $\alpha_{i-i} = 0.750$*,* $\alpha_{o-o} = 0.725$*,* $\alpha_{i-o} = 0.800$*, and* $\beta_{o\to o} = 0$*. The simulations show weak separation, local clustering, or variable final structures rather than one stable axis. Outer cells are shown in red. Inner cells are shown in blue.*

***Movie S3. Outer-to-outer long-range attraction converts radial patterning into one axis.***

*Time course of five representative simulations starting from a spherical aggregate. Simulations used* $\alpha_{i-i} = 0.775$*,* $\alpha_{o-o} = 0.950$*,* $\alpha_{i-o} = 0.875$*, and* $\beta_{o\to o} = 0.135$*. With outer-to-outer long-range attraction, the outer cells collect into one pole and the inner cells form the opposite domain. Outer cells are shown in red. Inner cells are shown in blue.*

***Movie S4. The same adhesion parameters do not form one stable axis without long-range attraction.***

*Time course of five representative simulations starting from a spherical aggregate. Simulations used* $\alpha_{i-i} = 0.775$, $\alpha_{o-o} = 0.950$, $\alpha_{i-o} = 0.875$, *and* $\beta_{o\to o} = 0.135$. *Without outer-to-outer long-range attraction, the simulations show weak separation or local clustering rather than one stable axis. Outer cells are shown in red. Inner cells are shown in blue.*

***Movie S5. Same-orientation merging simulation without long-range attraction.***

*Time course of two polarized model gastruloids initialized in the same A-P orientation and simulated without long-range attraction. The two domains merge rapidly. Red and blue mark the two pre-patterned cell populations.*

***Movie S6. Same-orientation merging simulation with long-range attraction.***

*Time course of two polarized model gastruloids initialized in the same A-P orientation and simulated with outer-to-outer long-range attraction. The two domains merge rapidly. Red and blue mark the two pre-patterned cell populations.*

***Movie S7. Opposite-orientation merging simulation without long-range attraction.***

*Time course of two polarized model gastruloids initialized in opposite A-P orientations and simulated without long-range attraction. The anterior domains merge, but the two posterior domains remain separated at opposite ends of the fused aggregate. Red and blue mark the two pre-patterned cell populations.*

***Movie S8. Opposite-orientation merging simulation with long-range attraction.***

*Time course of two polarized model gastruloids initialized in opposite A-P orientations and simulated with outer-to-outer long-range attraction. The two posterior domains move around the fused aggregate and converge into one domain. Red and blue mark the two pre-patterned cell populations.*

***Movie S9. Same-orientation human gastruloid merging experiment.***

*Time-lapse brightfield and fluorescence imaging of two human gastruloids cultured under the standard CHIR99021 dose and initially aligned in the same orientation. CHIR99021 was applied from 0 to 24 h after seeding. One gastruloid was transferred into a well containing another gastruloid at 48 h after seeding.*

***Movie S10. Opposite-orientation human gastruloid merging experiment.***

*Time-lapse brightfield and fluorescence imaging of two human gastruloids cultured under the standard CHIR99021 dose and initially aligned in opposite orientations. CHIR99021 was applied from 0 to 24 h after seeding. One gastruloid was transferred into a well containing another gastruloid at 48 h after seeding. The posterior domains move around the fused aggregate and approach each other.*

***Movie S11. Human gastruloid merging under the standard CHIR99021 dose.***

*Time-lapse brightfield and fluorescence imaging of human gastruloid merging experiments under the standard CHIR99021 dose. CHIR99021 was applied from 0 to 24 h after seeding. One gastruloid was transferred into a well containing another gastruloid at 48 h after seeding. Three single gastruloids and six merging gastruloid pairs are shown for comparison.*

***Movie S12. Human gastruloid merging under the reduced CHIR99021 dose.***
*Time-lapse brightfield and fluorescence imaging of human gastruloid merging experiments under the reduced CHIR99021 dose. CHIR99021 was applied from 0 to 24 h after seeding. One gastruloid was transferred into a well containing another gastruloid at 48 h after seeding. Three single gastruloids and six merging gastruloid pairs are shown for comparison.*

***Movie S13. Gene-mechanical simulation in which outer cells move along the surface and converge.***
*Time-lapse simulation of the three-gene gene-mechanical circuit. The circuit first generates radial gene-expression differences. It then activates mechanical interactions that move the outer population along the surface and into one pole.*

***Movie S14. Alternative gene-mechanical simulation with core invasion and convergence.***
*Time-lapse simulation of an alternative gene-mechanical circuit. The circuit generates radial gene-expression differences and then activates mechanical interactions that cause one population to move inward, invade the core, and converge.*

***Movie S15. Step-by-step instruction video for DevSim.***
*Step-by-step instruction video for using the DevSim platform. The video is available on YouTube: https://youtu.be/BlEGs4XDDkY?si=zeIiclUJm0RLaQJ1.*

***Movie S16. DevSim simulation of a layered pattern.***
*DevSim-exported movie showing a gene-mechanical simulation that forms a layered multicellular pattern.*

***Movie S17. DevSim simulation of a bilobed pattern.***
*DevSim-exported movie showing a gene-mechanical simulation that forms a bilobed multicellular pattern.*

***Movie S18. DevSim simulation of a multilobed pattern.***
*DevSim-exported movie showing a gene-mechanical simulation that forms a multilobed multicellular pattern.*

***Movie S19. DevSim simulation of a striped pattern.***
*DevSim-exported movie showing a gene-mechanical simulation that forms a striped multicellular pattern.*

#### Supplemental Text 1

##### *DevSim* User Guidebook

##### Purpose

*DevSim* is a *MATLAB* app for simulating multicellular development with coarse-grained cells moving in an overdamped medium, gene circuits governed by Hill functions, and long- and short-range cell-to-cell interactions. It is designed to help users explore a range of developmental patterns (e.g., symmetry breaking, aggregation, peripheral patterning), visualize outcomes, and export measurements (gene expression over time, 3D shape descriptions).

**Availability and Citation**

**Software name:** *DevSim*

**Distribution:** https://github.com/hormoz-lab/Symmetry-Breaking/tree/Main/DevSim

**How to cite :** "DevSim (Developmental-Simulator), version 1.0. MATLAB app and example templates. https://github.com/hormoz-lab/Symmetry-Breaking/tree/Main/DevSim"

**Contact:** &

**License:** MIT license.

**System Requirements**

**MATLAB:** created/tested on **R2024b**; should run on recent releases.

**Toolboxes:** Parallel Computing Toolbox (for parfor).

**OS:** Windows & macOS tested (Linux not yet tested).

**Excel editing:** The "Edit …" buttons attempt to open Microsoft Excel. If Excel isn't present, open the .xlsx files manually in the user's editor of choice.

**Hardware:** No GPU required. With the default settings (~1,350 cells; 8 sims; dt=0.2; Tmax=50), an M-series MacBook Pro (M4) completes in ~2 minutes. Any modern Intel i5 or better, or any Apple-silicon (M-series) Mac is fine

**Folder layout and files inside DevSim**

```
DevSim/
    DevSim_GUI.m
    RunSimulation.m
    Active/
        UserParams.xlsx
        Network Settings.xlsx
    Templates/
        Symmetry_Breaking_Template/
            UserParams.xlsx
            Network Settings.xlsx
        Poly-Lobed_Template/
            UserParams.xlsx
            Network Settings.xlsx
        …
    OUTPUT1/
    OUTPUT2/
```

…

**Template Pack**

A **template pack** is simply a folder containing the two .xlsx files above (UserParams.xlsx and Network Settings.xlsx). Use the "Export Template" button in the *DevSim* GUI to create one; use the "Load Template" button to copy one into Active/. A Default Symmetry-Breaking Template Pack will be provided.

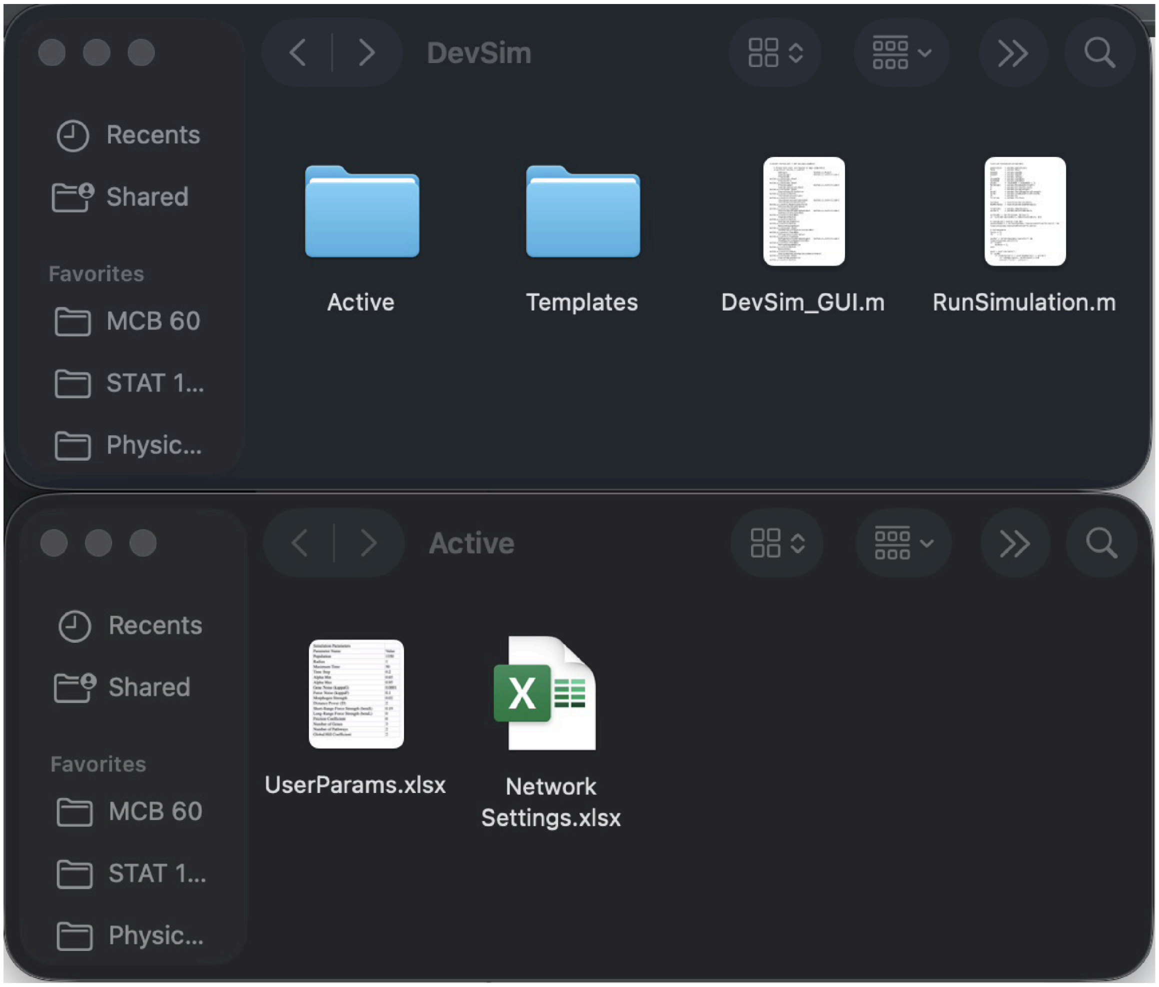


***Figure G1.*** *DevSim/ directory (top) and Active/ directory (bottom).*

<u>**Step-by-step Tutorial**</u>

**(A) Launch *DevSim***

1. Place the DevSim folder somewhere convenient. Ensure all components exist: DevSim/Active/; DevSim/DevSim_GUI.m; DevSim/RunSimulation.m; DevSim/Templates/.

2. Open MATLAB version: R2024b (older versions may not be able to support *DevSim*). In MATLAB, set the Current Folder to the DevSim root.

3. Run DevSim_GUI to open the app.

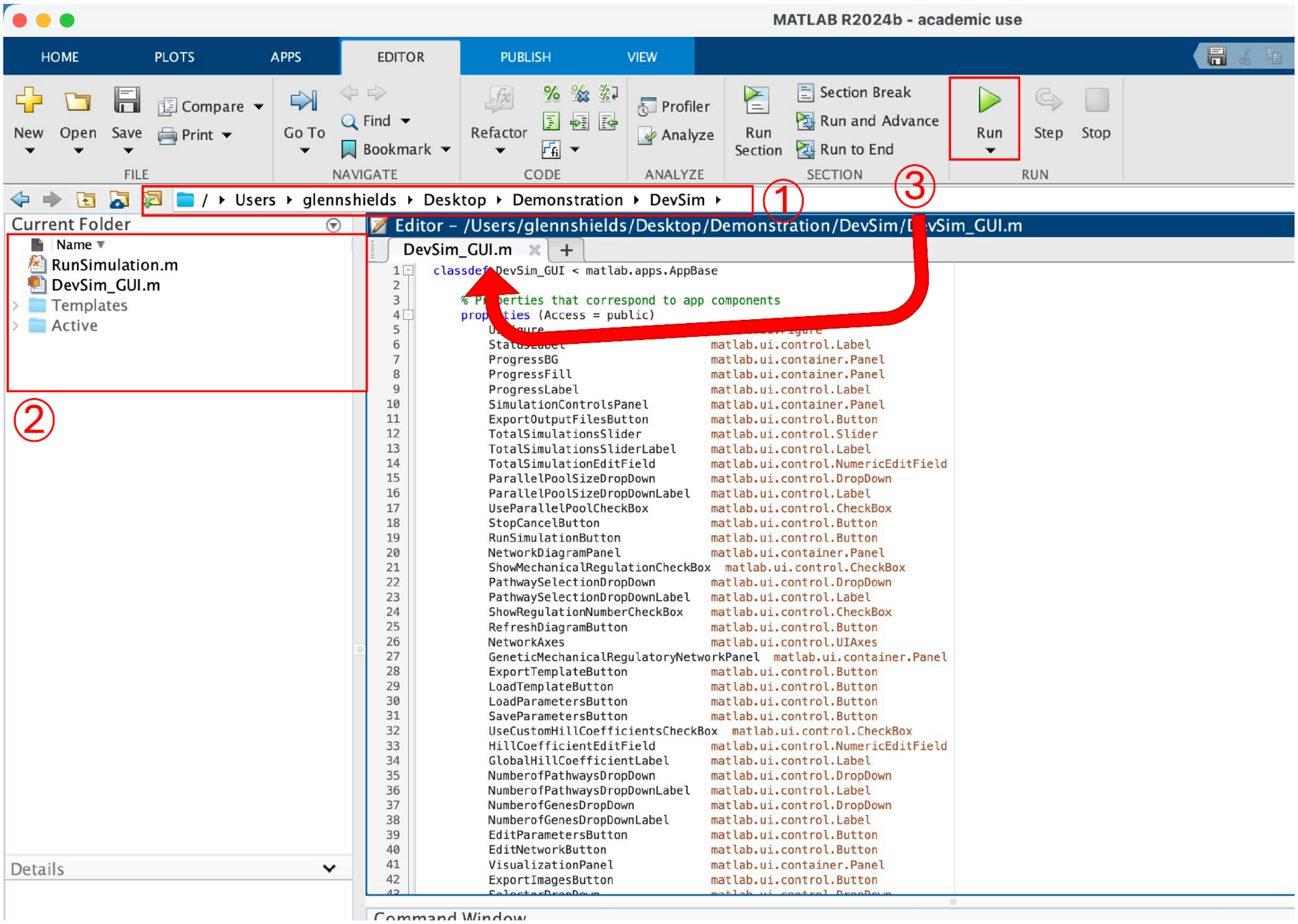


***Figure G2.*** *Launching DevSim (steps for section A). ① Open MATLAB R2024b (older versions may not support DevSim). In MATLAB, set the Current Folder to the DevSim root. ② Ensure all folders are properly nested, double-click on "DevSim_GUI.m" to load the script into the MATLAB window. ③ Make sure the open script is "DevSim_GUI.m" and press Run to open the DevSim GUI.*

**(B) Start from known default parameters**

1. When opened, *DevSim* loads default parameters. At any time, the user can press "Reset Parameters" to reset to the default parameters in the *DevSim* Graphical User Interface (GUI); pressing "Reset Parameters" will not affect the Active/ folder or the Active Excel workbooks. The default parameters in the GUI are as follows:

Population Size = 1350; Radius = 1; Maximum Time = 50; Time Step (dt) = 0.2; Alpha Min = 0.65; Alpha Max = 0.95; Gene Noise (kappaG) = 0.0001; Force Noise (kappaF) = 0.1; Morphogen Strength = 0.0185; Distance Power (D) = 2; Short-Range Force Strength (betaS) = 0.175; Long-Range Force Strength (betaL) = 0.125; Friction Coefficient = 0; Number of Genes = 3; Number of Pathways = 2; Global Hill Coefficient = 2.

Note: Parameters have minimal bounds to maximize flexibility; it is recommended to start from the defaults before exploring the large parameter space.

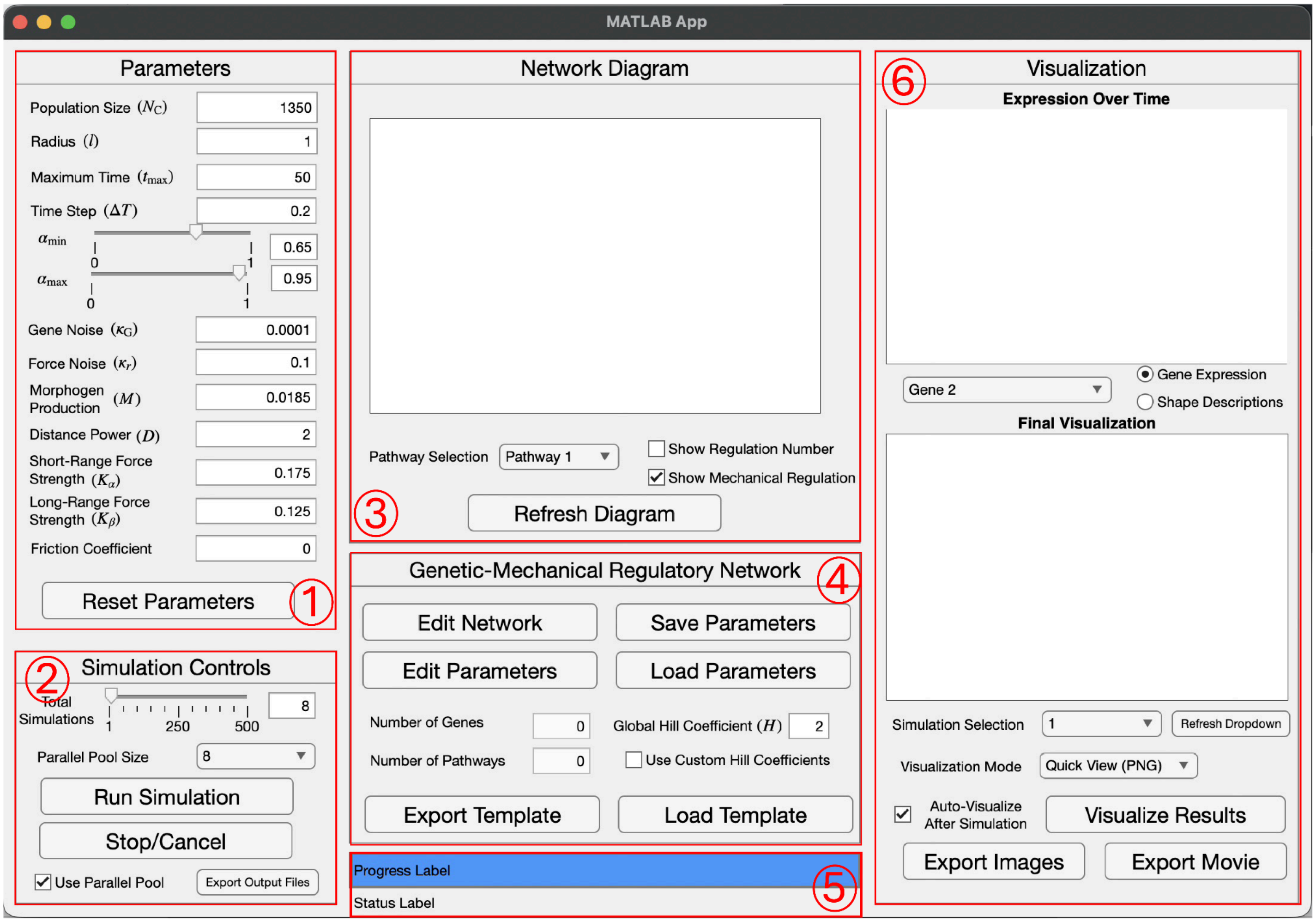


***Figure G3.*** *DevSim GUI with panels and important features highlighted in red and numbered. ① Parameters panel. ② Simulation Controls panel. ③ Network Diagram panel ④ Genetic-Mechanical Regulatory Network Pane (GMRN). ⑤ Progress Bar, Progress Label, and Status Label. ⑥ Visualization panel.*

**(C) Decide how many simulations and whether to use parallel**

In the Simulation Controls panel:

1. Set Total Simulations (1-500)
2. Toggle Use Parallel Pool and choose Pool Size (1-8) if the user wants to use simultaneous runs

**Warning (MATLAB Online users):** MATLAB Online doesn't support parallel pools - uncheck Use Parallel Pool box. Otherwise *DevSim* is fully compatible with the online version of MATLAB.

**Canceling:** "Stop/Cancel" button halts the simulation after the current batch; it does not interrupt an active parfor iteration. To stop immediately during a parfor, the user may need to close the app/MATLAB directly.

**(D) Inspect or set the size of the Genetic-Mechanical Regulatory Network (GMRN)**

1. Inspect Number of Genes and Number of Pathways in Genetic-Mechanical Regulatory Network panel.
2. Set Global Hill Coefficient (used everywhere unless overridden per-edge)
3. Check "Use Custom Hill Coefficients" to set per-edge Hill coefficients in the Excel workbook.

**Design Choice:** *DevSim* supports unlimited genes and unlimited regulatory pathways. Pathways are visualized in distinct colors (Pathway 1 = blue, 2 = red, 3 = green, …).

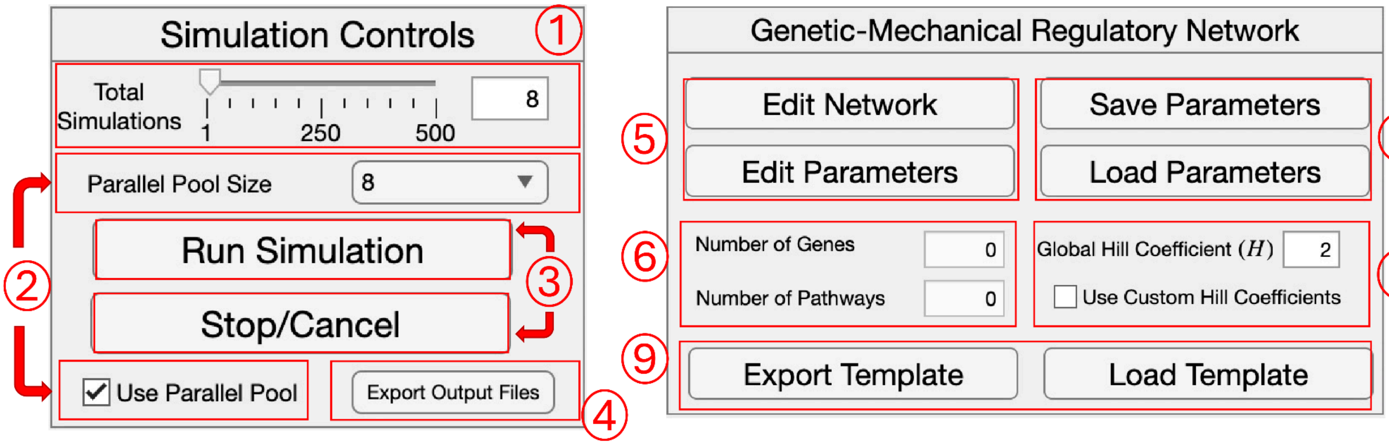


***Figure G4.*** *DevSim Simulation Controls (Left) and Genetic-Mechanical Regulatory Network (Right) panels. ① Total Simulations Slider and Edit Field. Users can enter the number of total simulations they want to run from 1 - 500. ② Parallel Pool Size Dropdown (top) and Use Parallel Pool checkbox (bottom). To use Parallel Pool, the Use Parallel Pool checkbox should be checked, and the user will pick the number of simultaneous simulations to run via MATLAB parfor in the parallel pool size dropdown from 1 - 8. If total simulations exceeds parallel pool size, the simulations will run in batches the size of the parallel pool size value. ③ Run Simulation button (top) pressed when user is satisfied with all the parameters and Genetic-Mechanical Regulatory Network and wants to run the simulation. Stop/Cancel button (bottom) pressed when user wants to stop the simulation after the current batch (will not stop simulation during a parfor loop). ④ Export Output Files button. Users can press this button to export the entire experiment into a single parent folder for archiving or sharing, they will first choose a destination directory; then when prompted, name their Parent folder. ⑤ Edit Network button (top) will open Network Settings.xlsx file via the Active folder directory when pressed. Edit Parameters button (bottom) will open UserParams.xlsx file via the Active folder directory when pressed. ⑥ Number of Genes Display (top) users can inspect the number of genes currently being read by Network Settings.xlsx file. Number of Pathways (bottom) users can inspect the number of pathways currently being read by Network Settings.xlsx file. ⑦ Save Parameters Button (top) will store the current user parameter values entered in the Parameters panel inside of the DevSim GUI and also the Number of Genes, Number of Pathways, and Global Hill coefficient value entered in the GMRN panel into the UserParams.xlsx file via the Active folder Directory when pressed. Load Parameters Button (bottom) will load the user parameter values inside the UserParams.xlsx file in the Active folder Directory into the Parameters and GMRN panels. ⑧ Global Hill Coefficient Edit Field (top) and Use Custom Hill Coefficients (bottom). When "Use Custom Hill Coefficients" is checked, DevSim will read the user entries inside of the Active/NetworkSettings.xlsx in the HillCoeffs Sheet. ⑨ Export Template button (left) when pressed will prompt users to create a name for the template folder and once entered will transfer the Active file directory's Network Settings.xlsx and UserParams.xlsx into the user named template folder under the Templates/ file directory. Load Template button (top) when pressed will prompt users to select a Template via the Templates/ file directory; once selected, that template will be loaded into DevSim via replacing the Active file directory's files with the Template's files.*

**(E) Edit the network (Excel): internal versus external regulation**

1. Click Edit Network button in the GMRN panel to open **Active/NetworkSettings.xlsx**
2. Edit the desired number of genes in pathways at the top of the Gene Information sheet
3. For each gene edit:
   - Leak: basal expression
   - Degradation: linear decay constant
   - Initial Value: starting expression in [0,1].
   - Color: RGB triplet used in gene plots

4. Open the Gene Regulatory Network (K) sheet. For each pathway p, there are two gene number by gene number (NxN) blocks:
   - Internal (Intracellular) Regulation (left block colored light tan).
   - External (Intercellular) regulation (right block colored light blue)
5. Directionality: Rows = targets (receivers), Columns = sources (senders).
6. Enter weights: 0 = no edge (can also leave cell empty), > 0 = activation, < 0 = inhibition
7. If Use Custom Hill Coefficients is checked, open the Gene Regulatory Network (H) sheet and enter values > 0 only where the user want to override the global H (leave blank to inherit the global value).
8. Open the Short-Range Force Network (α) sheet. Enter values for the Short-range matrices.

Short-range modes (per gene): -2, -1, 0, +1, +2

9. Open the Long-Range Force Network (β) sheet. Entries define how gene expression regulates long-range force responses. Rows are receiver genes, and columns are sender genes. Use +1 for activation, -1 for inhibition, and 0 or blank for no contribution. Positive long-range force strength produces attraction. Negative long-range force strength produces repulsion.

**Interpretation:** For attraction, higher sender expression increases pull toward the sender (if +1) or reduces it (if -1). For repulsion, the sign has the analogous meaning

10. Save the workbook, go back to *DevSim*, and click the "Refresh Diagram" button in the Network Diagram panel to visualize the edits

**Edge semantics (meaning of entries):**

- Weight magnitude is the regulatory strength. The sign sets activation (positive) vs inhibition (negative).
- The Hill coefficient H is the sigmoid steepness: Large H = switch-like logic; small H = more gradual logic

Each nonzero edge weight is interpreted as the signed input strength to a sigmoidal regulation term (Hill function); the Hill coefficient controls steepness, not the sign.

IMPORTANT: If the workbook is not saved before going back to *DevSim*, none of the edits made in the workbook will be applied to *DevSim* or the current simulation.

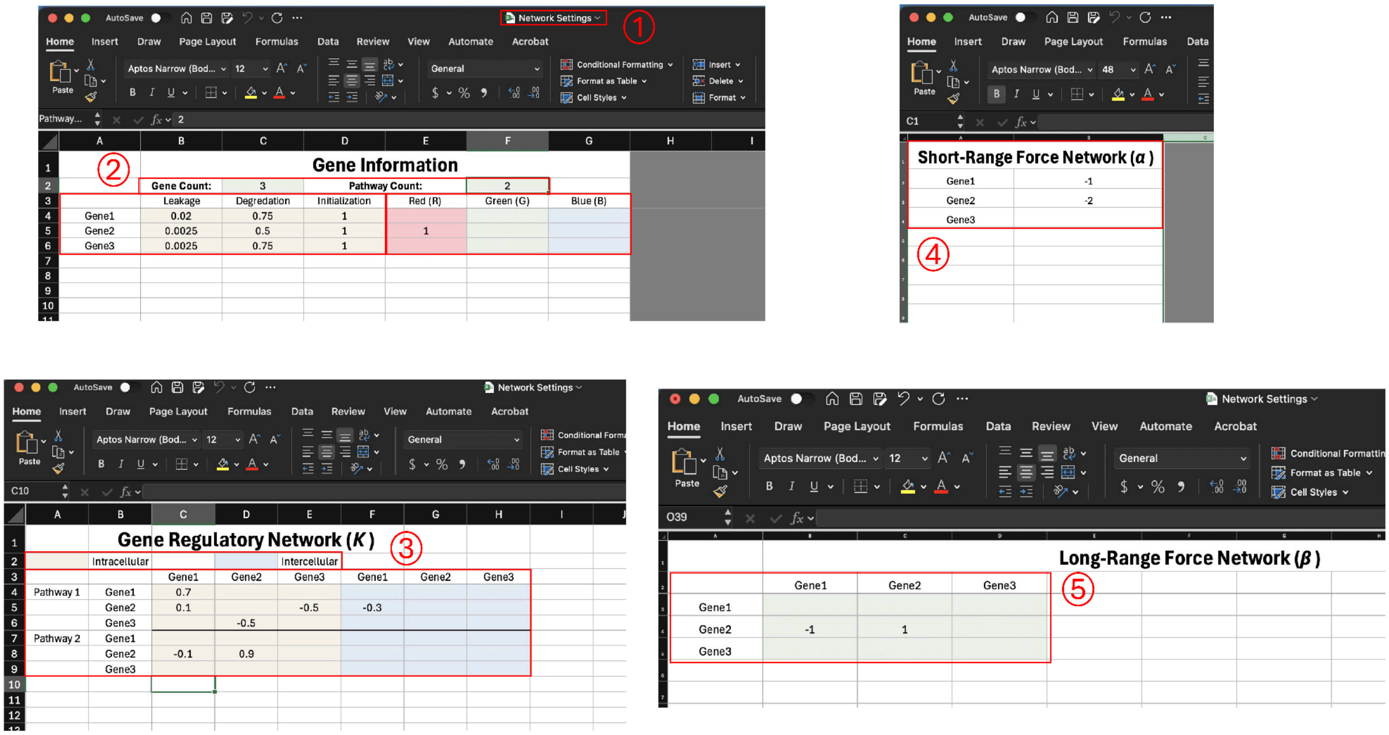


***Figure G5.*** *Network Settings.xlsx shown and can be edited by the user to change the DevSim GMRN. ① Network Settings.xlsx should open when pressing the Edit Network button found inside the DevSim GUI in the GMRN panel. ② General Gene Information can be changed in the Gene Information sheet. Top highlighted block encodes for Gene Count and Pathway Count. When these values are changed the other sheets within the Network Settings excel file will dynamically format to accommodate the new Gene and Pathway Count. Bottom left highlighted block encodes for each Gene's Leakage, Degradation and Initialization values. Bottom right highlighted block encodes for each Gene's RGB vector from 0 - 1 for visualization purposes. ③ Gene Regulatory Network (K) sheet is shown. There are two NxN block for each pathway's intracellular (internal) regulation and intercellular (external morphogen based) regulation. The blocks are color coded and multiple pathways are separated vertically and also by a solid black line. Entries in these blocks will show up inside of the DevSim GUI where solid lines are intracellular regulation, dashed lines are intercellular regulation , different colors separate the different pathways, and the rows are the target genes and the columns are the source genes. In order to add custom Hill Coefficients for GMRN connections, there is a separate Gene Regulatory Network (H) sheet with an identical layout where all the custom Hill Coefficients will be stored for their corresponding connections. ④ Short-Range Force Network (α) sheet is shown here. There are possible inputs: (-2, -1, 0, 1, 2) for the short-range force modes which can be changed for each gene. Entries in this column will be represented as gray arrows from the interacting gene(s) to A (Adhesion) to represent mechanical regulation.⑤ Long-range attraction force modes can be changed in the Long-Range Force Network (β) sheet encoding for a NxN matrix. The columns are represented as the sender gene (ligand) and the rows are represented as the receiver gene; in the DevSim Network Diagram, entries in this matrix are represented as gray arrows from the interacting genes to C_A (Chemotaxis Attraction) to represent mechanical regulation. If the Long-Range Force Strength coefficient in the DevSim GUI is <0, then the entries in this matrix are represented as gray arrows from interacting genes to C_R (Chemotaxis Repulsion) instead.*

**Optional Network Diagram viewing modes**

Toggle "Show Mechanical Regulation" in the Network Diagram panel to display gray connections to:

- A (adhesion/short-range), C_A (chemotaxis attraction), C_R (chemotaxis repulsion)
- These summarize how gene-dependent mechanical channels feed into forces – the unique logic of *DevSim*

Toggle "Show Regulation Number" in the Network Diagram panel to display, for each node (gene), incoming and outgoing counts for internal and external edges (self-loops included).

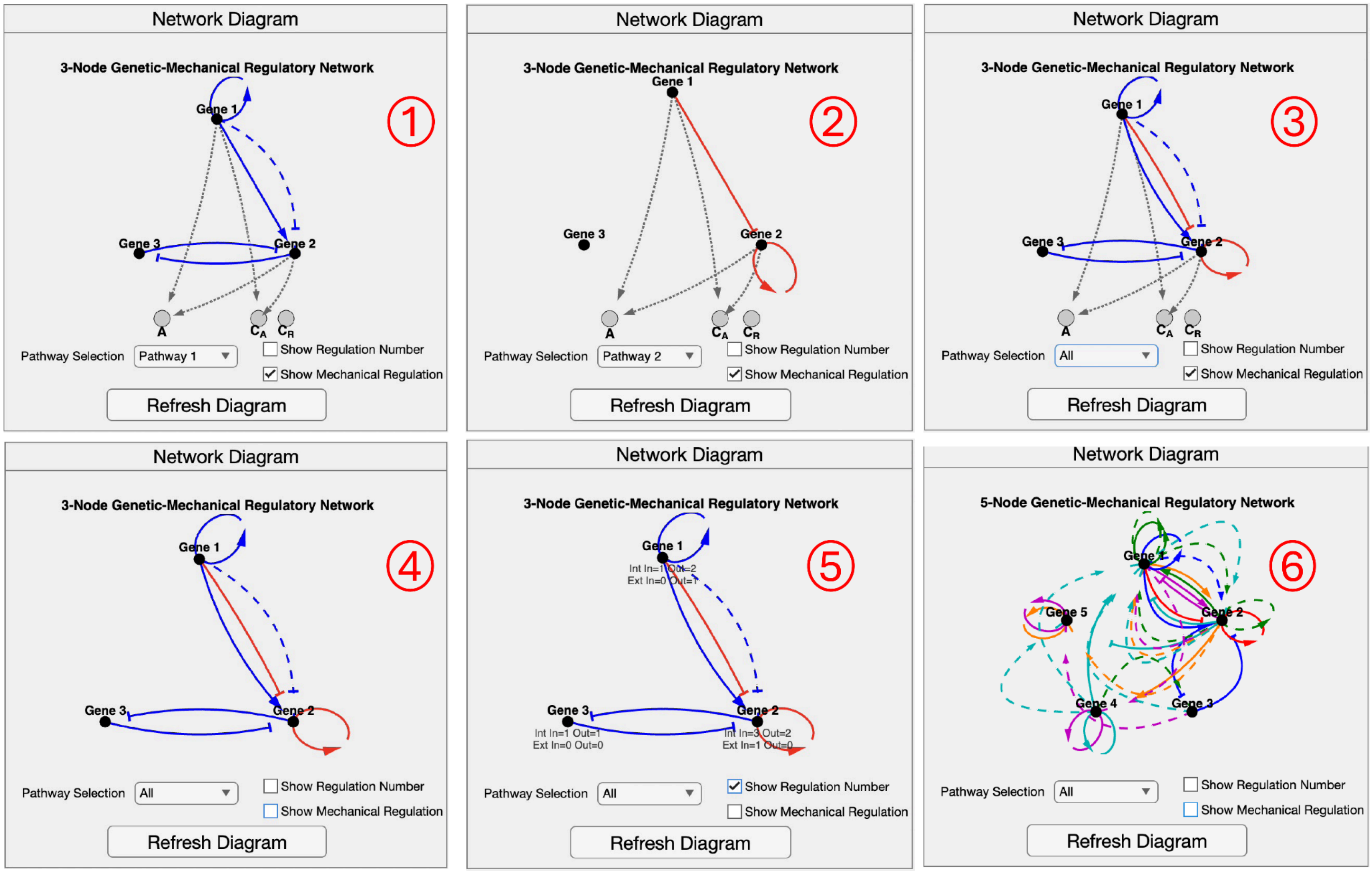


***Figure G6.*** *Six examples of DevSim GMRN diagram are shown, visualized in the Network Diagram panel in order showcase the graphing features and optional viewing modes. Single pathways or multiple pathways can be visualized: ① Default GMRN pathway 1 connections are shown by selecting Pathway 1 in the Pathway Selection Dropdown which are represented by blue connections. ② Default GMRN pathway 2 connections are shown by selecting Pathway 2 in the Pathway Selection Dropdown which are represented by red connections. ③ Default GMRN shown with both pathway 1 and pathway 2 by selecting "All" in the Pathway Selection Dropdown. ④ Gray Mechanical Regulation connections and nodes can be toggled off in the Network Diagram with the Show Mechanical Regulation checkbox ⑤ Regulation Numbers giving information about Internal and External connections entering a node and exiting a node (Gene) can be toggled on in the Network Diagram with the Show Regulation Number checkbox. ⑥ An Example of a different GMRN is given to showcase DevSim's capabilities; the example shown shows 5 genes with 6 Parallel Pathways and many connections.*

**(F) Run the Simulation**

1. Confirm all parameters are desired for simulation. Optional: press “Save Parameters” if the user wants to store the in-GUI parameters used for the current simulation in case an error occurs, the Network Settings excel file is not affected when pressing “Save Parameters” and “Load Parameters” button in the GMRN panel as the files will already be preserved if the user saved them according to the previous step E.

2. Click “Run Simulation” in the Simulation Controls panel. Doing so, *DevSim* will:

- Seed a random spherical cluster of cells.
- Delete All current OUTPUT* folders in the DevSim/ directory and create the user-defined number of new OUTPUT* folders according to how many “Total Simulations” the user selected inside of the GUI
- Create per-simulation noise in each OUTPUT folder under a “WorkSpace_Noise.mat” file
- Retrieves and unpacks the simulation parameters from the GUI and excel sheets
- Initializes the cells
- Runs the simulation via the GMRN logic, saving the cells’ initial position in the respective OUTPUT folder directory under a “WorkSpace_MatrixP_Initial.csv” file; the cells’ position and gene information at each time point in the simulation as “WorkSpace_Matrix_*.csv” files (the number of time points depends on dt and Tmax) then it saves the cells’ final position and gene information under a “WorkSpace_Matrix_Final.csv” file.
- Once complete the Status label will read “Simulation complete!”

**Live status and progress bar (simulation):** When the user clicks “Run Simulation”, the blue progress bar starts at 0% and the status reads “Starting simulation…” and then “Running Simulations…” when the simulations are in progress. The bar increases smoothly to 100% as batches complete. When all runs finish, the status changes to “Simulation complete!” and the bar is completely filled.

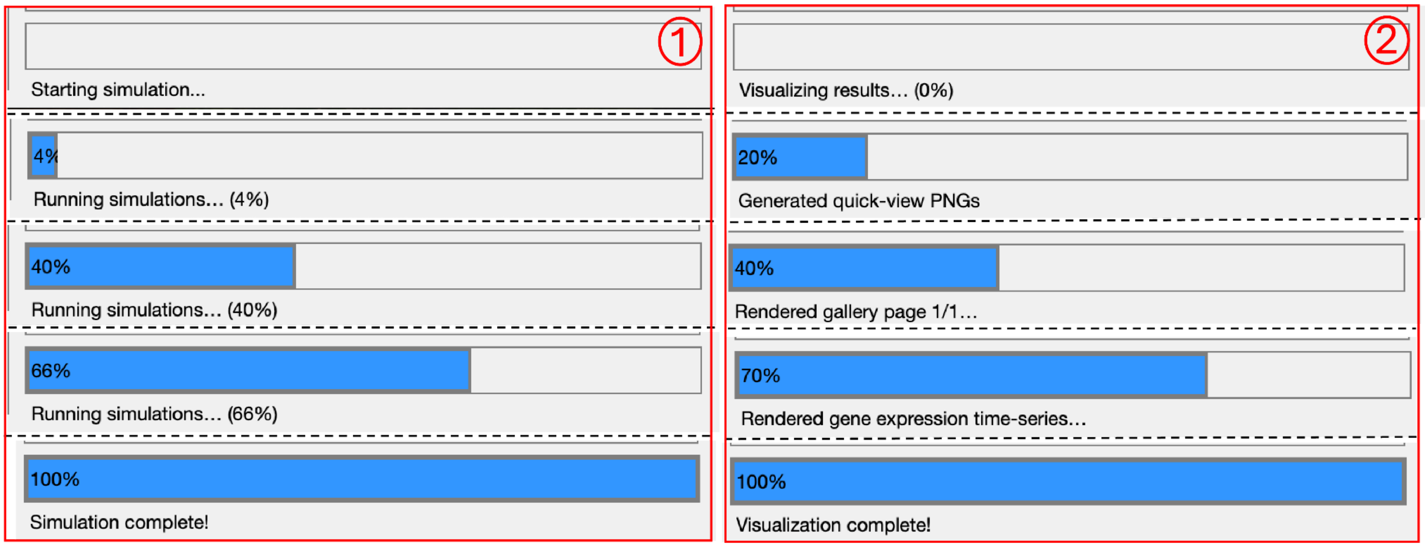


***Figure G7.*** *Run Simulation and Visualize Results Progress display. ① Column 1 shows the blue progress bar (when the user presses Run Simulation button) filling smoothly from 0% to 100% as batches complete. ② Column 2 shows the blue progress bar (when the user presses Visualize Results button) filling in stages as different visualizations complete.*

**(G) Visualize Results**

1. Once the simulation is complete and the status label reads “Simulation complete!”, click “Visualize Results” in the Visualization panel to generate the following inside of each OUTPUT directory:

- FinalSnapshot.png per simulation (for quick viewing inside of the GUI).
- GeneExpressionOverTime_Gene#.png (per-cell tracing in order to reveal bifurcations).
- ShapeOverTime_*.png for 12 shape descriptors (General Sphericity, Diameter Sphericity, Intercept Sphericity, Max. Projection Sphericity, Hayakawa Roundness, Spreading Index, Elongation Ratio, Pivotability Index, Hayakawa Flatness, Wilson Flatness, Huang Shape Factor, Corey Shape Factor).
- Gallery pages (up to 5x5 per page), will be saved in the DevSim/ directory as OUTPUT_Page#.png (multiple pages if needed).

2. Once results have been visualized and the Status label reads “Visualization complete!” inside the GUI, the first simulation should already be rendered in the GUI axes. Choose a mode in the “Visualization Mode” dropdown (Visualisation panel):

- Quick View (PNG) - fastest; pages through 3D pre-rendered snapshots of the cells.
- Interactive 3D View - renders a 3D scene the user can rotate, zoom, and perform other actions provided by MATLAB axis; allow for a short delay when selecting this view to load.
- Gallery View - shows a thumbnail grid of all 3D pre-rendered cells.

3. Use the Selector (underneath the top plot located in the visualization panel):

- In Gene Expression mode, pick Gene 1…N (shows all cells’ trajectories for that gene in one simulation).
- In Shape Descriptors mode, pick one of the 12 shape descriptions (time series for that simulation).

**Auto-Visualize:** enable “Auto-Visualize After Simulation” checkbox located in the visualization panel in order to immediately generate and render visualizations when a simulation finishes.

**Live Status and progress bar (visualization):** When “Visualize Results” is clicked, the progress bar resets to 0% and the status reads “Visualizing results…”; updates occur in steps:

- 0% → 20%: pre-render Quick View (PNG) snapshots.
- 20% → 40%: render Gallery pages
- 40% → 70%: render gene-expression time series
- 70 → 100%: render shape-descriptor time series and finalize. When finished, the status reads “Visualization complete!” and the first 3D view and graphs are automatically rendered in the Visualization panel axes.

**Export Images (optional):** Use the “Export Images” button (Visualization panel) to save all visualization PNGs into one parent folder in one step.

1. Press the Export Images button in the Visualization panel. A file viewer window will open, then choose a destination directory for the parent folder to be saved to, press Open to use the selected option.

2. Enter a parent folder name for this export when prompted by the popup window. *DevSim* creates <destination>/<ParentName>/

3. Inside the parent folder, *DevSim* writes OUTPUT* folders (each containing its respective PNGs) and the Gallery view PNG(s) at the top level.

4. When complete, check <ParentName>/OUTPUT*/ for per-run snapshots, gene-time series, and shape-description PNGs; check <ParentName>/ for gallery page(s).

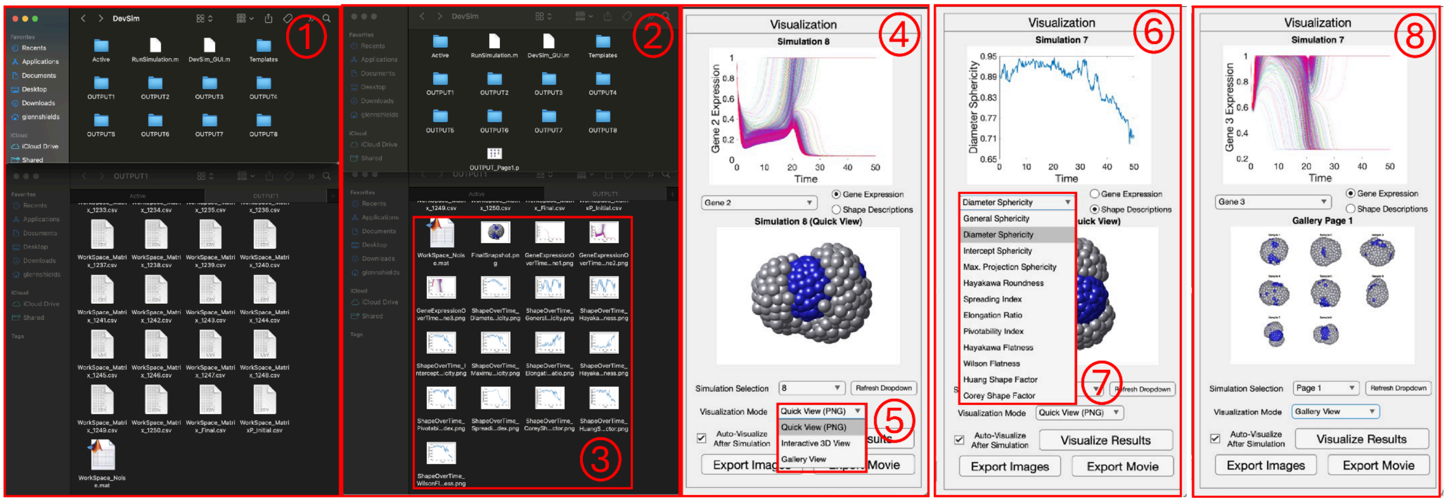

*Figure G8.* ① *DevSim file directory after a simulation has finished is shown with the OUTPUT folders stored in the directory (top).The OUTPUT1 folder is opened as an example to show the workspace matrices and how cell positions and gene values are stored over time (bottom).* ② *DevSim file directory after visualization has finished is shown with the OUTPUT folders from the simulation and also the Gallery View PNG file (top). The OUTPUT1 folder is opened as an example to show what is stored after visualization* ③ *after visualization the 3D cell renderings, Gene expression over time, and shape descriptor over time PNG's are generated and stored within their respective OUTPUT files* ④ *Visualization panel shown after DevSim simulation has been run and the results have been visualized. Simulation 8 quick view PNG 3D image is shown in axis below and Simulation 8 Gene 2 expression over time is shown in axis above.* ⑤ *The 3 different visualization modes are shown in a dropdown menu.* ⑥ *Visualization panel shown. Simulation 7 quick view PNG 3D image is shown in axis below and Simulation 7 Diameter Sphericity Shape Descriptor over time is shown in axis above.* ⑦ *The 12 different shape descriptions are shown in a dropdown menu.* ⑧ *Visualization panel shown with Simulation 7 Gene 3 expression over time shown in axis above, and Gallery View of all 8 simulations run shown in axis below.*

**(H) Export Movie (optional):** Use the "Export Movie" button to create an MP4 of a single simulation with user-defined time window, stride, and camera view.

1. "Click Export Movie button (Visualization panel), once pressed the Export Movie Settings popup appears, the user can then select the desired settings:

- "Output #": choose the simulation index (e.g., 1) corresponding to an OUTPUT* folder.
- "Start time" and "End time": any values between 0 and Tmax.
- "Frames per second (FPS)": sets the MP4 playback FPS (does not change sampling stride).

2. "Click OK once satisfied, the Resolution (frame stride) popup appears. Choose an option to control at what resolution the movie will be created with:

- "Full (1x)": uses every saved timepoint (longest render, longest movie).
- "Half (2x), Quarter (4x), Eighth (8x), Sixteenth (16x), Thirty-second (32x), and Sixty-fourth (64x)": uses every #x frames when rendering a movie (e.g. Half uses every other frame, Quarter uses every 4 frames, etc.) to shorten render/duration of movie.

- "Custom": enter any positive integer stride

3. "Click OK once satisfied, the Custom View popup appears enter Azimuth (-180 to 180) and Elevation (-90 to 90) at what camera angle the simulation will be rendered at

4. Press OK and a and a file viewer window will open, choose a name for the mp4, and choose a destination directory for the MP4 file to be saved to after the rendering.

5. Press OK and the movie will start rendering. The main progress bar tracks the progress of the movie rendering, and a frame-by-frame preview window shows each rendered frame (close this at any time to cancel the movie rendering).

6. Once finished, open the directory that was chosen to and play the MP4 file to view the movie.

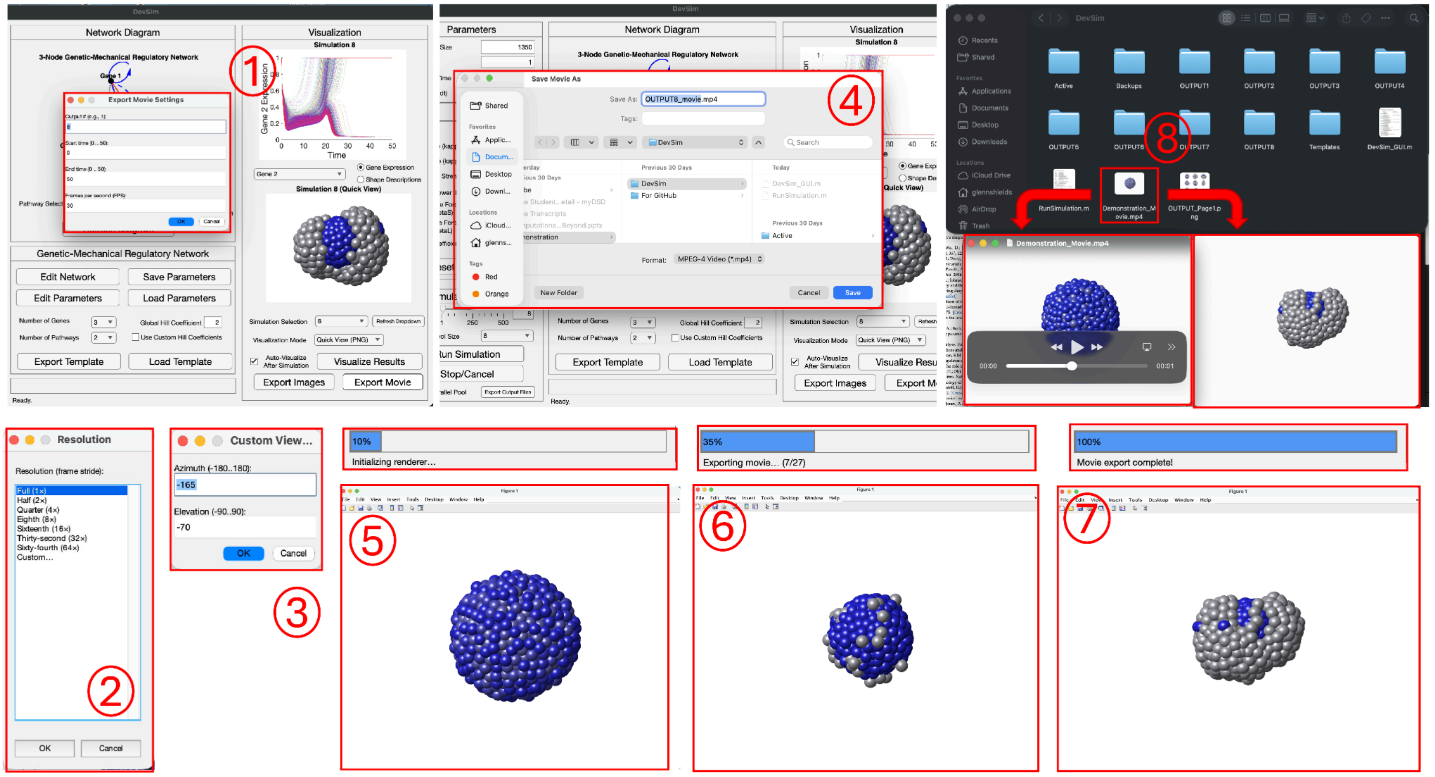


***Figure G9 here to show the export movie process.*** *① Export Movie Settings panel shown after user presses "Export Movie" button in the Visualization panel. Users will enter the Output number (corresponding to the sample they want to generate a movie for), the start and end times, and the Frames per second. ② Once the export movie settings have been entered, an additional panel will pop up called Resolution. Here the User will either pick "Full (1x), Half (2x), Quarter (4x), Eighth (8x), Sixteenth (16x), Thirty-second (32x), Sixty-fourth (64x) or "Custom" (integer) for their movie resolution. ③ Once the resolution has been picked, another pop up window called "Custom View" will prompt the user to enter an Azimuth and Elevation value. ④ Once the movie viewpoint has been chosen, a file directory viewer will pop up, prompting the user to choose a name for their .mp4 movie file, and to choose a directory to where that .mp4 movie file will be saved to. ⑤ Once the user is finished choosing all the settings for their movie, the movie will start to render. A frame-by-frame renderer will pop on so the user can view what it actually being rendered for their movie in real time which is shown via the bottom red box. The top red box shows the progress bar updates in DevSim as the movie is rendering. ⑥ Another timepoint choosing different rendering states in the movie that will be shown to the user (bottom red box). The top red box shows the progress bar as well as the status label showing useful information regarding the movie rendering. The (7/27) number shows that the movie has rendered 7 out of its 27 frames. ⑦ The final rendering*

*timepoint and progress bar are shown after the movie has finished rendering. ⑧The file directory that the .mp4 file was saved to is shown and the movie file is highlighted in red. Underneath are two instances of the same movie file open, the left one shows that it is indeed a playable .mp4 movie during an initial frame of the simulation; the right one shows a final frame of the movie.*

**(I) Import precomputed OUTPUT folders (Refresh Dropdown Option)**

*DevSim's* simulation and visualization are modular. If the user already have *DevSim*-compatible OUTPUT folders (e.g., OUTPUT1/ , OUTPUT2/ …); the user can load and visualize them without re-running simulations.

1. Prepare the *DevSim* folder: In DevSim/ directory, remove or move out any existing OUTPUT* folders to avoid conflicts.
2. Copy in results: Place the precomputed OUTPUT* folders directly under DevSim/ directory.
3. Open DevSim if not already running, and go to Visualization panel
4. Click the "Refresh" dropdown (Visualization panel). *DevSim* scans the DevSim/ directory, detects the imported OUTPUT* folders, and updates the "Simulation Selection" dropdown to match the count.
5. If PNG images are already present, choose a simulation in "Simulation Selection"; views render immediately.
6. If the imported results have not been visualized yet (no PNGs), set the "Total Simulations" box in Simulation Controls to the total number of OUTPUT folders that were imported into the DevSim/ directory. Then click "Visualize Results" to generate them and then follow instructions in section H from step 2 accordingly.

**(J) Save/load parameters and templates**

1. Save Parameters: writes current GUI parameter values to Active/UserParams.xlsx.
2. Load Parameters: reads Active/UserParams.xlsx into the GUI (useful when reopening *DevSim*)
3. Export Template: packages the duo (UserParams.xlsx, Network Settings.xlsx) into Templates/<name>/.
4. Load Template: copies a chosen template pack into Active/ (warning: if loading a template the current Active files will be replaced; the GUI will prompt and warn the user before this happens)

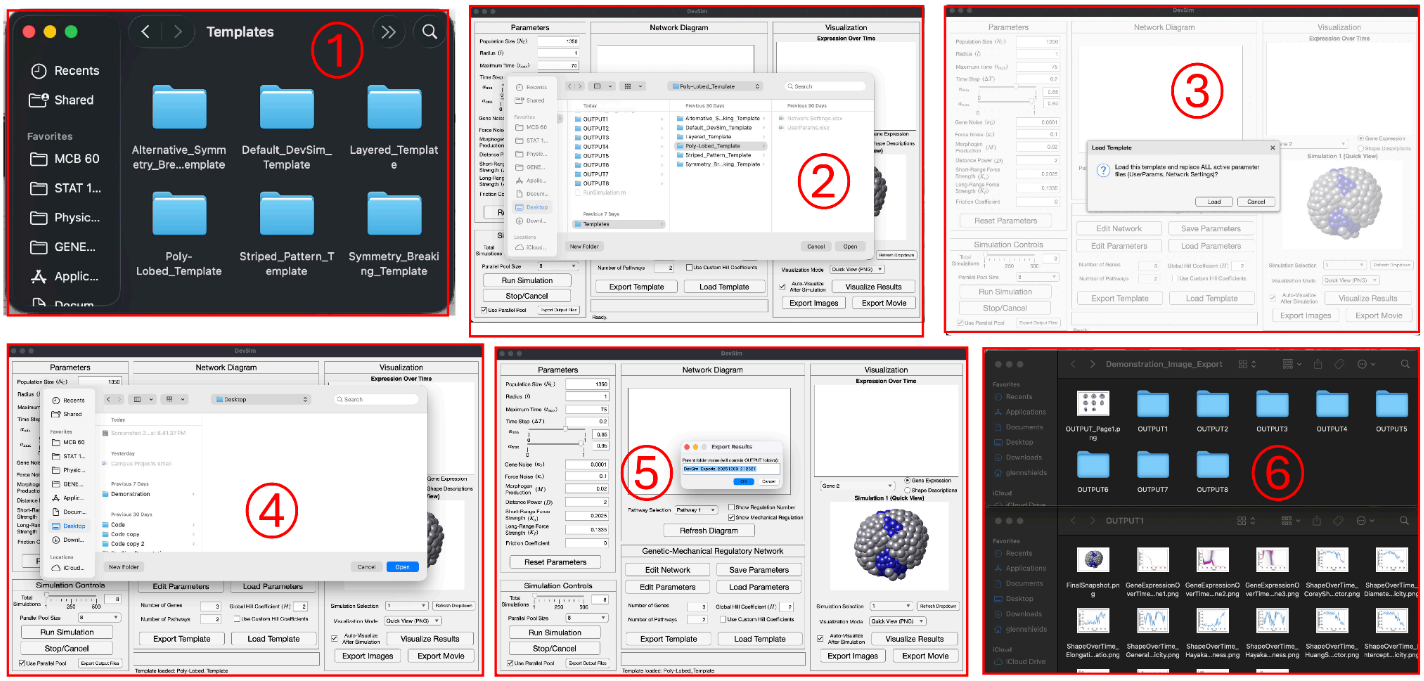


***Figure G10.*** *① Templates folder inside of DevSim directory is shown with the Default Template and a Demonstration Template created via pressing the Load Template button in the GMRN panel inside of DevSim GUI ② Shows the file directory users are shown after pressing the "Load Template" button; once a template is selected and the user presses "Open" in the file directory ③ DevSim will prompt the user to confirm that they want to load the selected template and as a result, replace all active parameter files where users can either go forward with the process by pressing Load or stop it by pressing Cancel. ④ Shows the file directory window users will be sent to once pressing Export Images Button in the Visualization panel inside of DevSim GUI. The users will select the location they want to save the parent folder holding all the .png files from DevSim's visualize results feature. ⑤ Once Users select a location for the parent folder they will be prompted to name the parent folder which will hold all the .png files in their respective OUTPUT folders ⑥ Parent folder created via Export Images button is shown for demonstration purposes called "Demonstration_Image_Export", within the directory there is the gallery view .png file, and the OUTPUT folders (above). An OUTPUT1 folder within the parent folder is opened and all of the .png files that DevSim created via the Visualize Results function are stored inside (below).*

**(K) Export Output Files:** use this feature to bundle an entire experiment into a single parent folder for archiving or sharing. Inside of the parent folder will include all the information for the experiment including the OUTPUT folders and all their contents, the gallery PNG's, and the Active folder containing the simulation settings that were used for that Simulation.

1. Press the Export Output Files button in the Simulation Controls panel. A file viewer window will open, then choose a destination directory for the parent folder to be saved to, press Open to use the selected option.

2. Enter a parent folder name for this export when prompted by the popup window. *DevSim* creates <destination>/<ParentName>/

3. Inside the parent folder, *DevSim* copies all OUTPUT* folders and their contents along with the Active folder containing the current simulation parameters, and the Gallery view PNG(s) saved at the top level.

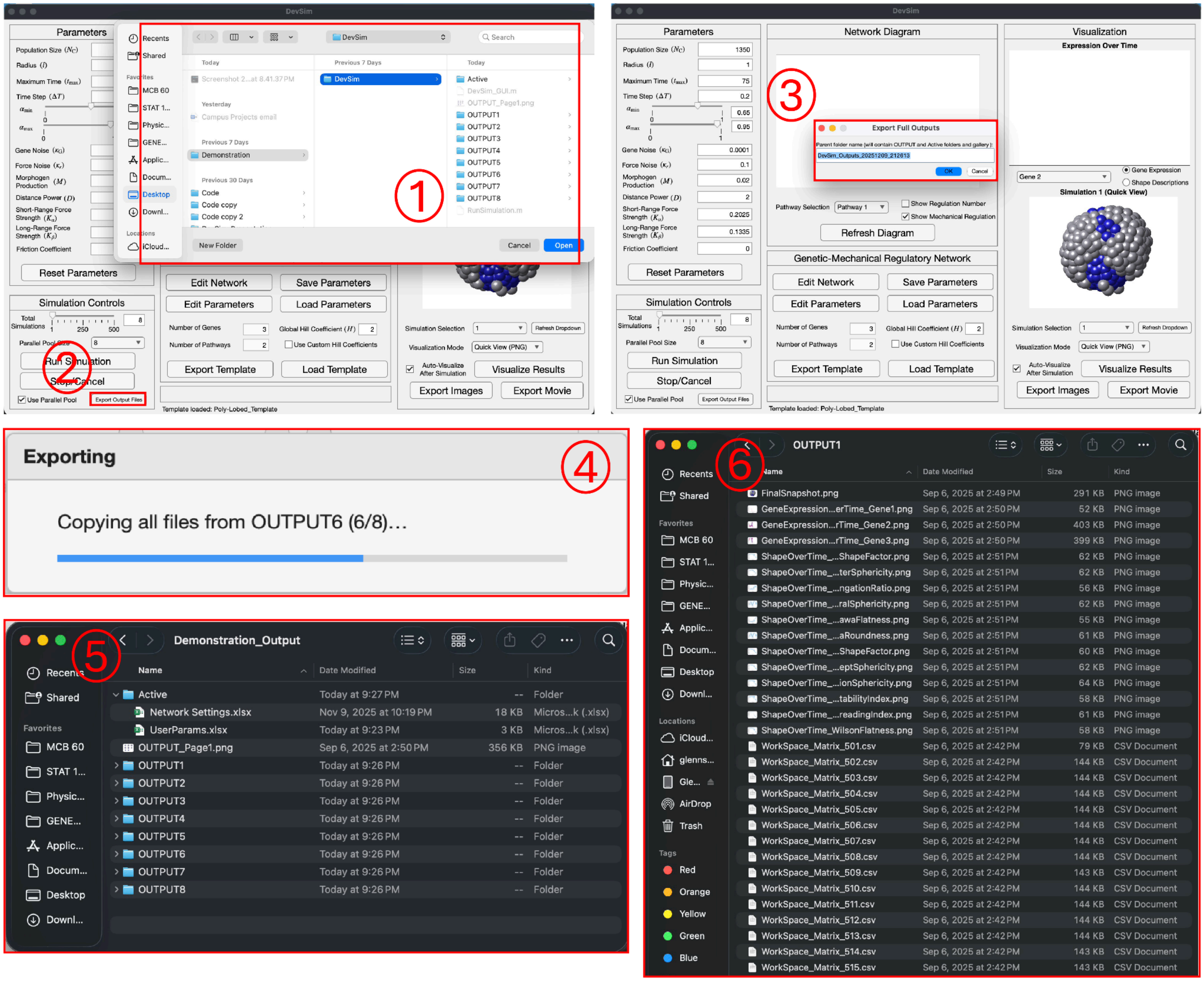

***Figure G11 Here showing export output files process.*** *① Shows the file directory Users are shown after pressing "Export Output Files Button". ② Export Output Files button located in the bottom of the Simulation Controls Panel that users will press to start the Export Output Files function. ③ Once users select a location for the parent folder, they will be prompted to name the parent folder which will hold all of the OUTPUT* folders, the Active folder, and the gallery view .png files. ④ Once the user selects the name for the parent folder, an exporting panel pop up showing the progress of copying all the OUTPUT folders into the Parent folder is shown. ⑤ Parent folder that was created for the demonstration and some of its contents are displayed for reference. ⑥ OUTPUT1 folder inside of the Parent folder that was created for the demonstration is displayed for reference.*

**Parameter Glossary (quick)**

Population: number of coarse-grained cells

Radius: Sets initial spacing scale and interaction distances

Maximum Time: total simulated time (in arbitrary units)

Time Step (dt): integration step size (smaller dt → more time points).

Alpha Max: Sets the upper bound for the typical balancing spacing in between cells

Alpha Min: Sets the lower bound for the typical balancing spacing in between cells

Gene Noise: Sets the gene noise amplitude

Force Noise: Sets the force noise amplitude

Morphogen Strength: Scales the external signal influence

Distance Power: Sets distance decay for external/morphogen signaling interactions

Short-Range Force Strength: Sets the short-range force amplitude

Long-Range Force Strength: Sets the long-range force amplitude

Friction Coefficient: Overdamped environment drag coefficient

**Troubleshooting (quick)**

- *DevSim* Folder is downloaded and the GUI is open but none of the Excel files can be accessed via "Edit Network" or "Edit Parameters" buttons, and features inside of the GUI don't work
  Make sure in the MATLAB window that the current folder is DevSim root, if not none of the features in *DevSim* will work properly. After setting the folder root, close the *DevSim* app and reopen it in MATLAB under the correct directory, then retry features.
- Diagram looks empty or wrong
  Click Refresh Diagram. Confirm "Number of Genes" and "Number of Pathways" in the GUI match the edited Excel ranges. Make sure Excel was saved before refreshing diagram inside the GUI
- Visualization panel is blank
  Press the "Refresh Dropdown" button, then select any simulation under the "Simulation Selection" dropdown. If there are no rendered images, most likely the visualization did not generate or save correctly. Press Visualize Results and wait for visualization to complete.

### METHOD DETAILS — Biophysical model

**High-level overview.** Each simulated cell carries an internal gene regulatory network that integrates (i) *intracellular* interactions among its own genes and (ii) *intercellular* signals emitted by neighboring cells and decaying with distance. The resulting gene-expression state is mapped onto two mechanical control parameters: a symmetric short-range interaction coefficient $\alpha_{m,n}$ governing contact forces, and a directed long-range interaction coefficient $\beta_{m,n}$ governing chemotactic-like forces. Through this coupling, gene regulation modulates cell–cell mechanical interactions, thereby shaping collective morphogenesis.

#### State

We consider a system of $N_\text{c}$ cells, each carrying an internal gene regulatory state comprising $N_\text{g}$ genes. For cell $m \in \{1, \dots, N_\text{c}\}$, the gene-expression state is represented by the vector

$$\mathbf{G}_m = \begin{bmatrix} G_{m,1} \\ \vdots \\ G_{m,N_\text{g}} \end{bmatrix}, \qquad G_{m,u} \in [0,1], \tag{1}$$

where $G_{m,u}$ denotes the normalized expression level of gene $u$ in cell $m$. Unless otherwise specified, gene-expression states are initialized to either 0 or 1.

Each cell has a spatial position $\mathbf{r}_m \in \mathbb{R}^3$. For any pair of cells $(m, n)$, we define the relative displacement vector, distance, and unit direction as

$$\mathbf{r}_m = [x_m \; y_m \; z_m], \qquad \mathbf{d}_{m,n} = \mathbf{r}_m - \mathbf{r}_n, \qquad d_{m,n} = \|\mathbf{d}_{m,n}\|, \qquad \hat{\mathbf{d}}_{m,n} = \frac{\mathbf{d}_{m,n}}{d_{m,n}} = [\hat{d}_{m,n,x} \; \hat{d}_{m,n,y} \; \hat{d}_{m,n,z}], \tag{2}$$

where $\|\cdot\|$ denotes the Euclidean norm.

Intracellular gene regulation is encoded by a third-order tensor $\mathbf{B}^{(\text{intra})} \in \{-1, 0, 1\}^{N_\text{g} \times N_\text{g} \times N_\text{p}}$. Each slice $p \in \{1, \dots, N_\text{p}\}$ corresponds to a distinct regulatory pathway. Within each slice, rows index target genes and columns index regulator genes, with entries $+1$, $-1$, and 0 denoting activation, inhibition, and no interaction, respectively. Intercellular gene regulation is defined analogously by $\mathbf{B}^{(\text{inter})} \in \{-1, 0, 1\}^{N_\text{g} \times N_\text{g} \times N_\text{p}}$.

Throughout, indices $m, n$ refer to interacting cells, while indices $u, v$ refer to interacting genes, corresponding to row and column indices of the associated regulatory tensors. Within each pathway $p$, intracellular regulation, intercellular regulation, short-range mechanical interactions, and long-range mechanical interactions are computed sequentially, as described below. The total update at each discrete time step is obtained by summing contributions across all $N_\text{p}$ pathways.

#### Intracellular regulation (Eq. 3)

The intracellular regulation block maps the current gene-expression state of cell $m$ to an effective regulatory response matrix $\mathbf{R}_m^{(\text{intra})} \in \mathbb{R}^{N_\text{g} \times N_\text{g}}$. This response encodes how each regulator gene influences each target gene within the same cell.

Regulatory interactions are specified by the tensor $\mathbf{B}^{(\text{intra})}$, together with pathway-specific thresholds $\mathbf{K}^{(\text{intra})}$ and sensitivities $\mathbf{H}^{(\text{intra})}$, both in $\mathbb{R}^{N_\text{g} \times N_\text{g}}$. For a given pathway, the combined matrix formulation of the intracellular response is

$$\boxed{\mathbf{R}_m^{(\text{intra})} = \mathbf{1} + \operatorname{sign}\big(\mathbf{B}^{(\text{intra})}\big) \circ \left[\frac{\mathbf{1}}{2} - \sigma_{\mathbf{K}^{(\text{intra})},\mathbf{H}^{(\text{intra})}}\big(\mathbf{G}'_m\big)\right] - \frac{|\mathbf{B}^{(\text{intra})}|}{2},} \tag{3}$$

where all operations are applied element-wise. Here, $\mathbf{G}_m$ is broadcast across $N_\text{g}$ columns to form $\mathbf{G}'_m \in \mathbb{R}^{N_\text{g} \times N_\text{g}}$, $\mathbf{1}$ denotes an all-ones matrix of compatible size, $\circ$ is the Hadamard product, and $\operatorname{sign}(\cdot)$ maps positive, negative, and zero entries to $+1$, $-1$, and 0, respectively.

The nonlinear response function $\sigma_{\mathbf{K},\mathbf{H}}$ is a Hill-type sigmoid defined element-wise by

$$\sigma_{\mathbf{K},\mathbf{H}}(\mathbf{G}) = \frac{\mathbf{K}^{\mathbf{H}}}{\mathbf{K}^{\mathbf{H}} + \mathbf{G}^{\mathbf{H}}}.$$

**Element-wise interpretation.** For an individual regulator–target gene pair, the matrix expression above is equivalent to the conditional formulation

$$R_m^{(\text{intra})} = \begin{cases} \dfrac{(G'_m)^{H^{(\text{intra})}}}{(K^{(\text{intra})})^{H^{(\text{intra})}} + (G'_m)^{H^{(\text{intra})}}}, & B^{(\text{intra})} = +1 \quad \text{(activation)}, \\[2ex] \dfrac{(K^{(\text{intra})})^{H^{(\text{intra})}}}{(K^{(\text{intra})})^{H^{(\text{intra})}} + (G'_m)^{H^{(\text{intra})}}}, & B^{(\text{intra})} = -1 \quad \text{(inhibition)}, \\[2ex] 1, & B^{(\text{intra})} = 0 \quad \text{(no regulation)}. \end{cases}$$

Thus, activating interactions yield a monotone increasing Hill response, inhibitory interactions yield the complementary decreasing response, and absent interactions contribute a constant factor of unity. Here we illustrate how the conditional (*element* representation) formulation can be easily derived from the combined (**matrix** representation) formulation: for any element, when $B = 1$, $R = 1 + (1) \cdot \left(\frac{1}{2} - \frac{K^H}{K^H+G^H}\right) - \frac{1}{2} = \frac{G^H}{K^H+G^H}$; when $B = -1$, $R = 1 + (-1) \cdot \left(\frac{1}{2} - \frac{K^H}{K^H+G^H}\right) - \frac{1}{2} = \frac{K^H}{K^H+G^H}$; when $B = 0$, $R = 1 + 0 \cdot \left(\frac{1}{2} - \frac{K^H}{K^H+G^H}\right) - \frac{0}{2} = 1$.

**Interpretation.** The matrix formulation provides a unified expression that simultaneously encodes activation, inhibition, and null regulation. The sign of $\mathbf{B}^{(\text{intra})}$ gates the direction of regulation, while the Hill nonlinearity maps gene-expression inputs into $[0, 1]$. The additive and subtractive constant terms ensure that, when $B^{(\text{intra})} = 0$, the response is identically 1, and when $B^{(\text{intra})} \neq 0$, the response spans the unit interval with opposite monotonicity for activation versus inhibition.

The resulting matrix $\mathbf{R}_m^{(\text{intra})}$ constitutes the intracellular regulatory output for cell $m$ and is used in subsequent update steps.

#### Intercellular regulation (Eq. 4)

The intercellular regulation block aggregates morphogen-like signals emitted by all other cells into an effective regulatory response matrix $\mathbf{R}_m^{(\text{inter})} \in \mathbb{R}^{N_\text{g} \times N_\text{g}}$ for cell $m$. Signals decay with intercellular distance according to a power-law kernel and are subsequently transformed by a Hill-type nonlinearity.

Intercellular regulatory interactions are specified by the tensor $\mathbf{B}^{(\text{inter})}$, together with thresholds $\mathbf{K}^{(\text{inter})}$ and sensitivities $\mathbf{H}^{(\text{inter})}$. Morphogen production strengths are encoded by $\mathbf{M}_\text{S} \in \mathbb{R}^{N_\text{g} \times 1}$, distance-decay exponents by $\mathbf{D}_\text{S} \in \mathbb{R}^{N_\text{g} \times 1}$, and $l$ denotes the characteristic cell radius. The combined matrix formulation of the intercellular response is

$$\boxed{\begin{aligned} \mathbf{R}_m^{(\text{inter})} &= \mathbf{1} + \operatorname{sign}\big(\mathbf{B}^{(\text{inter})}\big) \circ \left[\frac{\mathbf{1}}{2} - \sigma_{\mathbf{K}^{(\text{inter})},\mathbf{H}^{(\text{inter})}}\big(\mathbf{S}'_m\big)\right] - \frac{|\mathbf{B}^{(\text{inter})}|}{2}, \\ \mathbf{S}_m &= \sum_{n \neq m} \frac{\mathbf{M}_\text{S} \circ \mathbf{G}_n}{\left(\dfrac{d_{m,n}}{l}\right)^{\mathbf{D}_\text{S}}}. \end{aligned}} \tag{4}$$

Here $d_{m,n}$ is the intercellular distance defined in Eq. 2. The aggregated signal $\mathbf{S}_m$ is broadcast across $N_\text{g}$ columns to form $\mathbf{S}'_m \in \mathbb{R}^{N_\text{g} \times N_\text{g}}$. All operations are applied element-wise.

**Element-wise interpretation.** For an individual regulator–target gene pair, the matrix expression above is equivalent to the conditional formulation

$$R_m^{(\text{inter})} = \begin{cases} \dfrac{(S'_m)^{H^{(\text{inter})}}}{(K^{(\text{inter})})^{H^{(\text{inter})}} + (S'_m)^{H^{(\text{inter})}}}, & B^{(\text{inter})} = +1 \quad \text{(activation)}, \\[2ex] \dfrac{(K^{(\text{inter})})^{H^{(\text{inter})}}}{(K^{(\text{inter})})^{H^{(\text{inter})}} + (S'_m)^{H^{(\text{inter})}}}, & B^{(\text{inter})} = -1 \quad \text{(inhibition)}, \\[2ex] 1, & B^{(\text{inter})} = 0 \quad \text{(no regulation)}. \end{cases}$$

This gating logic is identical to that used for intracellular regulation in Eq. (3), with $\mathbf{S}_m$ replacing the intracellular gene-expression input.

**Interpretation.** The sign of $\mathbf{B}^{(\text{inter})}$ determines whether intercellular signaling induces activation, inhibition, or no regulation. The Hill nonlinearity maps aggregated morphogen signals into the unit interval, while the additive and subtractive constant terms ensure that absent regulatory interactions yield a constant response of unity. The power-law kernel weights contributions from neighboring cells as $(l/d_{m,n})^{\mathbf{D}_\text{S}}$, assigning greater influence to nearer cells.

The resulting matrix $\mathbf{R}_m^{(\text{inter})}$ constitutes the intercellular regulatory output for cell $m$ and is used in subsequent update steps.

#### Resultant gene regulation (Eq. 5)

Intracellular and intercellular regulatory responses are combined multiplicatively to yield the net regulatory effect acting on each gene in cell $m$. For each target gene $u$, the resultant regulatory term is defined as

$$\boxed{R_{m,u} = \prod_{v=1}^{N_\text{g}} \left(\mathbf{R}_m^{(\text{intra})} \circ \mathbf{R}_m^{(\text{inter})}\right)_{u,v}.} \tag{5}$$

Entries corresponding to absent regulatory edges are excluded from the product. If a gene has no regulators in a pathway, that pathway contributes zero to the production term. This product aggregates contributions from all regulator genes $v$, with unity corresponding to no net regulation. Finally, for each target gene $u$, the product of regulatory inputs $v$ within each parallel pathway $p$, as calculated above, are summed across all $N_p$ pathways to yield a joint regulatory term for gene $u$.

#### Gene-to-mechanics readouts (Eqs. 6–9)

Gene-expression states modulate mechanical interactions through two distinct channels: when $d < 2l$ short-range contact forces kicks in and when $d > 2l$ long-range directed forces in, where $l$ denotes the characteristic cell radius.

**Short-range force (Eqs. 6–7).** Short-range mechanical interactions are symmetric in the interacting cell pair $(m, n)$. Regulation of short-range forces is encoded by the vector $\mathbf{B}^{(\text{S})} \in \{2, -2, 1, -1, 0\}^{N_\text{g}\times 1}$, where the entries correspond to non-specific activation (+2), non-specific inhibition (−2), specific activation (+1), specific inhibition (−1), and no regulation (0). This encoding maps gene-expression states to an effective regulatory output $\mathbf{R}_{m,n}^{(\text{S})} \in \mathbb{R}^{N_\text{g}\times 1}$.

The combined matrix formulation and its equivalent element-wise representation are

$$
\boxed{
\begin{aligned}
\boldsymbol{R}^{(\mathrm{S})}_{m,n} &= \big[\mathbf{1} - \mathrm{sign}\big\{|\mathbf{B}^{(\mathrm{S})}|\big\}\big] \\
&\quad + \big[\mathbf{1} - \big|\mathrm{sign}\big\{|\mathbf{B}^{(\mathrm{S})}| - \mathbf{1}\big\}\big|\big] \left[\frac{\mathbf{1} - \mathrm{sign}\big\{\mathbf{B}^{(\mathrm{S})}\big\}}{2} (\mathbf{1} - \mathbf{G}_m) \circ (\mathbf{1} - \mathbf{G}_n) + \frac{\mathbf{1} + \mathrm{sign}\big\{\mathbf{B}^{(\mathrm{S})}\big\}}{2} \mathbf{G}_m \circ \mathbf{G}_n\right] \\
&\quad + \big[\mathbf{1} - \big|\mathrm{sign}\big\{|\mathbf{B}^{(\mathrm{S})}| - \mathbf{2}\big\}\big|\big] \left[\frac{\mathbf{1} - \mathrm{sign}\big\{\mathbf{B}^{(\mathrm{S})}\big\}}{2} \left(\mathbf{1} - \frac{\mathbf{G}_m + \mathbf{G}_n}{2}\right) + \frac{\mathbf{1} + \mathrm{sign}\big\{\mathbf{B}^{(\mathrm{S})}\big\}}{2} \left(\frac{\mathbf{G}_m + \mathbf{G}_n}{2}\right)\right], \\
\Leftrightarrow R^{(\mathrm{S})}_{m,n} &= \begin{cases}
\dfrac{G_m + G_n}{2}, & B^{(\mathrm{S})} = +2 \quad \text{(non-specific activation)}, \\
1 - \dfrac{G_m + G_n}{2}, & B^{(\mathrm{S})} = -2 \quad \text{(non-specific inhibition)}, \\
G_m\, G_n, & B^{(\mathrm{S})} = +1 \quad \text{(specific activation)}, \\
(1 - G_m)(1 - G_n), & B^{(\mathrm{S})} = -1 \quad \text{(specific inhibition)}, \\
1, & B^{(\mathrm{S})} = 0 \quad \text{(no regulation)}.
\end{cases}
\end{aligned}
}
\tag{6}
$$

The conditional formulation follows from the combined expression by the same gating logic used in Eq. (3), with regulation mode determining whether gene activities are combined jointly (via products) or independently (via averages).

The short-range mechanical readout and resulting force are then given by

$$
\boxed{\alpha_{m,n} = \alpha_{\max} - K_\alpha \prod_{u=1}^{N_\mathrm{g}} R^{(\mathrm{S})}_{m,n,u}, \qquad \mathbf{F}^{(\mathrm{S})}_{m,n} = \left(1 - \frac{d_{m,n}}{2\, l\, \alpha_{m,n}}\right) \hat{\mathbf{d}}_{m,n}, \quad d_{m,n} < 2l.}
\tag{7}
$$

Here $\alpha_{m,n} > 0$ sets the distance at which the contact force changes sign (with smaller values corresponding to stronger effective adhesion) and $\alpha_{\max} > 0$ sets the baseline (minimum) interaction strength, $K_\alpha$ controls the dynamic range of gene-dependent modulation, and $d_{m,n}$ and $\hat{\mathbf{d}}_{m,n}$ are defined in Eq. 2.

**Interpretation.** Gene-expression states are first mapped to an effective short-range interaction strength $\alpha_{m,n}$ through the discrete regulatory output $\mathbf{R}^{(\mathrm{S})}_{m,n}$. Depending on the regulatory mode, gene activities from the two cells are combined either conjunctively (via Hadamard products) or independently (via averaged sums). This interaction strength then determines the magnitude and sign of the short-range force: cells attract at intermediate separations and repel at very short distances. Because $\alpha_{m,n}$ depends multiplicatively on gene-expression states, the balance between attraction and repulsion is dynamically regulated by gene activity.

**Long-range force (Eqs. 8–9).** Long-range mechanical interactions are directed between an ordered cell pair $(m, n)$, where $m$ is the receiver and $n$ is the sender. Gene expression regulates long-range interactions through two components: (i) a signal-receiving capacity $\mathbf{R}^{(\mathrm{L})}_m$ that depends on the receiver cell, and (ii) a signal-sending kernel $\boldsymbol{\beta}_{m,n}$ that depends on the sender cell and spatial geometry. Their joint action determines the effective long-range force.

Regulation is encoded by the matrix $\mathbf{B}^{(\mathrm{L})} \in \{1, -1, 0\}^{N_\mathrm{g} \times N_\mathrm{g}}$, with entries corresponding to activation (+1), inhibition (−1), or no regulation (0). The receiver-side regulatory gate is defined by the combined (**matrix representation**) and equivalent conditional (*element-wise representation*) formulations:

$$
\begin{aligned}
&\mathbf{R}_m^{(\mathrm{L})\prime} = \mathbf{1} - \operatorname{sign}\left\{\mathbf{B}^{(\mathrm{L})}\right\} \circ \left(\frac{\mathbf{1}}{2} - \boldsymbol{G}_m'\right) - \frac{\left|\mathbf{B}^{(\mathrm{L})}\right|}{2} \\
&\Leftrightarrow \quad R_m^{(\mathrm{L})\prime} = \begin{cases} G_m', & B^{(\mathrm{L})} = 1 \quad \text{(activation)}, \\ 1 - G_m', & B^{(\mathrm{L})} = -1 \quad \text{(inhibition)}, \\ 1, & B^{(\mathrm{L})} = 0 \quad \text{(no regulation)}. \end{cases}
\end{aligned}
\tag{8}
$$

Here $\mathbf{G}_m$ is expanded to $N_\mathrm{g}$ columns and transposed to yield $\mathbf{G}_m' \in \mathbb{R}^{N_\mathrm{g}\times N_\mathrm{g}}$. The intermediate matrix $\mathbf{R}_m^{(\mathrm{L})\prime}$ is reduced by taking the product along each row. Entries equal to 1 (corresponding to absence of regulation) are mapped to 0, representing no contribution to long-range signaling. The resulting gene-resolved vector is then expanded across three spatial coordinates to form $\mathbf{R}_m^{(\mathrm{L})} \in \mathbb{R}^{N_\mathrm{g}\times 3}$. The three columns correspond to the Cartesian spatial directions $(x, y, z)$ and allow gene-dependent regulation of vector-valued forces. The conditional formulation follows from the combined expression by the same gating logic as in Eq. (3).

The signal-sending kernel and resulting long-range force are given by

$$
\boldsymbol{\beta}_{m,n,v} = K_\beta \frac{\mathbf{M}_\mathrm{F} \circ \mathbf{G}_n}{\left(\dfrac{d_{n,m}}{l}\right)^{\mathbf{D}_\mathrm{F}+1}} \hat{\mathbf{d}}_{n,m,v}, \qquad \mathbf{F}_{m,n,v}^{(\mathrm{L})} = \sum_{u=1}^{N_\mathrm{g}} \mathbf{R}_{m,n,u,v}^{(\mathrm{L})} \circ \boldsymbol{\beta}_{m,n,u,v}
\tag{9}
$$

Here $K_\beta$ controls the overall strength of long-range interactions, $\mathbf{M}_\mathrm{F} \in \mathbb{R}^{N_\mathrm{g}\times 1}$ specifies the force contributions of individual morphogens, and $d_{m,n}$ and $\hat{\mathbf{d}}_{m,n}$ are defined in Eq. (2). The additional index $v \in \{1, 2, 3\}$ denotes the three spatial coordinates $(x, y, z)$; thus both $\boldsymbol{\beta}_{m,n}$ and $\mathbf{R}_m^{(\mathrm{L})}$ are expanded across spatial dimensions so that gene-dependent regulation modulates vector-valued forces component-wise.

**Interpretation.** A sender cell $n$ exerts a directed attractive or repulsive force on receiver cell $m$ that decays with distance as $1/d_{m,n}^{\mathbf{D}_\mathrm{F}+1}$. The sender's gene-expression state determines the magnitude of the emitted signal, while the receiver's gene-expression state gates its responsiveness. The direction of the force is set by the unit displacement vector $\hat{\mathbf{d}}_{n,m}$. Together, these mechanisms enable gene-dependent, non-reciprocal, long-range mechanical interactions in three spatial dimensions.

#### Discrete-time updates (Eqs. 10–11)

Gene-expression states and cell positions are advanced in discrete time with step size $\Delta T$. Let $\mathbf{L}$ denote promoter leakage (basal production rate) and $\boldsymbol{\mu}$ the gene-specific degradation rates. At simulation time $T$, the gene state is updated as

$$
\mathbf{G}(T + \Delta T) = \mathbf{G}(T) + \boldsymbol{\xi}_G + \Delta T\Big[\mathbf{L} - \boldsymbol{\mu} \circ \mathbf{G}(T) + \mathbf{R}_m\Big]
\tag{10}
$$

Cell positions are updated according to overdamped dynamics,

$$
\mathbf{r}(T + \Delta T) = \mathbf{r}(T) + \frac{1}{\eta}\boldsymbol{\xi}_r + \frac{\Delta T}{\eta}\Big[\sum_{n\neq m} \mathbf{F}_{m,n}^{\mathrm{short}} + \sum_{n\neq m} \mathbf{F}_{m,n}^{\mathrm{long}}\Big]
\tag{11}
$$

Here $\eta$ denotes the effective viscous drag coefficient. The gene and positional noise terms, $\boldsymbol{\xi}_G$ and $\boldsymbol{\xi}_r$, are random matrices of appropriate dimensions, sampled independently from zero-mean Gaussian distributions.

**Simulation step.** At each time step, the following operations are performed:

(i) Compute intercellular signaling inputs $\mathbf{S}_m$ for all cells;

(ii) Evaluate intracellular and intercellular regulatory responses $\mathbf{R}_m^{(\mathrm{intra})}$ and $\mathbf{R}_m^{(\mathrm{inter})}$, and combine them to obtain $\mathbf{R}_m$;

(iii) Update gene-expression states according to Eq. (10), including leakage, degradation, regulatory input, and stochastic fluctuations;

(iv) Compute mechanical readouts $\alpha_{m,n}$ and $\boldsymbol{\beta}_{m,n}$ from gene states;

(v) Evaluate all pairwise short- and long-range forces and advance cell positions according to Eq. (11).

#### Practical update order

For clarity, a single simulation step proceeds as follows:

1. Compute the intercellular signaling inputs $\mathbf{S}_m$ for all cells using Eq. (4).
2. Evaluate intracellular and intercellular regulatory responses, $\mathbf{R}_m^{(\text{intra})}$ (Eq. 3) and $\mathbf{R}_m^{(\text{inter})}$ (Eq. 4), and combine them to obtain the resultant regulatory term $\mathbf{R}_m$ (Eq. 5).
3. Update the gene-expression state $\mathbf{G}$ using Eq. (10). Values are subsequently clipped to the interval $[0, 1]$.
4. For each ordered cell pair $(m, n)$, evaluate the short-range regulatory readout (Eq. 6) when $d_{m,n} < 2l$, and the long-range regulatory readout (Eq. 8) when $d_{m,n} \geq 2l$.
5. For each interacting pair, compute the corresponding mechanical coefficients $\alpha_{m,n}$ (Eq. 7) and $\boldsymbol{\beta}_{m,n}$ (Eq. 9).
6. Update cell positions using Eq. (11).

#### Numerical Example: A three-node gene regulatory network enabling robust symmetry breaking via differential control of short- and long-range forces (Fig. 6).

**Intracellular regulation (corresponding to Eq. 3): $\mathbf{B}^{(\text{intra})}$, $\mathbf{K}^{(\text{intra})}$, $\mathbf{H}^{(\text{intra})}$, and $\mathbf{R}^{(\text{intra})}$**

$$\mathbf{B}_1^{(\text{intra})} = \begin{bmatrix} 1 & 0 & 0 \\ 1 & 0 & -1 \\ 0 & -1 & 0 \end{bmatrix}, \quad \mathbf{B}_2^{(\text{intra})} = \begin{bmatrix} 0 & 0 & 0 \\ -1 & 1 & 0 \\ 0 & 0 & 0 \end{bmatrix},$$

$$\mathbf{K}_1^{(\text{intra})} = \begin{bmatrix} 0.7 & 0 & 0 \\ 0.1 & 0 & -0.5 \\ 0 & -0.5 & 0 \end{bmatrix}, \quad \mathbf{K}_2^{(\text{intra})} = \begin{bmatrix} 0 & 0 & 0 \\ -0.1 & 0.9 & 0 \\ 0 & 0 & 0 \end{bmatrix}, \quad \mathbf{H}_1^{(\text{intra})} = \begin{bmatrix} 2 & 2 & 2 \\ 2 & 2 & 2 \\ 2 & 2 & 2 \end{bmatrix}, \quad \mathbf{H}_2^{(\text{intra})} = \begin{bmatrix} 2 & 2 & 2 \\ 2 & 2 & 2 \\ 2 & 2 & 2 \end{bmatrix},$$

$$\Rightarrow \quad \mathbf{R}_1^{(\text{intra})} = \begin{bmatrix} \frac{G_1^2}{0.7^2 + G_1^2} & 1 & 1 \\ \frac{G_1^2}{0.1^2 + G_1^2} & 1 & \frac{0.5^2}{0.5^2 + G_3^2} \\ 1 & \frac{0.5^2}{0.5^2 + G_2^2} & 1 \end{bmatrix}, \quad \mathbf{R}_2^{(\text{intra})} = \begin{bmatrix} 1 & 1 & 1 \\ \frac{0.1^2}{0.1^2 + G_1^2} & \frac{G_2^2}{0.9^2 + G_2^2} & 1 \\ 1 & 1 & 1 \end{bmatrix}. \tag{12}$$

**Intercellular regulation (corresponding to Eq. 4): $\mathbf{B}^{(\text{inter})}$, $\mathbf{K}^{(\text{inter})}$, $\mathbf{H}^{(\text{inter})}$, $\mathbf{M}_\text{S}$, $\mathbf{D}_\text{S}$, $l$, and $\mathbf{R}^{(\text{inter})}$**

$$
\begin{aligned}
&\mathbf{B}_1^{(\text{inter})} = \begin{bmatrix} 0 & 0 & 0 \\ -1 & 0 & 0 \\ 0 & 0 & 0 \end{bmatrix}, \quad \mathbf{B}_2^{(\text{inter})} = \begin{bmatrix} 0 & 0 & 0 \\ 0 & 0 & 0 \\ 0 & 0 & 0 \end{bmatrix}, \\
&\mathbf{K}_1^{(\text{inter})} = \begin{bmatrix} 0 & 0 & 0 \\ -0.3 & 0 & 0 \\ 0 & 0 & 0 \end{bmatrix}, \quad \mathbf{K}_2^{(\text{inter})} = \begin{bmatrix} 0 & 0 & 0 \\ 0 & 0 & 0 \\ 0 & 0 & 0 \end{bmatrix}, \quad \mathbf{H}_1^{(\text{inter})} = \begin{bmatrix} 2 & 2 & 2 \\ 2 & 2 & 2 \\ 2 & 2 & 2 \end{bmatrix}, \quad \mathbf{H}_2^{(\text{inter})} = \begin{bmatrix} 2 & 2 & 2 \\ 2 & 2 & 2 \\ 2 & 2 & 2 \end{bmatrix}, \\
&\mathbf{M}_\text{S} = \begin{bmatrix} 0.02 \\ 0.02 \\ 0.02 \end{bmatrix}, \quad \mathbf{D}_\text{S} = \begin{bmatrix} 2 \\ 2 \\ 2 \end{bmatrix}, \quad l = 1, \\
\Rightarrow \quad &\mathbf{R}_1^{(\text{inter})} = \begin{bmatrix} 1 & 1 & 1 \\ \dfrac{0.3^2}{0.3^2 + S_1^2} & 1 & 1 \\ 1 & 1 & 1 \end{bmatrix}, \quad \mathbf{R}_2^{(\text{inter})} = \begin{bmatrix} 1 & 1 & 1 \\ 1 & 1 & 1 \\ 1 & 1 & 1 \end{bmatrix}.
\end{aligned}
\tag{13}
$$

where $\mathbf{S}$ is computed according to Eq. 4.

**Resultant genetic regulation (corresponding to Eq. 5): $\mathbf{R}$**

$$
\mathbf{R} = \begin{bmatrix} \dfrac{G_1^2}{0.7^2 + G_1^2} \\ \dfrac{G_1^2}{0.1^2 + G_1^2} \cdot \dfrac{0.5^2}{0.5^2 + G_3^2} \cdot \dfrac{0.3^2}{0.3^2 + S_1^2} + \dfrac{0.1^2}{0.1^2 + G_1^2} + \dfrac{G_2^2}{0.9^2 + G_2^2} \\ \dfrac{0.5^2}{0.5^2 + G_2^2} \end{bmatrix}. \tag{14}
$$

**Short-range force (corresponding to Eqs. 6–7): $\mathbf{B}^{(\text{S})}$, $\mathbf{R}^{(\text{S})}$, $\alpha_\text{max}$, $K_\alpha$, $\alpha$, $l$, and $\mathbf{F}^{(\text{S})}$**

$$
\mathbf{B}^{(\text{S})} = \begin{bmatrix} -1 \\ -2 \\ 0 \end{bmatrix}, \qquad \mathbf{R}_{m,n}^{(\text{S})} = \begin{bmatrix} (1 - G_{m,1}) \cdot (1 - G_{n,1}) \\ 1 - \dfrac{G_{m,2} + G_{n,2}}{2} \\ 1 \end{bmatrix}
$$

$$
\alpha_\text{max} = 0.95, \qquad K_\alpha = 0.2, \qquad \alpha_{m,n} = 0.95 - 0.2\,(1 - G_{m,1}) \cdot (1 - G_{n,1}) \ \cdot \ \left(1 - \frac{G_{m,2} + G_{n,2}}{2}\right), \quad l = 1, \tag{15}
$$

$$
\mathbf{F}_{m,n}^{(\text{S})} = \left[1 - \frac{d_{m,n}}{2 \cdot 1 \left(0.95 - 0.175\,(1 - G_{m,1}) \cdot (1 - G_{n,1}) \ \cdot \ \left(1 - \dfrac{G_{m,2} + G_{n,2}}{2}\right)\right)}\right] \hat{\mathbf{d}}_{m,n}, \qquad d_{m,n} < 2 \cdot 1.
$$

where $d_{m,n}$ and $\hat{\mathbf{d}}_{m,n}$ are computed according to Eq. 2.

**Long-range force (corresponding to Eqs. 8–9): $\mathbf{B}^{(\mathrm{L})}$, $\mathbf{R}^{(\mathrm{L})}$, $K_\beta$, $\mathbf{M}_\mathrm{F}$, $l$, $\mathbf{D}_\mathrm{F}$, $\boldsymbol{\beta}$, and $\mathbf{F}^{(\mathrm{L})}$**

$$\mathbf{B}^{(\mathrm{L})} = \begin{bmatrix} 0 & 0 & 0 \\ -1 & 1 & 0 \\ 0 & 0 & 0 \end{bmatrix}, \qquad \mathbf{R}^{(\mathrm{L})}_m = \begin{bmatrix} 0 & 0 & 0 \\ G_{m,2} \cdot \big(1 - G_{m,1}\big) & G_{m,2} \cdot \big(1 - G_{m,1}\big) & G_{m,2} \cdot \big(1 - G_{m,1}\big) \\ 0 & 0 & 0 \end{bmatrix},$$

$$K_\beta = -0.13, \qquad \mathbf{M}_\mathrm{F} = \begin{bmatrix} 1 \\ 1 \\ 1 \end{bmatrix}, \quad l = 1, \quad \mathbf{D}_\mathrm{F} = \begin{bmatrix} 2 \\ 2 \\ 2 \end{bmatrix},$$

$$\boldsymbol{\beta}_{m,n} = \begin{bmatrix} 0 & 0 & 0 \\ -0.13\, G_{n,2}\, \dfrac{\hat{d}_{m,n,x}}{d^3_{m,n}} & -0.13\, G_{n,2}\, \dfrac{\hat{d}_{m,n,y}}{d^3_{m,n}} & -0.13\, G_{n,2}\, \dfrac{\hat{d}_{m,n,z}}{d^3_{m,n}} \\ 0 & 0 & 0 \end{bmatrix} \tag{16}$$

$$\mathbf{F}^{(\mathrm{L})}_{m,n} = -0.13\, G_{n,2} \cdot G_{m,2} \cdot \big(1 - G_{m,1}\big)\, \frac{1}{d^3_{m,n}} \begin{bmatrix} \hat{d}_{m,n,x} & \hat{d}_{m,n,y} & \hat{d}_{m,n,z} \end{bmatrix}.$$

**Discrete-time updates (Eq. 10-11): G and r**

$$\begin{bmatrix} G_{m,1}(T+0.2) \\ G_{m,2}(T+0.2) \\ G_{m,3}(T+0.2) \end{bmatrix} = \begin{bmatrix} G_{m,1}(T) \\ G_{m,2}(T) \\ G_{m,3}(T) \end{bmatrix} + \begin{bmatrix} \xi_{G_{m,1}} \\ \xi_{G_{m,2}} \\ \xi_{G_{m,3}} \end{bmatrix} + 0.2 \left[ \begin{bmatrix} 0.02 \\ 0.0025 \\ 0.0025 \end{bmatrix} - \begin{bmatrix} 0.75 \\ 0.5 \\ 0.75 \end{bmatrix} \circ \begin{bmatrix} G_{m,1}(T) \\ G_{m,2}(T) \\ G_{m,3}(T) \end{bmatrix} \right]$$

$$+\, 0.2 \begin{bmatrix} \dfrac{G^2_{m,1}(T)}{0.7^2 + G^2_{m,1}(T)} \\ \dfrac{G^2_{m,1}(T)}{0.1^2 + G^2_{m,1}(T)} \cdot \dfrac{0.5^2}{0.5^2 + G^2_{m,3}(T)} \cdot \dfrac{0.3^2}{0.3^2 + S^2_{m,1}(T)} + \dfrac{0.1^2}{0.1^2 + G^2_{m,1}(T)} + \dfrac{G^2_{m,2}(T)}{0.9^2 + G^2_{m,2}(T)} \\ \dfrac{0.5^2}{0.5^2 + G^2_{m,2}(T)} \end{bmatrix}. \tag{17}$$

$$\mathbf{r}_m(T+0.2) = \mathbf{r}_m(T) + \begin{bmatrix} \xi_{m,x} \; \xi_{m,y} \; \xi_{m,z} \end{bmatrix} + 0.2 \Big[ \sum_{n \neq m} \mathbf{F}^{\mathrm{S}}_{m,n} + \sum_{n \neq m} \mathbf{F}^{\mathrm{L}}_{m,n} \Big]. \tag{18}$$

where $\xi_{G_{m,1}}$, $\xi_{G_{m,2}}$, and $\xi_{G_{m,3}}$ are random number, sampled from zero-mean Gaussian distributions with standard deviation of $0.0001\sqrt{0.2}$; $\xi_{S_{m,1}}$, $\xi_{S_{m,2}}$, and $\xi_{S_{m,3}}$ are random number, sampled from zero-mean Gaussian distributions with standard deviation $0.1\sqrt{0.2}$; $\mathbf{F}^{(\mathrm{S})}_{m,n}$ is computed according to Eq. 15 and $\mathbf{F}^{(\mathrm{L})}_{m,n}$ is computed according to Eq. 16.

**What the three-gene motifs are doing (intuitive map).**

$G_1$ acts as a *timer*: while $G_1$ remains high, the $(1-G_1)$ terms in Eqs. 15–16 prevent $\alpha$ and $\boldsymbol{\beta}$ from responding. As $G_1$ declines, mechanical responses switch on. $G_2$ and $G_3$ form a mutual-inhibition toggle (*bifurcation*) that partitions cells during the establishment stage; during the maintenance stage, this partition is stabilized (*lock-in*), after which $\alpha$ and $\boldsymbol{\beta}$ drive mechanical reorganization. Long-range pulls concentrate the peripheral cells while short-range bonds maintain cohesion where needed, thereby producing a single axis from homogeneous initial conditions, as shown in Fig. 6.
